# Extreme selective sweeps displaced archaic admixture across the human X chromosome around 50,000 years ago

**DOI:** 10.1101/503995

**Authors:** L. Skov, M.C. Macià, E. Lucotte, M.I.A. Cavassim, D. Castellano, T. Mailund, M.H. Schierup, K. Munch

## Abstract

The X chromosome in non-African populations has less diversity and less Neanderthal introgression than expected. We analyzed X chromosome diversity across the globe and discovered seventeen chromosomal regions, where haplotypes of several hundred kilobases have recently reached high frequencies in non-African populations only. The selective sweeps must have occurred more than 45,000 years ago because the ancient Ust’-Ishim male also carries its expected proportion of these haplotypes. Surprisingly, the swept haplotypes are entirely devoid of Neanderthal introgression, which implies that a population without Neanderthal admixture contributed the swept haplotypes. It also implies that the sweeps must have happened after the main interbreeding event with Neanderthals about 55,000 BP. These swept haplotypes may thus be the only genetic remnants of an earlier out-of-Africa event.

**One Sentence Summary:** After humans expanded out of Africa, the X chromosome experienced a burst of extreme natural selection that removed Neanderthal admixture.

## Main Text

The main wave of modern humans out of Africa around 60 ky ago was associated with several dramatic events, including admixture with Neanderthals (*1*), severe population bottlenecks, and exposure to new environments. Genetic studies have suggested that this most recent exodus may have displaced a previous wave out of Africa about 100-200 ky ago (*2*). Several fossils suggest that anatomically modern humans were indeed present in Israel about 180 ky ago (*3*) and in China 80-120 kyr ago (*4*), but see also (*5*). Despite its unique inheritance pattern, the X chromosome has received relatively little attention as a marker of migration processes (*6, 7*). The X chromosome is expected to be a hotspot for the accumulation of sex-antagonistic loci (*8*), as well as direct competition between the X and Y chromosomes for transmission in male meiosis (*9–11*). As a consequence, the X chromosome is uniquely involved with the evolution of reproductive barriers between species (*12–14*).

We have previously reported that very strong selective sweeps specifically targeted the X chromosome in each of the great ape species and that these sweeps together affected more than twenty percent of the chromosome (*15*). The sweeps in each great ape species coincide with large chromosomal regions with strong recurrent selective sweeps in the human-chimpanzee ancestor evident as a lack of incomplete lineage sorting between human, chimpanzee, and gorilla (*16, 17*). The swept regions also strongly overlap parts of the human X chromosome where Neanderthal introgression is extremely rare (*18*). Taken together, this suggests that genetic variants, which induce reproductive barriers by affecting male meiosis, are also subject to strong and recurrent episodes of positive selection.

Here we undertake a detailed investigation of X chromosome diversity in human populations using the Simons Genome Diversity Project of high coverage genomes sampled across the globe (*19*). We restricted the analysis to males where the X chromosome is haploid in order to analyze haplotypic variation without the need for computational phasing. We excluded males with missing data and males not showing the XY karyotype (*20*). We further removed African males with any evidence of recent European admixture (Materials and Methods). This left us with 162 males of which 140 are non-Africans (Table S1).

We first investigated the pairwise genetic distances of haplotypes in 100 kb windows for African and non-African individuals separately (Figure 1). The distributions of genetic distances are very different with a large proportion of highly similar X chromosome windows among non-African individuals (see also Figure S1). Whereas 17% of non-African 100kb haplotypes have a pairwise distance to other sampled individuals below 5e-5 (5 differences per 100 kb), this is true for only 2% of African 100kb haplotypes.

**Fig. 1.**
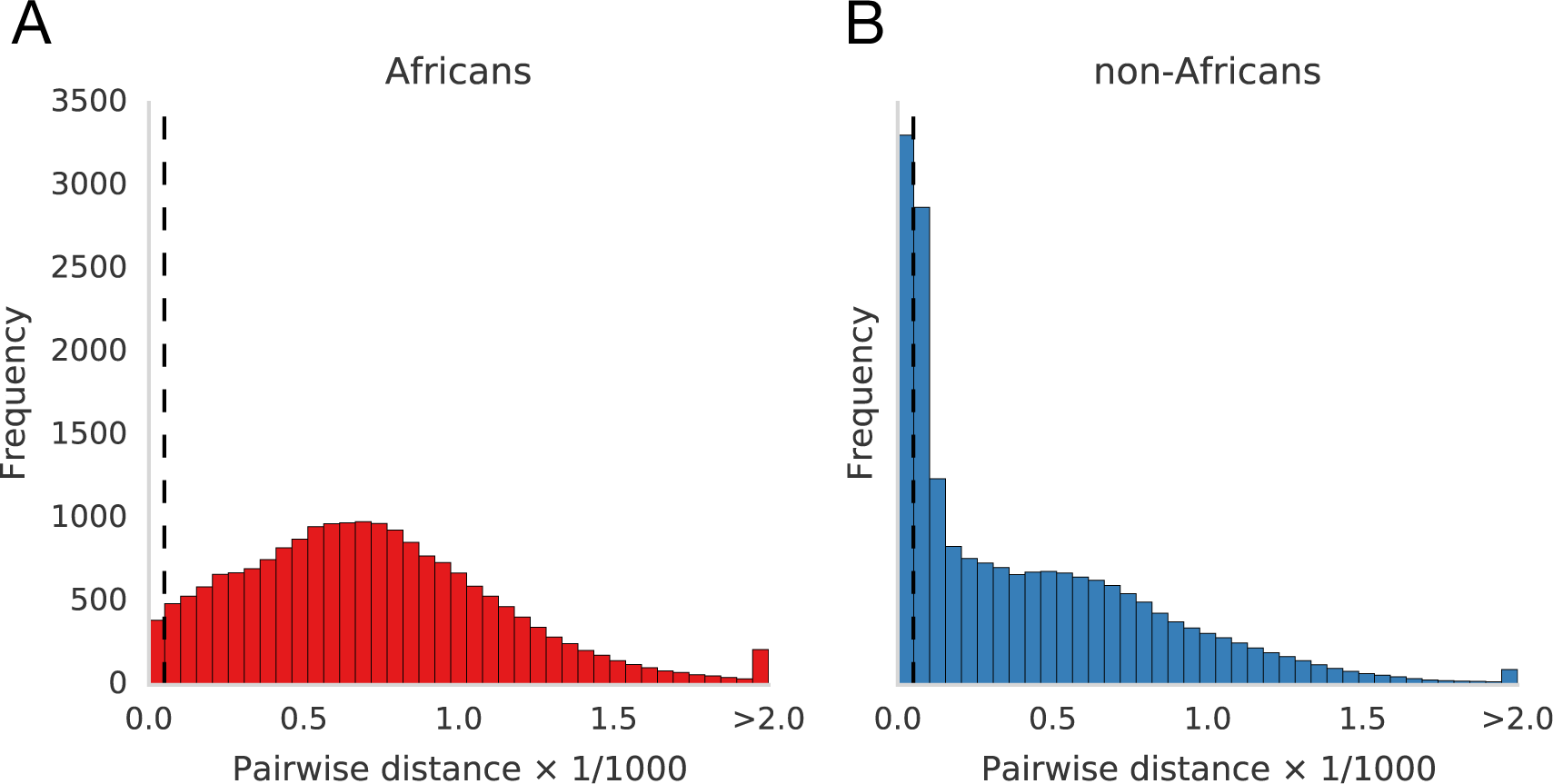
Excess of highly similar X haplotypes outside Africa. Distributions of pairwise sequence distances in non-overlapping 100kb windows. (A) Pairwise distances between African individuals. (B) Pairwise distances non-African individuals. Dashed line marks a pair-wise sequence distance of 5e-5.

We then wanted to investigate if the abundance of low pair-wise differences in 100 kb windows of non-Africans is the result of individual haplotypes appearing in high frequencies. To do this, we searched for long haplotypes with a very small genetic distance to many other haplotypes. Specifically, we identified haplotypes that are at least 500kb in length and which have a sequence distance smaller than 5e-5 to at least 25% of the individual haplotypes in the dataset (corresponding to at least 40 males). A haplotype that satisfies these criteria is referred to as an ECH (Extended Common Haplotype). We searched for ECHs using sliding windows of 500kb (step 100kb). The maximum sequence distance of 5e-5 was chosen because is corresponds to an expected coalescence time <60,000 years, post-dating the main exodus from Africa (*21*). We find that ECHs localize to distinct regions on the chromosome. Each of these regions is centered by a clear peak in the proportion of non-African haplotypes that we call as ECHs (Figure 2). For 17 of these regions, this peak includes the haplotypes of more than half of the non-African males. Even though ECHs are highly localized, they together cover 29% of the X chromosome (45.6 Mb).

**Fig. 2.**
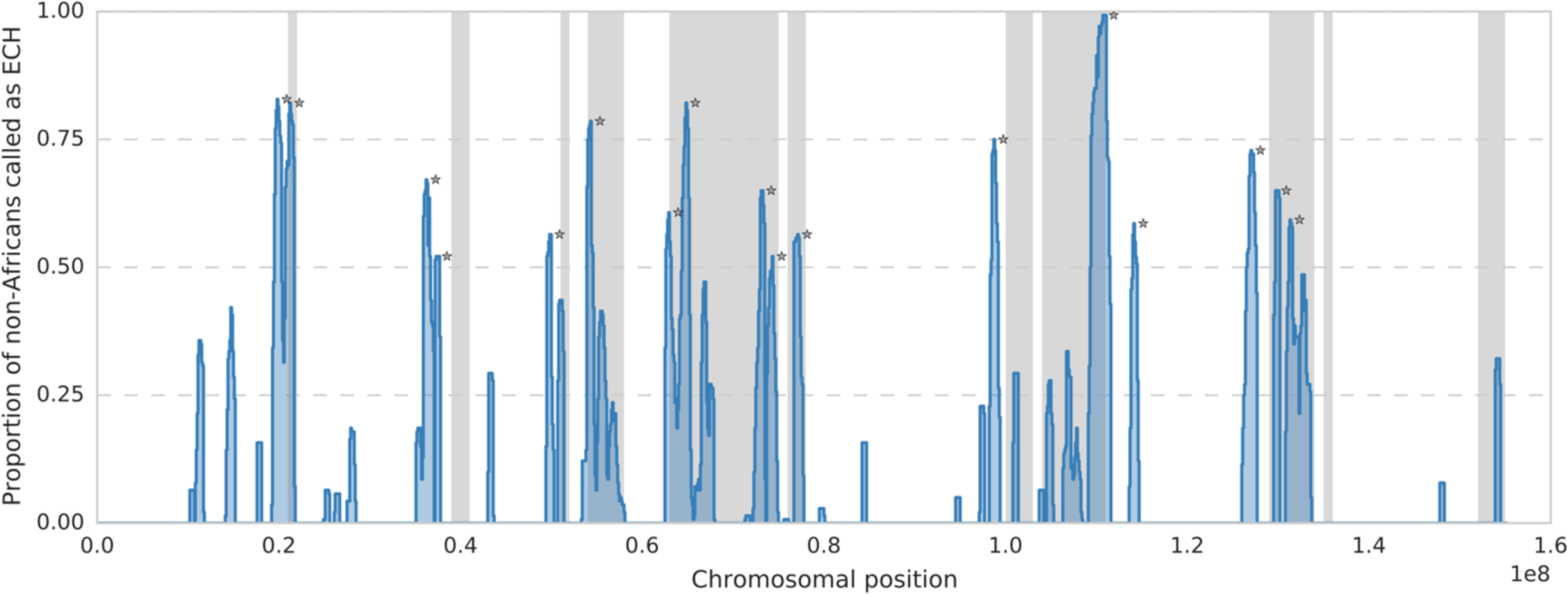
Proportion of non-African haplotypes called as ECH in each 100kb window across the complete X chromosome (blue). Stars mark the 17 most extreme regions with proportion of ECHs is larger than 50%. Regions depleted of incomplete lineage sorting between human, chimpanzee, and gorilla are shown in grey.

To determine if each of the swept regions represents one or several distinct haplotype clusters, we followed the approach of Tishkoff et al. (*22*) to directly visualize the haplotypes and their relationship (Materials and Methods). To this end, we identify a region around each peak in which at least 90% of the swept haplotypes are included (Materials and Methods and Table S2). Figure 3A provides an example of a 900kb region where non-Africans form a single clade with almost no diversity. This implies that a single haplotype has swept diversity in all non-Africans. For comparison, Figure 3C shows a typical region of the X chromosome without any evidence of swept haplotypes. Surprisingly, four of the 17 most extremely swept regions have two, clearly separated, low-diversity clades. One example of this is shown in Figure 3B, revealing two independent haplotype sweeps targeting overlapping regions of the X chromosome. Haplotype visualizations for all low-diversity regions are available as Data S1.

**Fig. 3.**
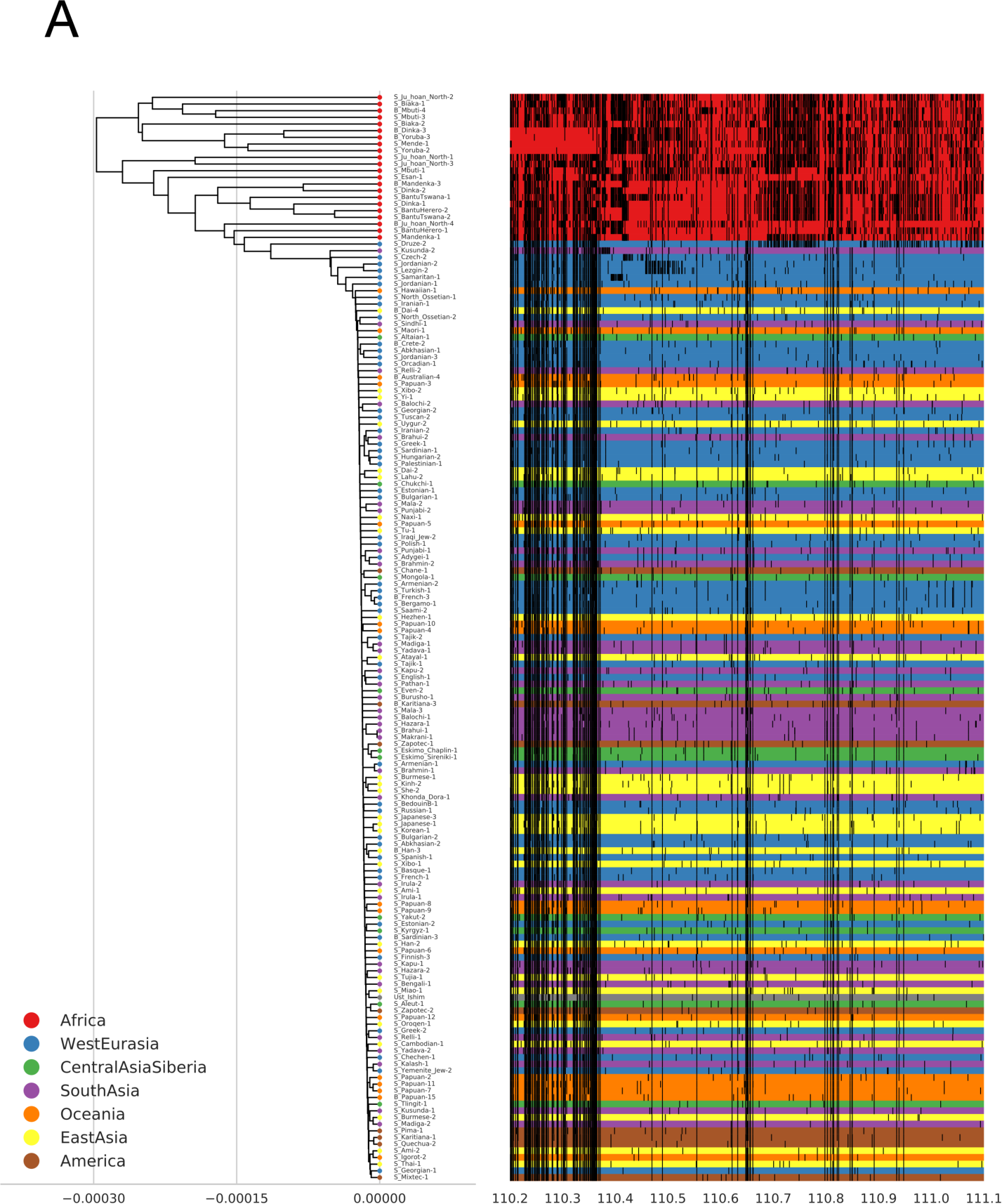
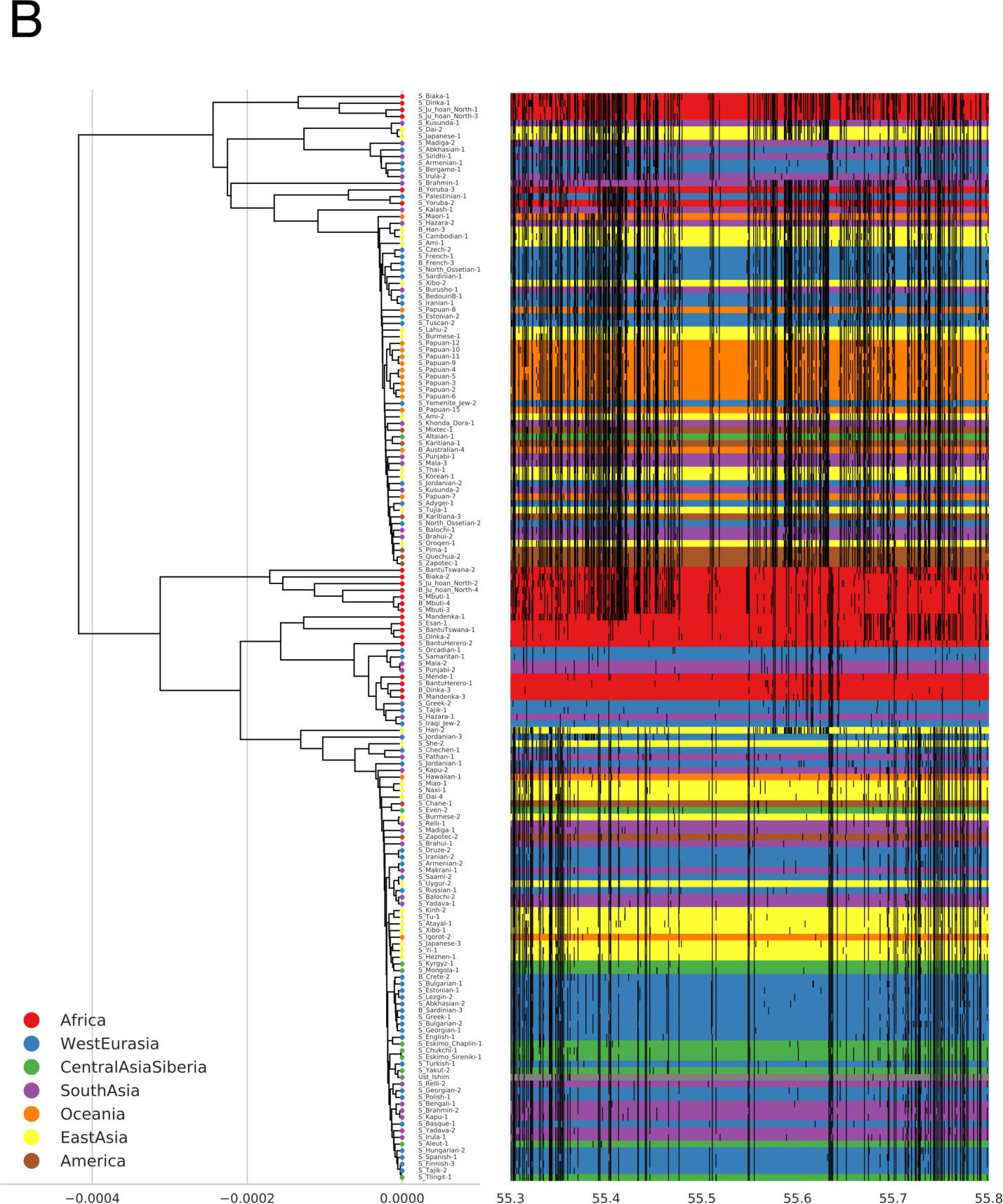
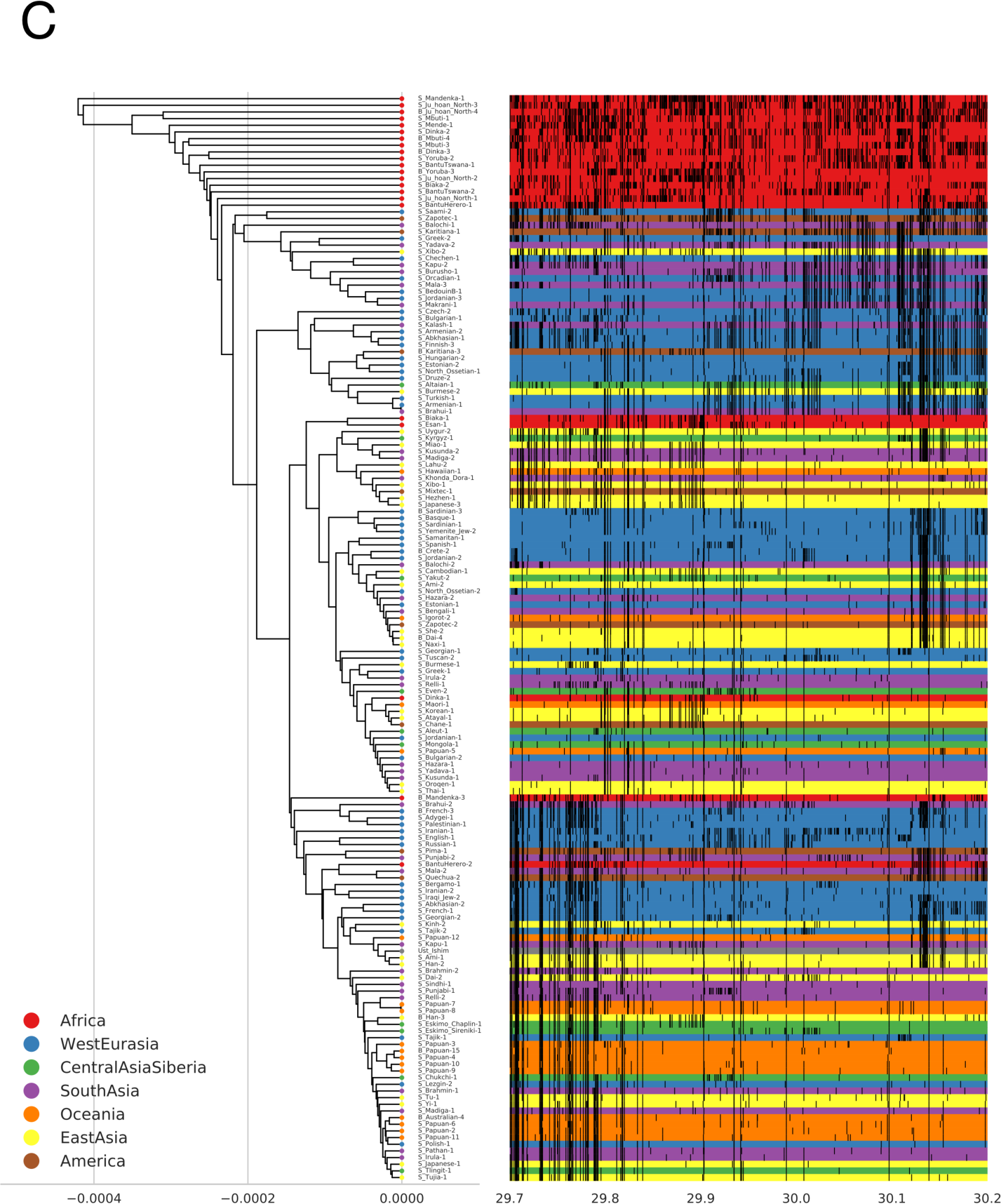
Examples of the relationship among haplotypes in identified regions: The left side of each figure is a UPGMA tree of the individual haplotypes shown as horizontal lines on the right. Haplotypes are color-coded according to geographical region. The ancient Ust’-Ishim individual is marked with grey. Vertical black bars on each haplotype represents non-reference SNPs. (A) 900kb region with all non-Africans forming a low diversity clade (coordinates: 110,200,000-111,100,000). (B) 500kb region with two clearly separated low diversity clades affecting most non-Africans (coordinates 55,300,000-55,800,000). (C) For comparison, a 500kb region where no ECHs were identified (coordinates: 29,700,000-30,200,000).

The haplotype plots further reveal that each sweep affected individuals from all of the major non-African geographical regions (Figures 3 and S3). In addition, each individual is part of many sweeps; on average, each non-African male is called as an ECH in 10.5 of the 17 most extreme regions surrounding peaks (5 and 95 percentiles: 6 and 14). These two observations together suggest that the sweeps must have occurred after the main exodus from Africa, but before the subsequent colonization of the world.

To narrow the period in which sweeps must have happened, we included analysis of the ancient Ust’-Ishim genome dated at 45,000 BP (*21*). The Neanderthal admixture in this ancient male is consistent with ancestry basal to present-day Europeans and East Asians. If the low divergence haplotypes did indeed rise to high frequency before humans spread across Eurasia, then we expect the Ust’-Ishim individual to be part of as many sweeps as living non-Africans. When we add the Ust’-Ishim male to the haplotype plots (Figure 3 and Materials and Methods), we find that the Ust’-Ishim falls inside a cluster of non-African ECHs in 11 of the 17 most extreme regions, very close to the non-African mean of 10.5.

It is highly unlikely that the large number of ECHs rose to high frequencies by genetic drift. If we conservatively assume a recombination rate of 1 cM/Mb and a human effective population size of one thousand during the out-of-Africa bottleneck, the probabilities that any single 500kb haplotype raises to frequencies higher than 0.4, 0.6, and 0.8 before it recombines are estimated to 6e-2, 4e-3, and 8e-05 (Materials and Methods). Background selection (linked selection on deleterious variants) may increase genetic drift in regions with many functional sites and low recombination rate. However, background selection is not expected to uniquely affect non-Africans and is not expected to create the large clades of highly similar haplotypes that is generated by a selective sweep. As an additional means to measure this signature of a selective sweep, we used ARGweaver (*23*) to estimate the time to the most recent common ancestor (TMRCA) for half of the sampled individuals divided by the TMRCA estimated for all sampled individuals. This relative TMRCA_half_ (*23*) is insensitive to background selection, but will be reduced where a selective sweep forces the recent common ancestry of many haplotypes. We find that the relative TMRCA_half_ of swept regions is sharply reduced, with a mean of only 0.06 compared to the chromosome-wide average of 0.15 (t-test p-value 2.7-20). In the 17 most extreme regions the mean is further reduced to only 0.04 (t-test p-value 9.7e-19). Below we will refer to the identified regions as selective sweeps.

The identified selective sweeps are as strong or even stronger than the most dramatic sweeps previously found in humans. Ten sweeps span between 500kb and 1.8Mb in more than 50% of non-Africans (Table S2). The strongest sweep span 900kb in 91% of non-Africans and affects 53% of non-Africans across a 1.8Mb region. For comparison, the strongest sweep previously reported surrounds the lactase gene and spans 800kb in 77% of European Americans (*24*). The selection coefficient on the genetic variant driving this sweep was estimated to 0.15 (*24*) suggesting even stronger selection for several of the X chromosome sweeps we have identified.

The swept regions we identify here may be recurrent targets of strong selection during human evolution. To investigate this possibility, we intersect our findings with our previously reported evidence of selective sweeps in the human-chimpanzee ancestor (*16*). We find a strong overlap between the sweeps reported here and regions swept during the 2-4 my that separated the human-chimpanzee and human-gorilla speciation events (*17, 25*) shown as grey regions in Figure 2 (Jaccard stat.: 0.17, p-value: <1e-5) (Materials and Methods). This suggests that the identified regions of the X chromosome are continually subjected to extreme positive selection.

Prompted by previous reports of wide regions on the X chromosome without archaic admixture, we applied a new method to call genomic segments of archaic human ancestry in each non-African male X chromosome ((*26*) and Materials and Methods). In line with previous findings (*27*), we find that the average proportion of archaic admixture on X chromosome is 0.9%. Restricting the analysis to regions where we have called ECHs (29.7% of the X chromosome) the archaic admixture proportion is only 0.4%, compared to 1.1% in regions where we do not call any ECHs (t-test p-value 9e-64). If sweeps and reduced admixture levels are causally related, then the swept haplotypes (ECHs) should contain less admixed sequence than haplotypes that are not swept in the same 100kb window. To investigate this, we identified all chromosomal windows where ECHs are called. For each such 100kb window we computed the separate mean archaic admixture proportion of the ECHs and the haplotypes not called as ECHs. By computing paired means for each window, we control for biases imposed by the distribution of ECHs across the chromosome. We find that swept haplotypes are almost entirely devoid of archaic admixture whereas non-swept haplotypes in the same regions have admixture proportions between 0.005 and 0.012 depending on the geographical region (Figure 4). In ECHs, the mean proportion of archaic admixture is only 1.2e-4 corresponding to a reduction by 98%. This is consistent with no archaic admixture since the inference of archaic admixture tracts is associated with a small false positive rate (*26*). The extreme depletion of archaic admixture in ECHs is independent of geographical region (Figure 4).

**Fig. 4.**
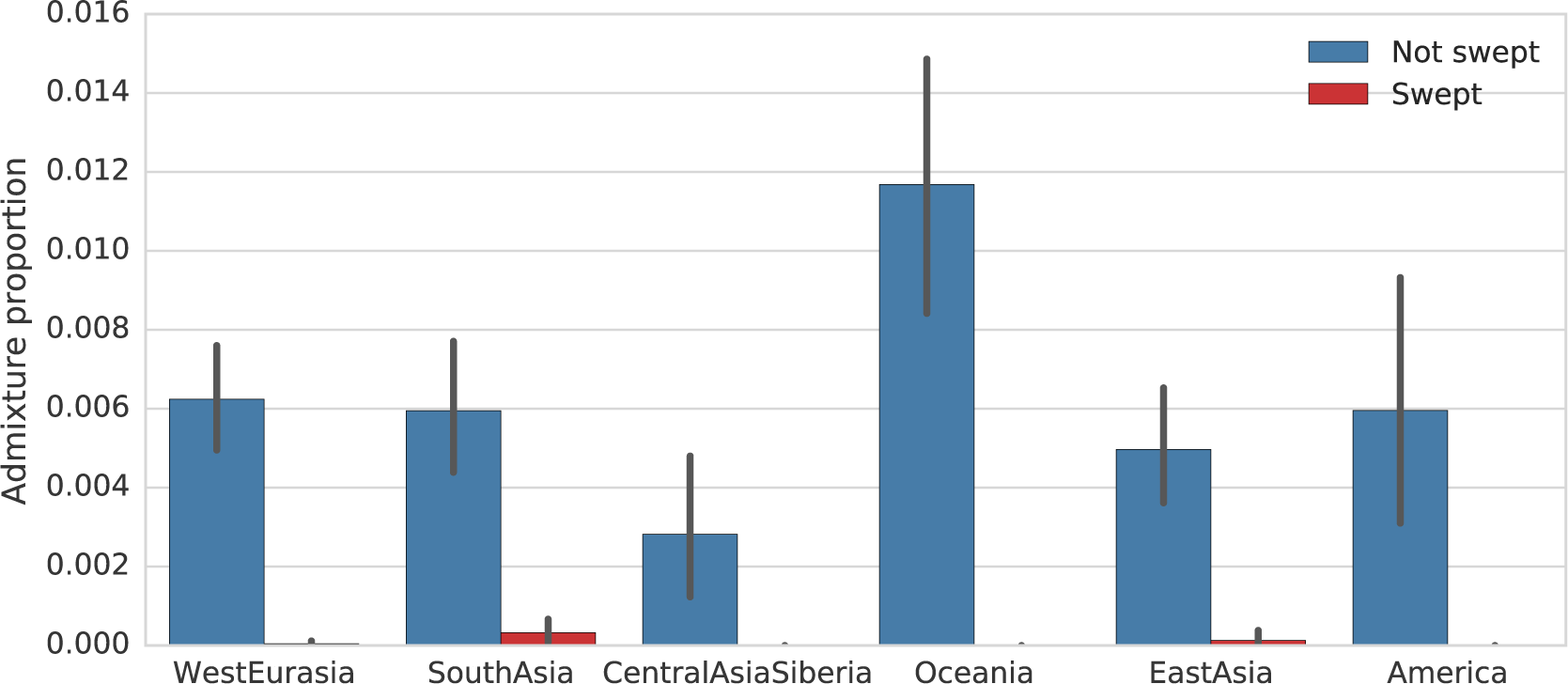
Admixture proportions in chromosomal regions of partial sweeps. Mean admixture proportions of in swept haplotypes (ECHs) (red) and in remaining haplotypes (blue) computed for each 100kb window.

The absence of archaic content suggests that the swept haplotypes were contributed from a population with little or no archaic admixture. This population could be an earlier wave out of Africa passing through the Middle East without meeting and interbreeding with Neanderthals. The fact that sweeps displaced Neanderthal admixture allow us to further narrow the period in which the sweeps must have occurred. From the length of Neanderthal admixture segments in the Ust’-Ishim male, the main Neanderthal admixture event has been estimated to have happened 7,000-13,000 years before he lived 45,000 BP (*21*). This implies that sweeps displacing admixture must have happened during the narrow time-span of roughly 10,000 years, which separated the Neanderthal admixture event (55,000 BP) and the Ust’-Ishim male (45,000 BP).

What triggered this remarkable burst of extreme selective sweeps is a mystery. However, the overlap to regions swept in great apes and the human-chimpanzee ancestor (*15, 16*) suggests that a general mechanism unique to the X chromosome must be responsible. The affected regions are not significantly enriched for any gene ontology, and protein-coding genes are not enriched for genes with elevated expression in testis (Materials and Methods). We have previously suggested the involvement of testis-expressed ampliconic genes, which are post-meiotically expressed in mouse testis (*28, 29*). One of the swept regions (shown in Figure S6) has an ampliconic region at its center, and ampliconic regions are significantly proximal to the swept regions (permutation test, 0.036), in many cases lining a sweep. We hypothesize that our observations are due to meiotic drive in the form of an inter-chromosomal conflict between the X and the Y chromosomes for transmission to the next generation. If an averagely even transmission in meiosis is maintained by a dynamic equilibrium of antagonizing drivers on X and Y, it is possible that the main bottlenecked out-of-Africa population was invaded by drivers retained in earlier out-of-Africa populations. If this hypothesis is true, the swept regions represent the only remaining haplotypes from such early populations not admixed with Neanderthals.

## Author contributions

L.S, M.C.M, M.I.C.A, D.C, T.M. and K.M conducted data analysis. E.L contributed data curation and annotation. K.M devised analysis. M.H.S and K.M. wrote the main text. L.S. and K.M. wrote supplementary text;

## Data and materials availability

All data is available in the main text or the supplementary materials. Code for the analysis is deposited on GitHub: https://github.com/kaspermunch/humanXsweeps.

## Supplementary Materials

Materials and Methods, Figures S1-S6, Tables S1-S2

References (*15–17, 19, 20, 22, 23, 26, 29–35*)

