## Supplementary material for "Extreme selective sweeps displaced archaic admixture across the human X chromosome around 50,000 years ago"

**This PDF file includes:**

Materials and Methods
Figs. S1 to S6
Tables S1 to S2
Captions for Data S1

**Other Supplementary Materials for this manuscript include the following:**

Data S1

### Materials and Methods

Dataset

We analyzed male X chromosomes from the fully public subset of genomes published as part of the Simons Genome Diversity Project (*19*). We exclude African individuals with evidence of gene flow from non-African populations: Mozabite, Saharawi show extensive non-African ancestry and Neanderthal admixture (*19*), Masai, Somali show a non-African component in STRUCTURE analysis (*19*), Luhya and close populations Luo and BantuKenya, show a non-African component in Admixture analysis (*30*). Following (*19*), we also exclude five samples (S_Finnish-1, S_Finnish-2, S_Mansi-1, S_Mansi-2, S_Palestinian-2) "based on missing X chromosome data in initial processing, for themselves or a second sample." We further exclude S_Lezgin-1 for which sex is not assigned, as well as S_Palestinian-2 and S_Naxi-2, which shows patterns of sequencing coverage not congruent with their assigned sex (*20*). This filtering leaves us with X chromosomes from 162 males among which 140 are non-Africans.

Pairwise sequence differences

For each pair of individuals, we computed the proportion of pairwise differences along the chromosome in non-overlapping windows of 100kb. Pairwise difference based on fewer than 50,000 called bases were discarded and considered missing data. Figure S1 shows the distributions of pairwise differences for each geographical region.

Identification of Extended Common Haplotypes (ECHs)

For each male, we identify haplotypes that are at least 500kb in length and which have a proportion of differences smaller than 5e-5 to at least 25% of the individuals in the dataset (at least 40 males). Note that we do not require that pairwise distances among such >40 males all fall below 5e-5. Imposing this additional requirement would limit our ability to identify large regions subject to sweeps and subsequent recombination. We search for these haplotypes using sliding windows of 500kb (step 100kb). Haplotypes in each individual identified by these criteria are referred to as ECHs. Figure S2 shows the chromosomal locations of ECHs in each individual. Figure S3 shows the chromosomal distribution of ECHs across individuals from each geographical region.

Operational definition of regions with high proportions of ECHs

In each region where ECHs are identified, we use a peak detection algorithm (*31*) to find the position where most individuals are part of an ECH and identify from that the consecutive 100kb windows that share this maximal number of individuals. We discard peaks with ECH frequencies smaller than 0.25. The coordinates of these peak regions are reported as "peak start" and "peak end" in table T2. We operationally define a region around each peak where the proportion of individuals called as ECH is at least 90% of the peak value. In Figure S4, each peak is shown as a red dot and the 90%-region around is shown as a red bar. We further define a set of wider regions where the proportion of individuals called as ECH rise above 75% of the peak value. These are shown as black bars in Figure S4. Table T2 lists chromosomal coordinates of both 90%-regions and 75%-regions along with the minimum proportion of non-African haplotypes called as ECH across each region and the proportion haplotypes that span each entire region.

Haplotype structure plots

Following Tishkoff et al. (*22*), we produced haplotype plots to visualize the relationship among haplotypes the 90%-regions around each peak (given in Table S2). The left side of each figure is a UPGMA tree based on jukes-cantor corrected sequence distances between male haplotypes. The tree clusters the individual haplotypes, which are shown as horizontal lines on the right, color-coded according to geographical region. Vertical black bars on each haplotype represent non-reference SNPs. Plots for all regions are provided as supplementary data (Data S1).

Incomplete lineage sorting between human, chimpanzee, and gorilla

In a previous analysis (*17*), we estimated the proportion of incomplete lineage sorting (ILS) between human, chimpanzee, and gorilla along the human genome. Here we analyzed autosomes and the X chromosome using the same protocol but published only estimates for autosomes. As in (*17*), the mean proportions of ILS are computed for non-overlapping 1Mb windows along the X chromosome. Windows, where the proportion of non-missing data falls below 30%, are not reported. The average proportion of ILS along the X chromosome is 13%. Following (*16*), mega-base windows are called as low-ILS windows. These regions, totaling 12% of the chromosome, are shown as grey regions in Figure 2 if the proportion of ILS falls below 5%.

Estimated TMRCA and relative $TMRCA_{half}$ along the chromosome

We ran ARGweaver (*23*) on 100kb alignments of the 162 male individuals. We used an effective population size of 10,000, a per generation recombination rate of 1.6e-8 (*32*), a per generation mutation rate of 1.8e-8, 20 discrete coalescence time points, and a maximal coalescence time of 200,000 generations. We ran two independent MCMC chains for 5,000 iterations sampling gene trees across the analyzed window for every ten steps. For each set of sampled trees, we computed the mean time to the most recent common ancestor (TMRCA) and the TMRCA for half of the sampled individuals ($TMRCA_{half}$). We analyzed the correlation of each summary statistic across iterations to ensure convergence. We discard the initial 1000 iterations as burn-in and compute the mean of each statistic across two chains. We compute the relative $TMRCA_{half}$ as $TMRCA_{half}$/TMRCA (Figure S5).

Estimating the probability that long haplotypes rise to high frequencies by genetic drift

To gauge the likelihood that our observations are the results of genetic drift, we estimate the probability (*P*) that any single haplotype in a population reaches a frequency *f* in less than *G* generations without recombining as

$$P = 1 - (1 - p)^{\frac{3}{4}N_e},$$

where *p* is the probability that this happens for a single haplotype in the population and $N_e$ is the effective autosomal population size. We simulated 10,000,000 allele frequency trajectories starting from an initial haplotype frequency of $1/N_e$ and computed *p* as

$$p = \frac{1}{n} \sum_{i=0}^{n} d_i e^{-rLg_i},$$

Where *n* is the number of sampled trajectories, *r* is the per base recombination rate, *L* is the length of the haplotype, $d_i$ is an indicator variable that is one if trajectory *i* reaches frequency *f* in less than *G* generations and zero otherwise, and $g_i$ is the number of generations at which trajectory *i* first reaches *f*. We simulate using *G*=344 generations (10,000 years assuming 29 years per generation). We conservatively set the autosomal $N_e$ to a constant size of only 1,000 and the recombination rate to only one mM/Mb (~2/3 of X chromosome average). The probability *P* for a range of frequencies *f* and a haplotype length of 500kb is given in the table below. When no sampled trajectories reach *f*, *p* must be smaller than *exp(-rLG)/n* corresponding to a *P* of <2.7e-6.

| **Frequency (f)** | **Estimated probability (P)** |
|---|---|
| 0.1 | 8.0e-01 |
| 0.2 | 4.2e-01 |
| 0.3 | 1.7e-01 |
| 0.4 | 5.9e-02 |
| 0.5 | 1.9e-02 |
| 0.6 | 4.4e-03 |
| 0.7 | 7.8e-04 |
| 0.8 | 8.1e-05 |
| 0.9 | 2.7e-05 |
| 1.0 | <2.7e-06 |

Extended Common Haplotypes (ECH) shared with ancient Ust ishim male

The sequence of the Ust ishim male was available as part of the Simons Genome Diversity data set. We computed pairwise sequence distances 100kb windows between the Ust ishim male and all present-day males. Distances, where less than 20% of bases are called in both sequences, were discarded. The distribution of these sequence distances shows the same characteristic enrichment of short pair-wise distances as present-day non-Africans.

To quantify the extent to which the Ust'-Ishim male shares an ECH called in sampled non-Africans, we measure the sequence distance between the Ust'-Ishim male and each extant sample in the 90%-regions identified around each of the 17 most extremely swept regions. For five of the 17 regions (see table below), we find that at least 40 individuals (25%) have a distance to Ust ishim that is smaller than 3.125e-5. This cut-off corresponds to the one used in the remaining analysis (5e-5) but accounts for the sampling time of the Ust ishim male.

| **Start** | **End** |
|---|---|
| 21100000 | 21600000 |
| 36000000 | 36400000 |
| 98500000 | 98900000 |
| 114000000 | 114300000 |
| 126800000 | 127400000 |

This stringent cut-off allows only three base differences in each 100kb and any spurious differences in this ancient genome thus will raise the proportion of differences above the cut-off. To accommodate this, we instead adopt the alternative criterion, that the Ust'-Ishim haplotype falls inside a UPGMA cluster of non-African haplotypes called as ECH (Data S1). By this criterion the Ust'-Ishim shares an ECH in additional six of the 17 regions (see table below).

| **Start** | **End** |
|---|---|
| 54000000 | 54400000 |
| 64700000 | 65100000 |
| 73000000 | 73500000 |
| 74000000 | 74400000 |

| | |
|---|---|
| 110200000 | 111100000 |
| 129700000 | 130200000 |

Inference of admixture segments

We use a newly developed hidden Markov model to infer archaic segments (*26*). All scripts for using the method are online at the GitHub repository: https://github.com/LauritsSkov/Introgression-detection.

The method finds archaic genomic segments in non-Africans by identifying segments with a strong enrichment of single nucleotide variants that are not seen in an outgroup population (in this case Sub-Saharan populations). Non-African variants not present in the outgroup, show a ten-fold enrichment in archaic segments because these variants have been accumulated since the common ancestor of archaic and modern humans. We use an outgroup consisting of individuals from two datasets. We use all Sub-Saharan Africans (populations: YRI, MSL, ESN) from the 1000 Genomes Project that do not have a component shared with European population ((*30*) figure 2a) and all Sub-Saharan African populations from SGDP except Masai, Somali, Sharawi and Mozabite ((*19*) see supporting figure 8.1), which show signs of out-of-Africa admixture. For all data, we removed sites that fell within repeat-masked (*33*) regions (downloaded from the UCSC genome browser) as well as sites that were not in the strict callability mask for the 1000 Genomes Project:

(ftp://ftp.1000genomes.ebi.ac.uk/vol1/ftp/release/20130502/supporting/accessible_genome_masks/StrictMask/).

Since the method is sensitive to variation in mutation rate, we calculate the background mutation rate using the variant density of all variants from populations YRI, LWK, GWD, MSL and ESN in windows of 100 Kb divided by the mean variant density of the whole genome.

The hidden Markov model was trained using the whole genome and not only the X chromosome. The reason for this is that there are not enough archaic segments to accurately infer transition and emission parameters just using the X. To accommodate the smaller effective population size of the X chromosome, we scaled the emission values using the following approach: If males contribute 3/4 of the mutations, but the X chromosome only spends a third of its time in males, then the X chromosome mutation rate is 5/6=0.8333 of the autosomal one, i.e., if the autosomal emission of state 2 is 0.38, it should be 0.3166 for X.

Archaic segments are then called as consecutive regions where the posterior probability of the archaic state in the HMM is at least 0.8. Segments that are interleaved by more than 25Mb of missing data are split in two. 1kb windows with less than 200 bases called are excluded. We then compute the mean proportion of archaic sequence in each non-overlapping 100kb window.

In line with previous findings (*27*), the mean proportion of admixture segments identified in each X chromosome is 0.9%, (WestEurasia: 0.8%, SouthAsia: 0.9%, CentralAsiaSiberia: 0.6%, Oceania: 1.5%, EastAsia: 0.9%, America: 0.7%).

Gene ontology and tissue enrichment

Gene ontology enrichment of genes overlapping 90%-regions was done using GOrilla (*34*) but did not reveal any significant enrichment. Two sets of testis-expressed genes were obtained from the Human Protein Atlas: One set of 2237 genes showing elevated expression in the testis compared to other tissues, and one set of 1079 enriched with at least five-fold higher mRNA levels in testis compared to all other tissues. Neither set showed a significant overlap to 90%-regions using a Fisher's exact test.

Relationship between sweeps and ampliconic regions

We have previously reported that testis-expressed ampliconic regions (*29*) significantly overlap selective sweeps on the X chromosomes of great apes and of the human-chimpanzee ancestor (*15, 16*). Here we find one target region (shown in Figure 3B) that has an ampliconic region at its center. However, inflated nucleotide diversity due to over-collapsing of reads mapped to the reference genome may limit our ability to identify sweeps overlapping ampliconic regions. Using a permutation test (*35*) we find that the remaining ampliconic regions show a significantly reduced distance to the 90%-regions defined in table S2 (p-value: 0.036).

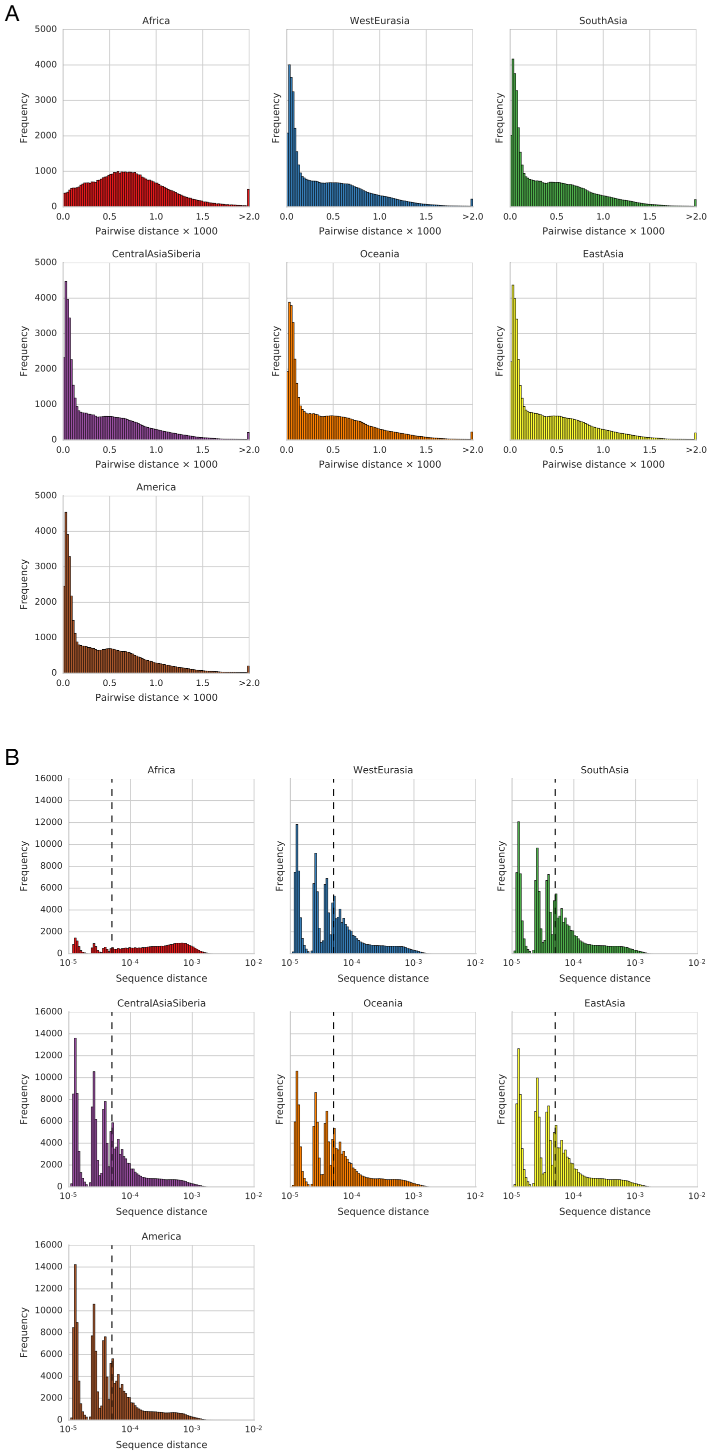


**Fig. S1.**

(A) Distributions of pairwise sequence distances in non-overlapping 100kb windows. (A) Pairwise distances between individuals from each major geographical region. (B) Same as A but with distances shown on a log scale. The dashed line represents cut-off used to call low-divergence haplotypes. Individual peaks represent the contributions of single differences.

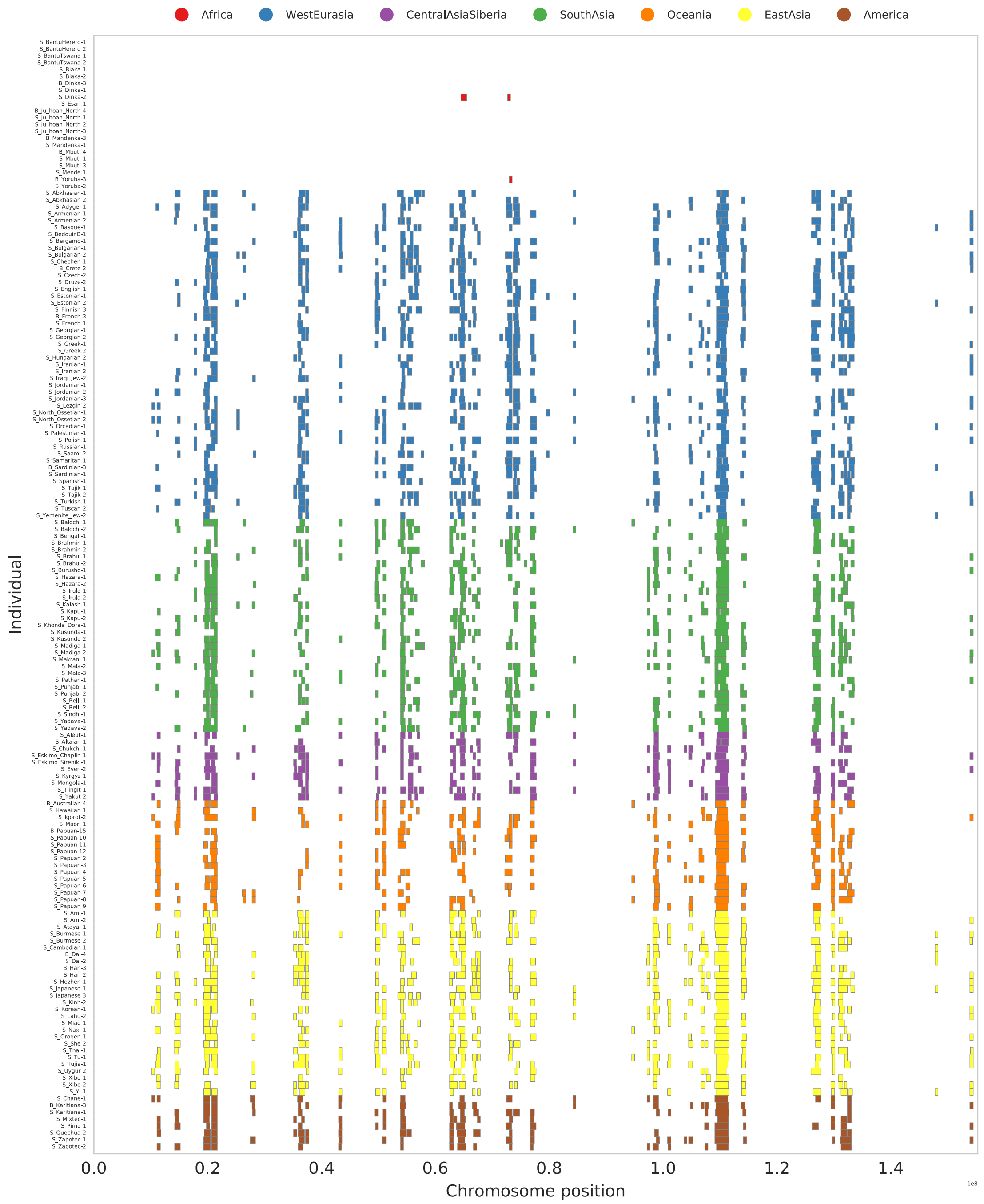


**Fig. S2.**

Chromosomal location of individual haplotypes that are called as ECHs.

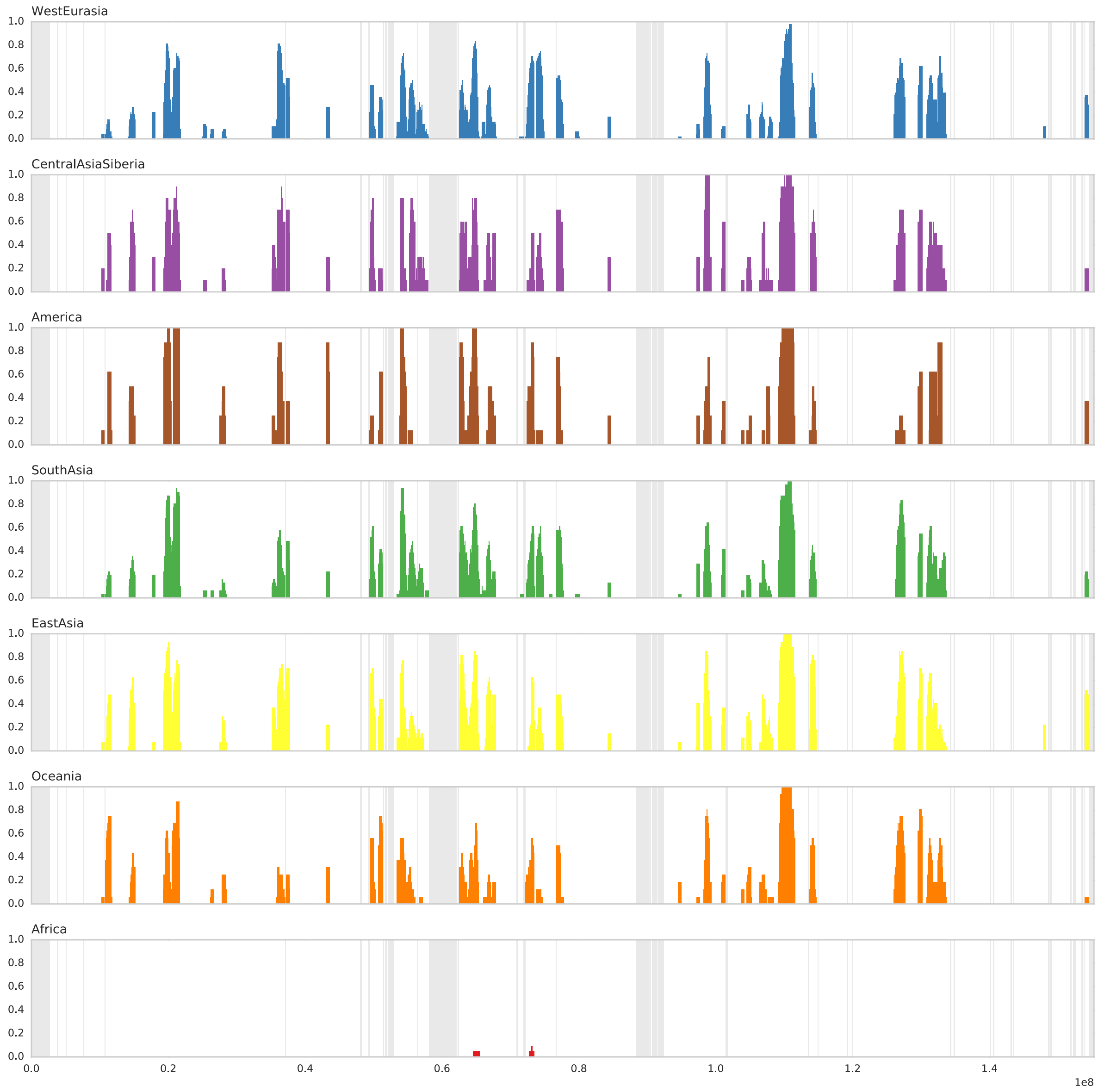


**Fig. S3.**

Proportion of individuals from each geographical region called as ECH in each 100kb window. Shaded regions represent regions with missing data.

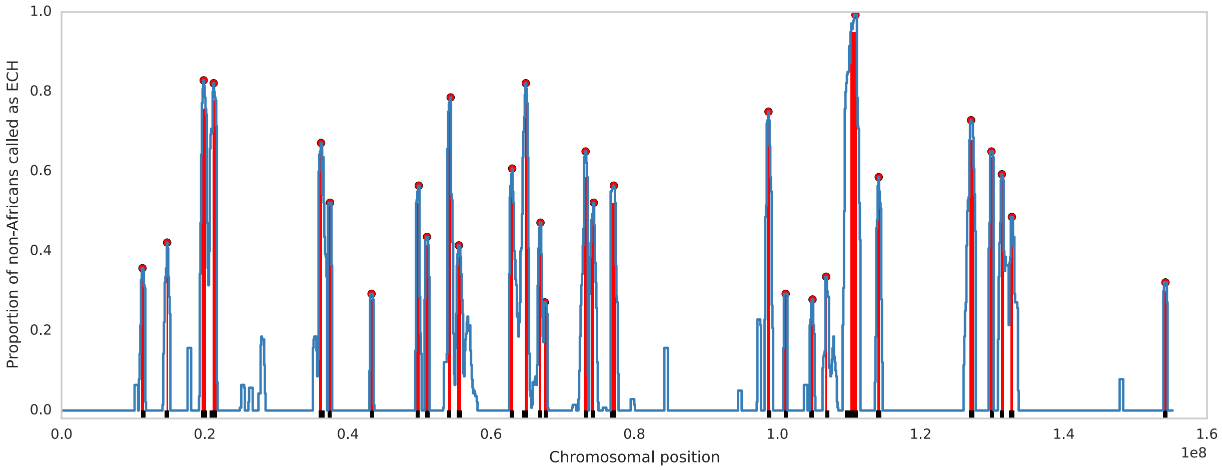


**Fig. S4.**

Operational definition of regions with high proportions of ECHs. Blue lines show the proportion of non-African individuals called as ECH in each 100kb window. Red dots show called peaks. Red bars show the extent of defined 90%-regions. Black bars show the extent of defined 75%-regions.

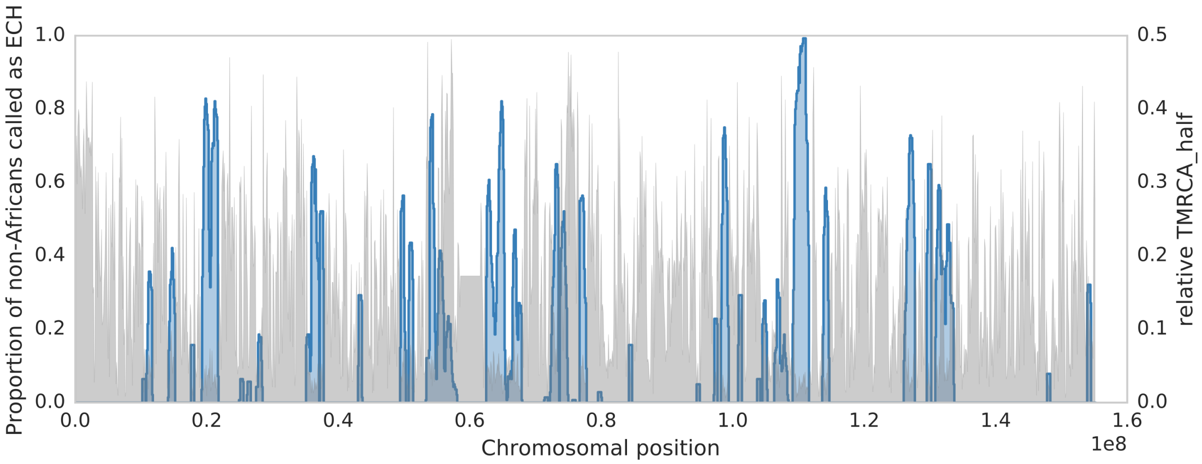


**Fig. S5.**

Proportion of non-African individuals that are part of a CNEH in each 100kb window (blue) with scale on the right axis. The relative $TMRCA_{half}$ (grey) with scale on the right axis.

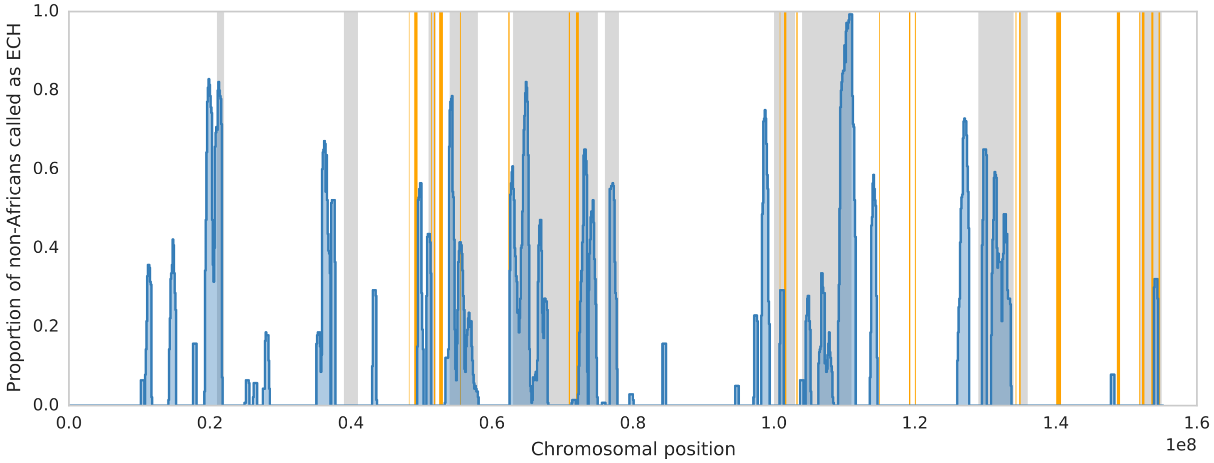


**Fig. S6.**

Location of ampliconic regions shown in orange.

| Sample ID | Population | Region |
|---|---|---|
| S_BantuHerero-1 | BantuHerero | Africa |
| S_BantuHerero-2 | BantuHerero | Africa |
| S_BantuTswana-2 | BantuTswana | Africa |
| S_BantuTswana-1 | BantuTswana | Africa |
| S_Biaka-1 | Biaka | Africa |
| S_Biaka-2 | Biaka | Africa |
| B_Dinka-3 | Dinka | Africa |
| S_Dinka-1 | Dinka | Africa |
| S_Dinka-2 | Dinka | Africa |
| S_Esan-1 | Esan | Africa |
| S_Ju_hoan_North-2 | Ju_hoan_North | Africa |
| S_Ju_hoan_North-3 | Ju_hoan_North | Africa |
| S_Ju_hoan_North-1 | Ju_hoan_North | Africa |
| B_Ju_hoan_North-4 | Ju_hoan_North | Africa |
| S_Mandenka-1 | Mandenka | Africa |
| B_Mandenka-3 | Mandenka | Africa |
| S_Mbuti-1 | Mbuti | Africa |
| S_Mbuti-3 | Mbuti | Africa |
| B_Mbuti-4 | Mbuti | Africa |
| S_Mende-1 | Mende | Africa |
| B_Yoruba-3 | Yoruba | Africa |
| S_Yoruba-2 | Yoruba | Africa |
| S_Chane-1 | Chane | America |
| S_Karitiana-1 | Karitiana | America |
| B_Karitiana-3 | Karitiana | America |
| S_Mixtec-1 | Mixtec | America |
| S_Pima-1 | Pima | America |
| S_Quechua-2 | Quechua | America |
| S_Zapotec-1 | Zapotec | America |
| S_Zapotec-2 | Zapotec | America |
| S_Aleut-1 | Aleut | CentralAsiaSiberia |
| S_Altaian-1 | Altaian | CentralAsiaSiberia |
| S_Chukchi-1 | Chukchi | CentralAsiaSiberia |
| S_Eskimo_Chaplin-1 | Eskimo_Chaplin | CentralAsiaSiberia |
| S_Eskimo_Sireniki-1 | Eskimo_Sireniki | CentralAsiaSiberia |
| S_Even-2 | Even | CentralAsiaSiberia |
| S_Kyrgyz-1 | Kyrgyz | CentralAsiaSiberia |
| S_Mongola-1 | Mongola | CentralAsiaSiberia |
| S_Tlingit-1 | Tlingit | CentralAsiaSiberia |
| S_Yakut-2 | Yakut | CentralAsiaSiberia |
| S_Ami-1 | Ami | EastAsia |
| S_Ami-2 | Ami | EastAsia |
| S_Atayal-1 | Atayal | EastAsia |
| S_Burmese-1 | Burmese | EastAsia |
| S_Burmese-2 | Burmese | EastAsia |
| S_Cambodian-1 | Cambodian | EastAsia |
| S_Dai-2 | Dai | EastAsia |
| B_Dai-4 | Dai | EastAsia |
| B_Han-3 | Han | EastAsia |
| S_Han-2 | Han | EastAsia |
| S_Hezhen-1 | Hezhen | EastAsia |
| S_Japanese-1 | Japanese | EastAsia |
| S_Japanese-3 | Japanese | EastAsia |
| S_Kinh-2 | Kinh | EastAsia |
| S_Korean-1 | Korean | EastAsia |
| S_Lahu-2 | Lahu | EastAsia |
| S_Miao-1 | Miao | EastAsia |
| S_Naxi-1 | Naxi | EastAsia |
| S_Oroqen-1 | Oroqen | EastAsia |
| S_She-2 | She | EastAsia |
| S_Thai-1 | Thai | EastAsia |
| S_Tu-1 | Tu | EastAsia |
| S_Tujia-1 | Tujia | EastAsia |
| S_Uygur-2 | Uygur | EastAsia |
| S_Xibo-2 | Xibo | EastAsia |

| | | |
|---|---|---|
| S_Xibo-1 | Xibo | EastAsia |
| S_Yi-1 | Yi | EastAsia |
| B_Australian-4 | Australian | Oceania |
| S_Hawaiian-1 | Hawaiian | Oceania |
| S_Igorot-2 | Igorot | Oceania |
| S_Maori-1 | Maori | Oceania |
| S_Papuan-6 | Papuan | Oceania |
| S_Papuan-7 | Papuan | Oceania |
| S_Papuan-5 | Papuan | Oceania |
| S_Papuan-8 | Papuan | Oceania |
| S_Papuan-4 | Papuan | Oceania |
| S_Papuan-12 | Papuan | Oceania |
| S_Papuan-10 | Papuan | Oceania |
| S_Papuan-3 | Papuan | Oceania |
| B_Papuan-15 | Papuan | Oceania |
| S_Papuan-9 | Papuan | Oceania |
| S_Papuan-2 | Papuan | Oceania |
| S_Papuan-11 | Papuan | Oceania |
| S_Balochi-2 | Balochi | SouthAsia |
| S_Balochi-1 | Balochi | SouthAsia |
| S_Bengali-1 | Bengali | SouthAsia |
| S_Brahmin-1 | Brahmin | SouthAsia |
| S_Brahmin-2 | Brahmin | SouthAsia |
| S_Brahui-2 | Brahui | SouthAsia |
| S_Brahui-1 | Brahui | SouthAsia |
| S_Burusho-1 | Burusho | SouthAsia |
| S_Hazara-1 | Hazara | SouthAsia |
| S_Hazara-2 | Hazara | SouthAsia |
| S_Irula-1 | Irula | SouthAsia |
| S_Irula-2 | Irula | SouthAsia |
| S_Kalash-1 | Kalash | SouthAsia |
| S_Kapu-1 | Kapu | SouthAsia |
| S_Kapu-2 | Kapu | SouthAsia |
| S_Khonda_Dora-1 | Khonda_Dora | SouthAsia |
| S_Kusunda-2 | Kusunda | SouthAsia |
| S_Kusunda-1 | Kusunda | SouthAsia |
| S_Madiga-2 | Madiga | SouthAsia |
| S_Madiga-1 | Madiga | SouthAsia |
| S_Makrani-1 | Makrani | SouthAsia |
| S_Mala-3 | Mala | SouthAsia |
| S_Mala-2 | Mala | SouthAsia |
| S_Pathan-1 | Pathan | SouthAsia |
| S_Punjabi-2 | Punjabi | SouthAsia |
| S_Punjabi-1 | Punjabi | SouthAsia |
| S_Relli-1 | Relli | SouthAsia |
| S_Relli-2 | Relli | SouthAsia |
| S_Sindhi-1 | Sindhi | SouthAsia |
| S_Yadava-2 | Yadava | SouthAsia |
| S_Yadava-1 | Yadava | SouthAsia |
| S_Abkhasian-2 | Abkhasian | WestEurasia |
| S_Abkhasian-1 | Abkhasian | WestEurasia |
| S_Adygei-1 | Adygei | WestEurasia |
| S_Armenian-2 | Armenian | WestEurasia |
| S_Armenian-1 | Armenian | WestEurasia |
| S_Basque-1 | Basque | WestEurasia |
| S_BedouinB-1 | BedouinB | WestEurasia |
| S_Bergamo-1 | Bergamo | WestEurasia |
| S_Bulgarian-1 | Bulgarian | WestEurasia |
| S_Bulgarian-2 | Bulgarian | WestEurasia |
| S_Chechen-1 | Chechen | WestEurasia |
| B_Crete-2 | Crete | WestEurasia |
| S_Czech-2 | Czech | WestEurasia |
| S_Druze-2 | Druze | WestEurasia |
| S_English-1 | English | WestEurasia |
| S_Estonian-1 | Estonian | WestEurasia |
| S_Estonian-2 | Estonian | WestEurasia |

| | | |
|---|---|---|
| S_Finnish-3 | Finnish | WestEurasia |
| S_French-1 | French | WestEurasia |
| B_French-3 | French | WestEurasia |
| S_Georgian-2 | Georgian | WestEurasia |
| S_Georgian-1 | Georgian | WestEurasia |
| S_Greek-2 | Greek | WestEurasia |
| S_Greek-1 | Greek | WestEurasia |
| S_Hungarian-2 | Hungarian | WestEurasia |
| S_Iranian-1 | Iranian | WestEurasia |
| S_Iranian-2 | Iranian | WestEurasia |
| S_Iraqi_Jew-2 | Iraqi_Jew | WestEurasia |
| S_Jordanian-3 | Jordanian | WestEurasia |
| S_Jordanian-2 | Jordanian | WestEurasia |
| S_Jordanian-1 | Jordanian | WestEurasia |
| S_Lezgin-2 | Lezgin | WestEurasia |
| S_North_Ossetian-1 | North_Ossetian | WestEurasia |
| S_North_Ossetian-2 | North_Ossetian | WestEurasia |
| S_Orcadian-1 | Orcadian | WestEurasia |
| S_Palestinian-1 | Palestinian | WestEurasia |
| S_Polish-1 | Polish | WestEurasia |
| S_Russian-1 | Russian | WestEurasia |
| S_Saami-2 | Saami | WestEurasia |
| S_Samaritan-1 | Samaritan | WestEurasia |
| S_Sardinian-1 | Sardinian | WestEurasia |
| B_Sardinian-3 | Sardinian | WestEurasia |
| S_Spanish-1 | Spanish | WestEurasia |
| S_Tajik-1 | Tajik | WestEurasia |
| S_Tajik-2 | Tajik | WestEurasia |
| S_Turkish-1 | Turkish | WestEurasia |
| S_Tuscan-2 | Tuscan | WestEurasia |
| S_Yemenite_Jew-2 | Yemenite_Jew | WestEurasia |

**Table S1.**

Sample ID, population and region for each of the 162 males, whos X chromosome was analyzed.

| Peaks | | | 90%-regions | | | | | 75%-regions | | | | | Data S1 |
|---|---|---|---|---|---|---|---|---|---|---|---|---|---|
| Start | End | Freq. ECH | Start | End | Length | Min freq. ECH | Freq. ECHs spanning region | Start | End | Length | Min freq. ECH | Freq. ECHs spanning region | Plot identifier |
| 112 | 114 | 0.36 | 111 | 115 | 4 | 0.33 | 0.32 | 111 | 117 | 6 | 0.31 | 0.28 | A |
| 147 | 148 | 0.42 | 147 | 149 | 2 | 0.41 | 0.41 | 144 | 150 | 6 | 0.32 | 0.22 | B |
| 198 | 199 | 0.83 | 196 | 202 | 6 | 0.76 | 0.67 | 195 | 203 | 8 | 0.64 | 0.54 | C |
| 212 | 213 | 0.82 | 211 | 216 | 5 | 0.78 | 0.74 | 207 | 217 | 10 | 0.66 | 0.53 | D |
| 362 | 363 | 0.67 | 360 | 364 | 4 | 0.64 | 0.63 | 359 | 367 | 8 | 0.52 | 0.31 | E |
| 373 | 377 | 0.52 | 372 | 377 | 5 | 0.51 | 0.51 | 372 | 377 | 5 | 0.51 | 0.51 | F |
| 431 | 435 | 0.29 | 431 | 436 | 5 | 0.28 | 0.28 | 431 | 436 | 5 | 0.28 | 0.28 | G |
| 498 | 500 | 0.56 | 495 | 500 | 5 | 0.52 | 0.52 | 495 | 500 | 5 | 0.52 | 0.52 | H |
| 509 | 512 | 0.44 | 508 | 513 | 5 | 0.41 | 0.41 | 508 | 514 | 6 | 0.34 | 0.33 | I |
| 543 | 544 | 0.79 | 540 | 544 | 4 | 0.75 | 0.75 | 539 | 544 | 5 | 0.66 | 0.66 | J |
| 554 | 556 | 0.41 | 553 | 558 | 5 | 0.39 | 0.3 | 552 | 559 | 7 | 0.34 | 0.2 | K |
| 629 | 630 | 0.61 | 627 | 631 | 4 | 0.55 | 0.51 | 626 | 632 | 6 | 0.52 | 0.37 | L |
| 648 | 649 | 0.82 | 647 | 651 | 4 | 0.77 | 0.75 | 644 | 652 | 8 | 0.64 | 0.44 | M |
| 668 | 670 | 0.47 | 667 | 670 | 3 | 0.46 | 0.45 | 666 | 671 | 5 | 0.39 | 0.32 | N |
| 674 | 676 | 0.27 | 674 | 678 | 4 | 0.26 | 0.26 | 674 | 679 | 5 | 0.24 | 0.24 | O |
| 731 | 733 | 0.65 | 730 | 735 | 5 | 0.59 | 0.58 | 730 | 735 | 5 | 0.59 | 0.58 | P |
| 743 | 744 | 0.52 | 740 | 744 | 4 | 0.48 | 0.46 | 739 | 745 | 6 | 0.42 | 0.34 | Q |
| 771 | 772 | 0.56 | 767 | 774 | 7 | 0.52 | 0.51 | 767 | 774 | 7 | 0.52 | 0.51 | R |
| 987 | 988 | 0.75 | 985 | 989 | 4 | 0.72 | 0.69 | 985 | 991 | 6 | 0.59 | 0.54 | S |
| 1009 | 1014 | 0.29 | 1009 | 1014 | 5 | 0.29 | 0.29 | 1009 | 1014 | 5 | 0.29 | 0.29 | T |
| 1048 | 1050 | 0.28 | 1047 | 1050 | 3 | 0.27 | 0.27 | 1045 | 1051 | 6 | 0.21 | 0.15 | U |
| 1067 | 1069 | 0.34 | 1067 | 1069 | 2 | 0.34 | 0.32 | 1067 | 1072 | 5 | 0.29 | 0.25 | V |
| 1107 | 1111 | 0.99 | 1102 | 1111 | 9 | 0.95 | 0.91 | 1094 | 1112 | 18 | 0.76 | 0.53 | X |
| 1141 | 1142 | 0.59 | 1140 | 1143 | 3 | 0.55 | 0.52 | 1138 | 1145 | 7 | 0.46 | 0.37 | Y |
| 1270 | 1271 | 0.73 | 1268 | 1274 | 6 | 0.68 | 0.6 | 1268 | 1275 | 7 | 0.61 | 0.52 | Z |
| 1297 | 1301 | 0.65 | 1297 | 1302 | 5 | 0.64 | 0.64 | 1297 | 1302 | 5 | 0.64 | 0.64 | $ |
| 1313 | 1314 | 0.59 | 1312 | 1316 | 4 | 0.58 | 0.56 | 1311 | 1316 | 5 | 0.47 | 0.46 | % |
| 1326 | 1329 | 0.49 | 1326 | 1329 | 3 | 0.49 | 0.49 | 1324 | 1331 | 7 | 0.37 | 0.26 | & |
| 1540 | 1544 | 0.32 | 1539 | 1544 | 5 | 0.3 | 0.3 | 1539 | 1545 | 6 | 0.27 | 0.25 | # |

**Table S2.**

Peaks in frequency ECHs called and regions around such peaks. The table gives genomic coordinates in 100kb (hg19) of each peak in the proportion of males called as ECH. The table also gives coordinates of 90%-regions and 75%-regions around each with the minimum proportion of non-African males that are called as ECH across the region (Min freq. ECH) and the proportion of non-African males ECH that span the entire region (Freq. ECHs spanning region).

**Data S1. (separate file)**

Haplotype structure visualizations for each identified region as in Figure 3 of the main text. The left side of each figure is a UPGMA tree of the individual haplotypes shown as horizontal lines on the right. Haplotypes are color-coded according to geographical region. The ancient Ust'-Ishim individual is marked with grey. Vertical black bars on each haplotype represents non-reference SNPs. The last column in Table S2 links each chromosomal region to a separate haplotype plot.
