## Supplementary material for "Extreme selective sweeps displaced archaic admixture across the human X chromosome around 50,000 years ago": Data S1

A

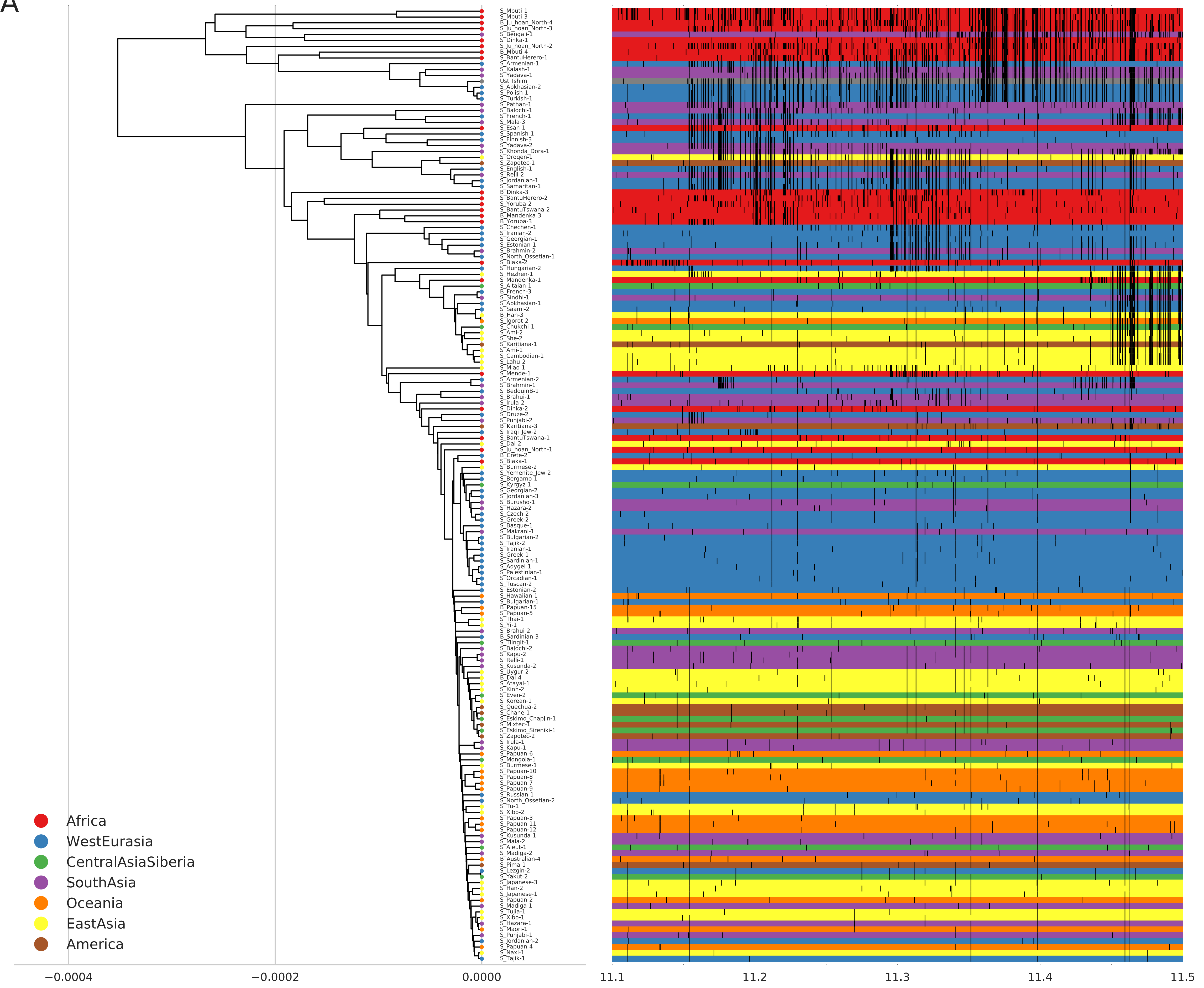

S_Mbuti-1
S_Mbuti-3
B_Ju_hoan_North-4
S_Ju_hoan_North-3
S_Bengali-1
S_Dinka-1
S_Ju_hoan_North-2
B_Mbuti-4
S_BantuHerero-1
S_Armenian-1
S_Kalash-1
S_Yadava-1
Ust_Ishim
S_Abkhasian-2
S_Polish-1
S_Turkish-1
S_Pathan-1
S_Balochi-1
S_French-1
S_Mala-3
S_Esan-1
S_Spanish-1
S_Finnish-3
S_Yadava-2
S_Khonda_Dora-1
S_Oroqen-1
S_Zapotec-1
S_English-1
S_Relli-2
S_Jordanian-1
S_Samaritan-1
B_Dinka-3
S_BantuHerero-2
S_Yoruba-2
S_BantuTswana-2
B_Mandenka-3
B_Yoruba-3
S_Chechen-1
S_Iranian-2
S_Georgian-1
S_Estonian-1
S_Brahmin-2
S_North_Ossetian-1
S_Biaka-2
S_Hungarian-2
S_Hezhen-1
S_Mandenka-1
S_Altaian-1
B_French-3
S_Sindhi-1
S_Abkhasian-1
S_Saami-2
B_Han-3
S_Igorot-2
S_Chukchi-1
S_Ami-2
S_She-2
S_Karitiana-1
S_Ami-1
S_Cambodian-1
S_Lahu-2
S_Miao-1
S_Mende-1
S_Armenian-2
S_Brahmin-1
S_BedouinB-1
S_Brahui-1
S_Irula-2
S_Dinka-2
S_Druze-2
S_Punjabi-2
B_Karitiana-3
S_Iraqi_Jew-2
S_BantuTswana-1
S_Dai-2
S_Ju_hoan_North-1
B_Crete-2
S_Biaka-1
S_Burmese-2
S_Yemenite_Jew-2
S_Bergamo-1
S_Kyrgyz-1
S_Georgian-2
S_Jordanian-3
S_Burusho-1
S_Hazara-2
S_Czech-2
S_Greek-2
S_Basque-1
S_Makrani-1
S_Bulgarian-2
S_Tajik-2
S_Iranian-1
S_Greek-1
S_Sardinian-1
S_Adygei-1
S_Palestinian-1
S_Orcadian-1
S_Tuscan-2
S_Estonian-2
S_Hawaiian-1
S_Bulgarian-1
B_Papuan-15
S_Papuan-5
S_Thai-1
S_Yi-1
S_Brahui-2
B_Sardinian-3
S_Tlingit-1
S_Balochi-2
S_Kapu-2
S_Relli-1
S_Kusunda-2
S_Uygur-2
B_Dai-4
S_Atayal-1
S_Kinh-2
S_Even-2
S_Korean-1
S_Quechua-2
S_Chane-1
S_Eskimo_Chaplin-1
S_Mixtec-1
S_Eskimo_Sireniki-1
S_Zapotec-2
S_Irula-1
S_Kapu-1
S_Papuan-6
S_Mongola-1
S_Burmese-1
S_Papuan-10
S_Papuan-8
S_Papuan-7
S_Papuan-9
S_Russian-1
S_North_Ossetian-2
S_Tu-1
S_Xibo-2
S_Papuan-3
S_Papuan-11
S_Papuan-12
S_Kusunda-1
S_Mala-2
S_Aleut-1
S_Madiga-2
B_Australian-4
S_Pima-1
S_Lezgin-2
S_Yakut-2
S_Japanese-3
S_Han-2
S_Japanese-1
S_Papuan-2
S_Madiga-1
S_Tujia-1
S_Xibo-1
S_Hazara-1
S_Maori-1
S_Punjabi-1
S_Jordanian-2
S_Papuan-4
S_Naxi-1
S_Tajik-1
Africa
WestEurasia
CentralAsiaSiberia
SouthAsia
Oceania
EastAsia
America
−0.0004
−0.0002
0.0000
11.1
11.2
11.3
11.4
11.5

B

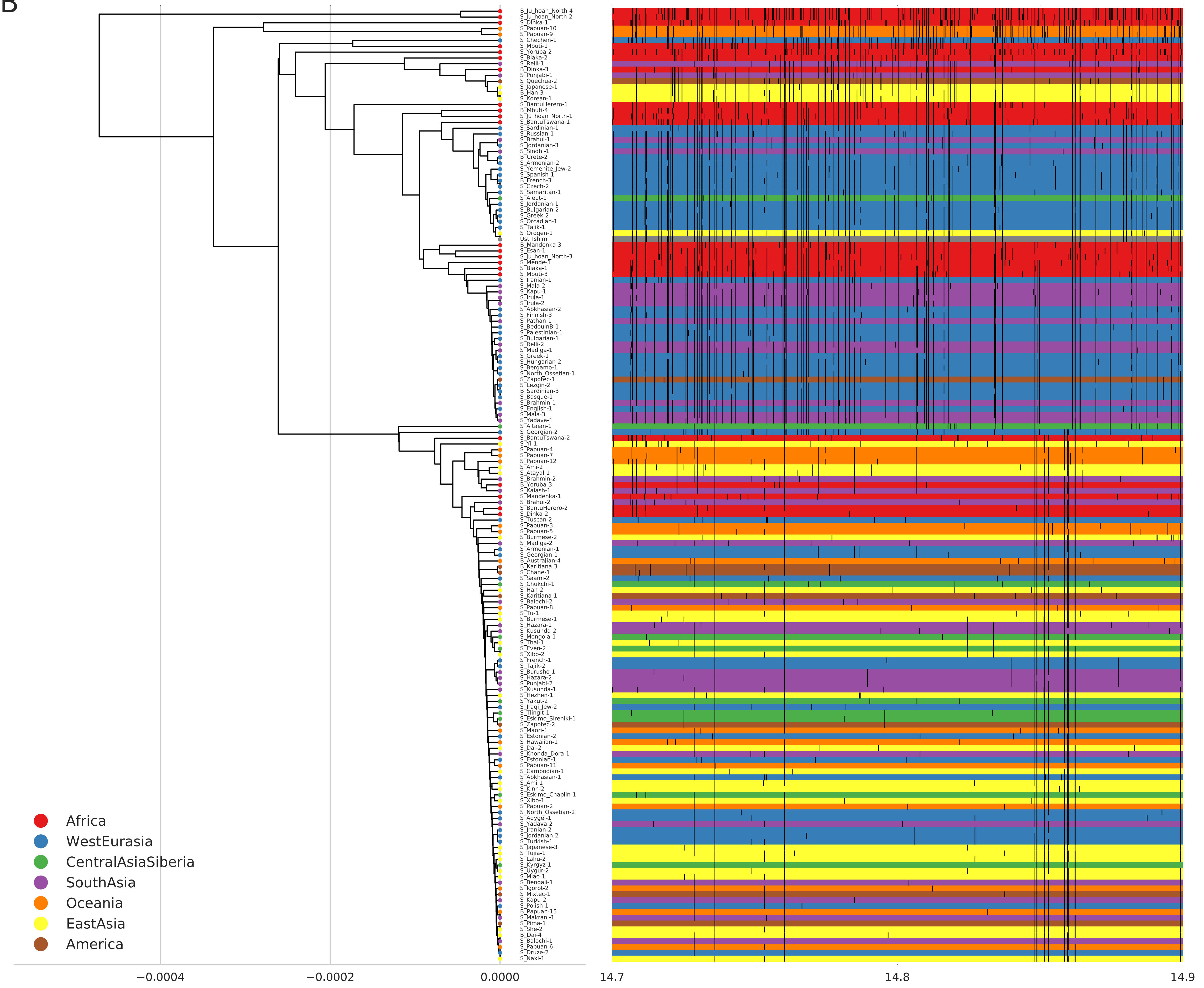

B_Ju_hoan_North-4
S_Ju_hoan_North-2
S_Dinka-1
S_Papuan-10
S_Papuan-9
S_Chechen-1
S_Mbuti-1
S_Yoruba-2
S_Biaka-2
S_Relli-1
B_Dinka-3
S_Punjabi-1
S_Quechua-2
S_Japanese-1
B_Han-3
S_Korean-1
S_BantuHerero-1
B_Mbuti-4
S_Ju_hoan_North-1
S_BantuTswana-1
S_Sardinian-1
S_Russian-1
S_Brahui-1
S_Jordanian-3
S_Sindhi-1
B_Crete-2
S_Armenian-2
S_Yemenite_Jew-2
S_Spanish-1
B_French-3
S_Czech-2
S_Samaritan-1
S_Aleut-1
S_Jordanian-1
S_Bulgarian-2
S_Greek-2
S_Orcadian-1
S_Tajik-1
S_Oroqen-1
Ust_Ishim
B_Mandenka-3
S_Esan-1
S_Ju_hoan_North-3
S_Mende-1
S_Biaka-1
S_Mbuti-3
S_Iranian-1
S_Mala-2
S_Kapu-1
S_Irula-1
S_Irula-2
S_Abkhasian-2
S_Finnish-3
S_Pathan-1
S_BedouinB-1
S_Palestinian-1
S_Bulgarian-1
S_Relli-2
S_Madiga-1
S_Greek-1
S_Hungarian-2
S_Bergamo-1
S_North_Ossetian-1
S_Zapotec-1
S_Lezgin-2
B_Sardinian-3
S_Basque-1
S_Brahmin-1
S_English-1
S_Mala-3
S_Yadava-1
S_Altaian-1
S_Georgian-2
S_BantuTswana-2
S_Yi-1
S_Papuan-4
S_Papuan-7
S_Papuan-12
S_Ami-2
S_Atayal-1
S_Brahmin-2
B_Yoruba-3
S_Kalash-1
S_Mandenka-1
S_Brahui-2
S_BantuHerero-2
S_Dinka-2
S_Tuscan-2
S_Papuan-3
S_Papuan-5
S_Burmese-2
S_Madiga-2
S_Armenian-1
S_Georgian-1
B_Australian-4
B_Karitiana-3
S_Chane-1
S_Saami-2
S_Chukchi-1
S_Han-2
S_Karitiana-1
S_Balochi-2
S_Papuan-8
S_Tu-1
S_Burmese-1
S_Hazara-1
S_Kusunda-2
S_Mongola-1
S_Thai-1
S_Even-2
S_Xibo-2
S_French-1
S_Tajik-2
S_Burusho-1
S_Hazara-2
S_Punjabi-2
S_Kusunda-1
S_Hezhen-1
S_Yakut-2
S_Iraqi_Jew-2
S_Tlingit-1
S_Eskimo_Sireniki-1
S_Zapotec-2
S_Maori-1
S_Estonian-2
S_Hawaiian-1
S_Dai-2
S_Khonda_Dora-1
S_Estonian-1
S_Papuan-11
S_Cambodian-1
S_Abkhasian-1
S_Ami-1
S_Kinh-2
S_Eskimo_Chaplin-1
S_Xibo-1
S_Papuan-2
S_North_Ossetian-2
S_Adygei-1
S_Yadava-2
S_Iranian-2
S_Jordanian-2
S_Turkish-1
S_Japanese-3
S_Tujia-1
S_Lahu-2
S_Kyrgyz-1
S_Uygur-2
S_Miao-1
S_Bengali-1
S_Igorot-2
S_Mixtec-1
S_Kapu-2
S_Polish-1
B_Papuan-15
S_Makrani-1
S_Pima-1
S_She-2
B_Dai-4
S_Balochi-1
S_Papuan-6
S_Druze-2
S_Naxi-1
Africa
WestEurasia
CentralAsiaSiberia
SouthAsia
Oceania
EastAsia
America
−0.0004
−0.0002
0.0000
14.7
14.8
14.9

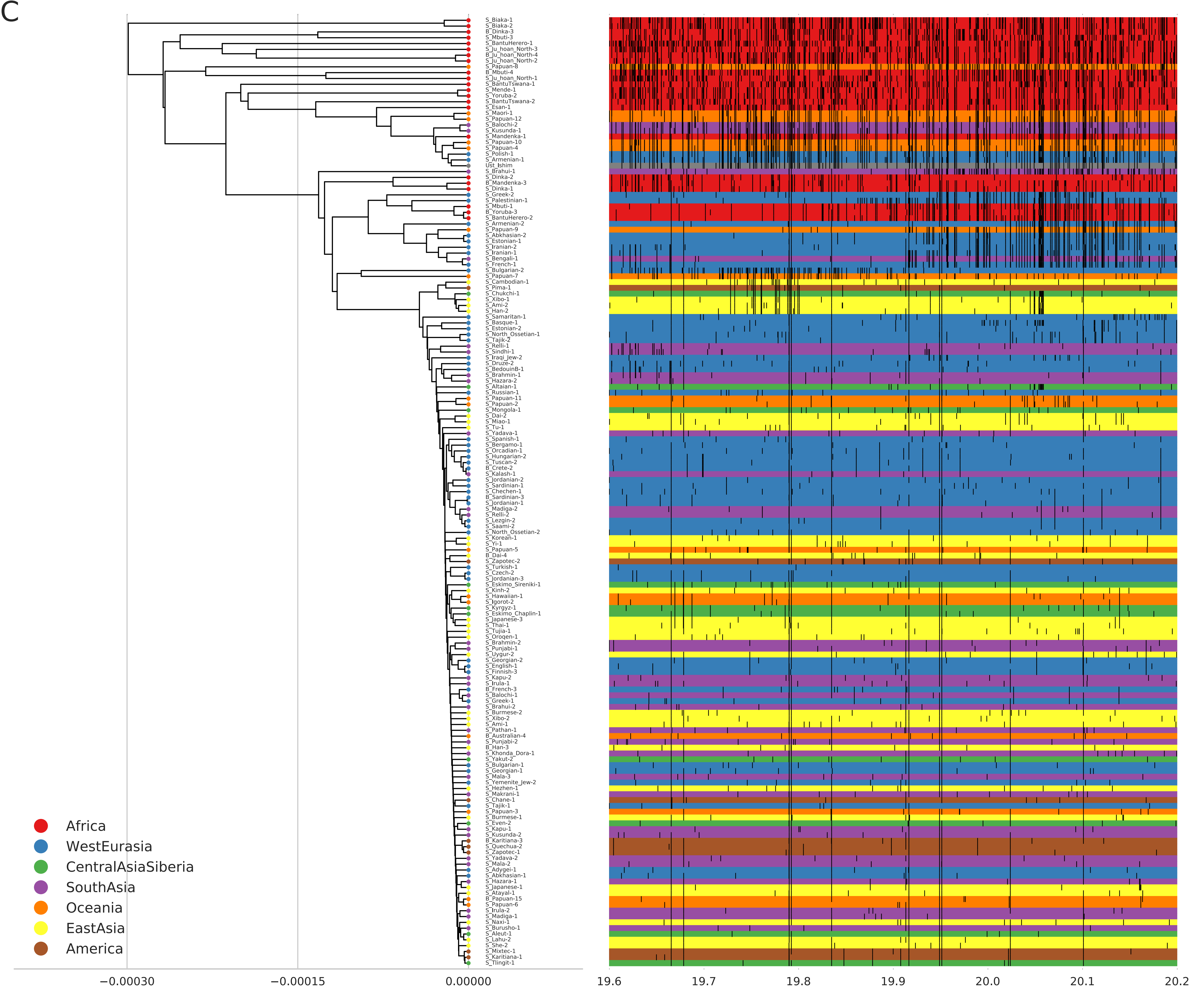
C
Africa
WestEurasia
CentralAsiaSiberia
SouthAsia
Oceania
EastAsia
America
−0.00030
−0.00015
0.00000
19.6
19.7
19.8
19.9
20.0
20.1
20.2

D

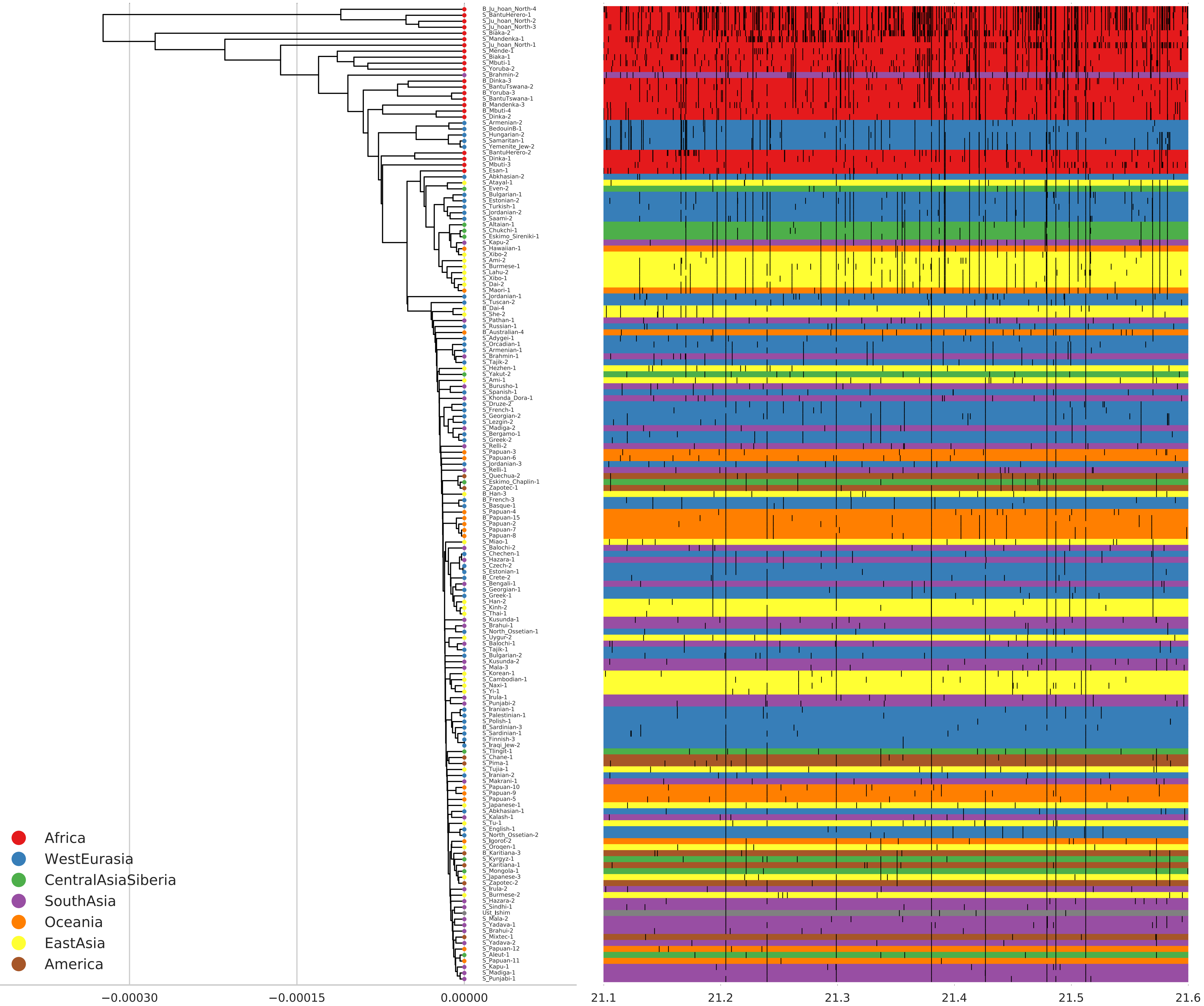

B_Ju_hoan_North-4
S_BantuHerero-1
S_Ju_hoan_North-2
S_Ju_hoan_North-3
S_Biaka-2
S_Mandenka-1
S_Ju_hoan_North-1
S_Mende-1
S_Biaka-1
S_Mbuti-1
S_Yoruba-2
S_Brahmin-2
B_Dinka-3
S_BantuTswana-2
B_Yoruba-3
S_BantuTswana-1
B_Mandenka-3
B_Mbuti-4
S_Dinka-2
S_Armenian-2
S_BedouinB-1
S_Hungarian-2
S_Samaritan-1
S_Yemenite_Jew-2
S_BantuHerero-2
S_Dinka-1
S_Mbuti-3
S_Esan-1
S_Abkhasian-2
S_Atayal-1
S_Even-2
S_Bulgarian-1
S_Estonian-2
S_Turkish-1
S_Jordanian-2
S_Saami-2
S_Altaian-1
S_Chukchi-1
S_Eskimo_Sireniki-1
S_Kapu-2
S_Hawaiian-1
S_Xibo-2
S_Ami-2
S_Burmese-1
S_Lahu-2
S_Xibo-1
S_Dai-2
S_Maori-1
S_Jordanian-1
S_Tuscan-2
B_Dai-4
S_She-2
S_Pathan-1
S_Russian-1
B_Australian-4
S_Adygei-1
S_Orcadian-1
S_Armenian-1
S_Brahmin-1
S_Tajik-2
S_Hezhen-1
S_Yakut-2
S_Ami-1
S_Burusho-1
S_Spanish-1
S_Khonda_Dora-1
S_Druze-2
S_French-1
S_Georgian-2
S_Lezgin-2
S_Madiga-2
S_Bergamo-1
S_Greek-2
S_Relli-2
S_Papuan-3
S_Papuan-6
S_Jordanian-3
S_Relli-1
S_Quechua-2
S_Eskimo_Chaplin-1
S_Zapotec-1
B_Han-3
B_French-3
S_Basque-1
S_Papuan-4
B_Papuan-15
S_Papuan-2
S_Papuan-7
S_Papuan-8
S_Miao-1
S_Balochi-2
S_Chechen-1
S_Hazara-1
S_Czech-2
S_Estonian-1
B_Crete-2
S_Bengali-1
S_Georgian-1
S_Greek-1
S_Han-2
S_Kinh-2
S_Thai-1
S_Kusunda-1
S_Brahui-1
S_North_Ossetian-1
S_Uygur-2
S_Balochi-1
S_Tajik-1
S_Bulgarian-2
S_Kusunda-2
S_Mala-3
S_Korean-1
S_Cambodian-1
S_Naxi-1
S_Yi-1
S_Irula-1
S_Punjabi-2
S_Iranian-1
S_Palestinian-1
S_Polish-1
B_Sardinian-3
S_Sardinian-1
S_Finnish-3
S_Iraqi_Jew-2
S_Tlingit-1
S_Chane-1
S_Pima-1
S_Tujia-1
S_Iranian-2
S_Makrani-1
S_Papuan-10
S_Papuan-9
S_Papuan-5
S_Japanese-1
S_Abkhasian-1
S_Kalash-1
S_Tu-1
S_English-1
S_North_Ossetian-2
S_Igorot-2
S_Oroqen-1
B_Karitiana-3
S_Kyrgyz-1
S_Karitiana-1
S_Mongola-1
S_Japanese-3
S_Zapotec-2
S_Irula-2
S_Burmese-2
S_Hazara-2
S_Sindhi-1
Ust_Ishim
S_Mala-2
S_Yadava-1
S_Brahui-2
S_Mixtec-1
S_Yadava-2
S_Papuan-12
S_Aleut-1
S_Papuan-11
S_Kapu-1
S_Madiga-1
S_Punjabi-1
Africa
WestEurasia
CentralAsiaSiberia
SouthAsia
Oceania
EastAsia
America
−0.00030
−0.00015
0.00000
21.1
21.2
21.3
21.4
21.5
21.6

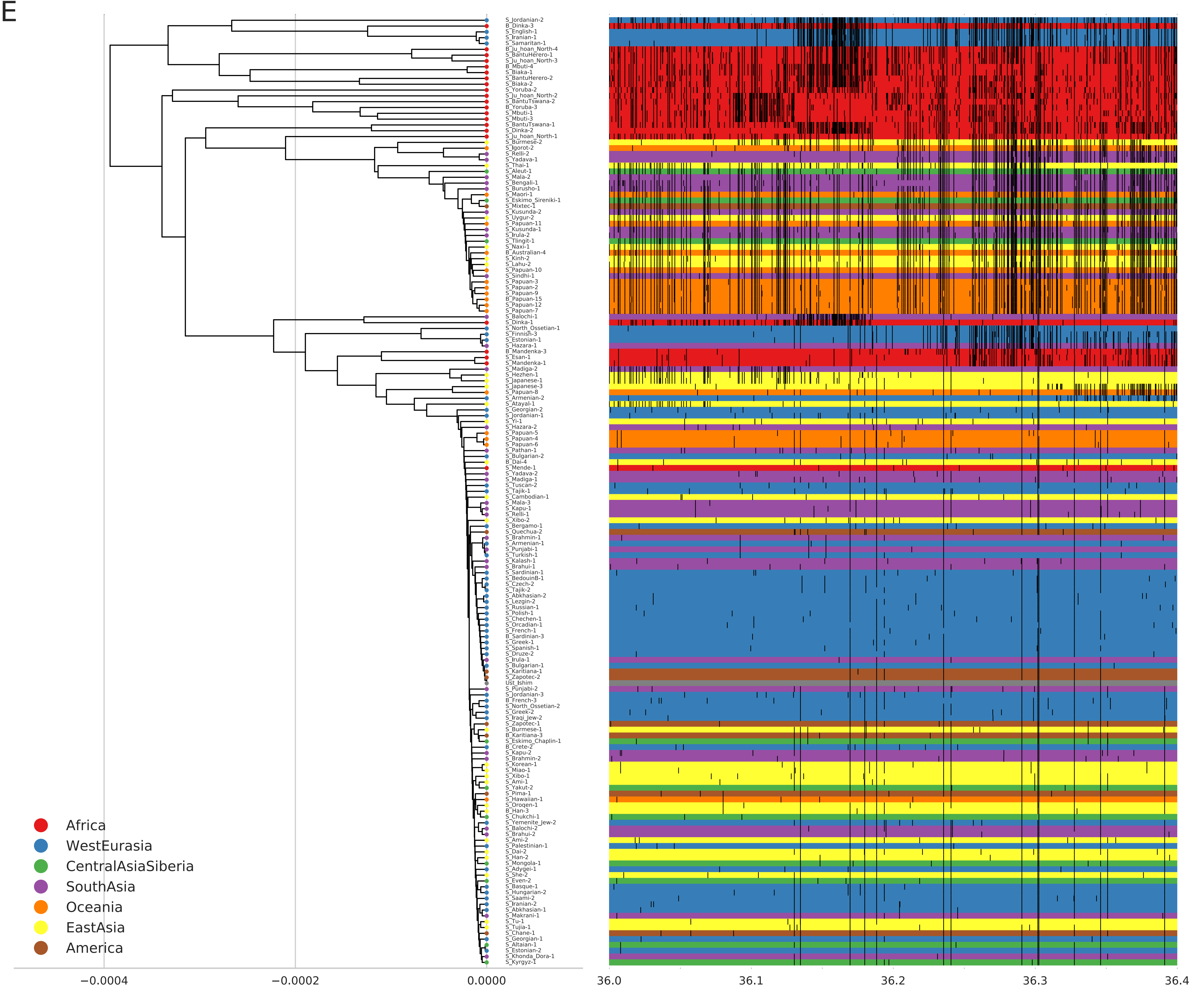
E
S_Jordanian-2
B_Dinka-3
S_English-1
S_Iranian-1
S_Samaritan-1
B_Ju_hoan_North-4
S_BantuHerero-1
S_Ju_hoan_North-3
B_Mbuti-4
S_Biaka-1
S_BantuHerero-2
S_Biaka-2
S_Yoruba-2
S_Ju_hoan_North-2
S_BantuTswana-2
B_Yoruba-3
S_Mbuti-1
S_Mbuti-3
S_BantuTswana-1
S_Dinka-2
S_Ju_hoan_North-1
S_Burmese-2
S_Igorot-2
S_Relli-2
S_Yadava-1
S_Thai-1
S_Aleut-1
S_Mala-2
S_Bengali-1
S_Burusho-1
S_Maori-1
S_Eskimo_Sireniki-1
S_Mixtec-1
S_Kusunda-2
S_Uygur-2
S_Papuan-11
S_Kusunda-1
S_Irula-2
S_Tlingit-1
S_Naxi-1
B_Australian-4
S_Kinh-2
S_Lahu-2
S_Papuan-10
S_Sindhi-1
S_Papuan-3
S_Papuan-2
S_Papuan-9
B_Papuan-15
S_Papuan-12
S_Papuan-7
S_Balochi-1
S_Dinka-1
S_North_Ossetian-1
S_Finnish-3
S_Estonian-1
S_Hazara-1
B_Mandenka-3
S_Esan-1
S_Mandenka-1
S_Madiga-2
S_Hezhen-1
S_Japanese-1
S_Japanese-3
S_Papuan-8
S_Armenian-2
S_Atayal-1
S_Georgian-2
S_Jordanian-1
S_Yi-1
S_Hazara-2
S_Papuan-5
S_Papuan-4
S_Papuan-6
S_Pathan-1
S_Bulgarian-2
B_Dai-4
S_Mende-1
S_Yadava-2
S_Madiga-1
S_Tuscan-2
S_Tajik-1
S_Cambodian-1
S_Mala-3
S_Kapu-1
S_Relli-1
S_Xibo-2
S_Bergamo-1
S_Quechua-2
S_Brahmin-1
S_Armenian-1
S_Punjabi-1
S_Turkish-1
S_Kalash-1
S_Brahui-1
S_Sardinian-1
S_BedouinB-1
S_Czech-2
S_Tajik-2
S_Abkhasian-2
S_Lezgin-2
S_Russian-1
S_Polish-1
S_Chechen-1
S_Orcadian-1
S_French-1
B_Sardinian-3
S_Greek-1
S_Spanish-1
S_Druze-2
S_Irula-1
S_Bulgarian-1
S_Karitiana-1
S_Zapotec-2
Ust_Ishim
S_Punjabi-2
S_Jordanian-3
B_French-3
S_North_Ossetian-2
S_Greek-2
S_Iraqi_Jew-2
S_Zapotec-1
S_Burmese-1
B_Karitiana-3
S_Eskimo_Chaplin-1
B_Crete-2
S_Kapu-2
S_Brahmin-2
S_Korean-1
S_Miao-1
S_Xibo-1
S_Ami-1
S_Yakut-2
S_Pima-1
S_Hawaiian-1
S_Oroqen-1
B_Han-3
S_Chukchi-1
S_Yemenite_Jew-2
S_Balochi-2
S_Brahui-2
S_Ami-2
S_Palestinian-1
S_Dai-2
S_Han-2
S_Mongola-1
S_Adygei-1
S_She-2
S_Even-2
S_Basque-1
S_Hungarian-2
S_Saami-2
S_Iranian-2
S_Abkhasian-1
S_Makrani-1
S_Tu-1
S_Tujia-1
S_Chane-1
S_Georgian-1
S_Altaian-1
S_Estonian-2
S_Khonda_Dora-1
S_Kyrgyz-1
Africa
WestEurasia
CentralAsiaSiberia
SouthAsia
Oceania
EastAsia
America
−0.0004
−0.0002
0.0000
36.0
36.1
36.2
36.3
36.4

F

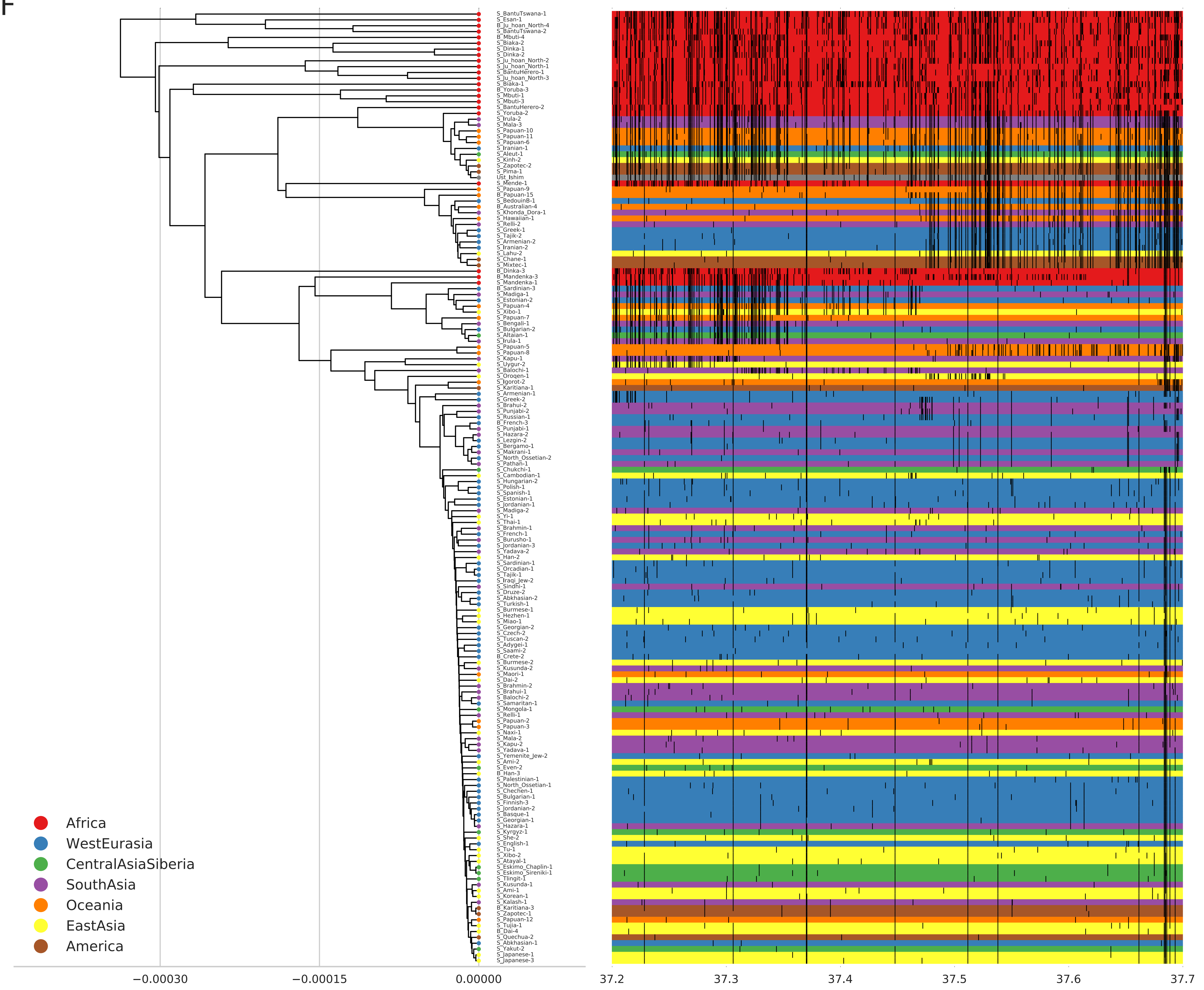

S_BantuTswana-1
S_Esan-1
B_Ju_hoan_North-4
S_BantuTswana-2
B_Mbuti-4
S_Biaka-2
S_Dinka-1
S_Dinka-2
S_Ju_hoan_North-2
S_Ju_hoan_North-1
S_BantuHerero-1
S_Ju_hoan_North-3
S_Biaka-1
B_Yoruba-3
S_Mbuti-1
S_Mbuti-3
S_BantuHerero-2
S_Yoruba-2
S_Irula-2
S_Mala-3
S_Papuan-10
S_Papuan-11
S_Papuan-6
S_Iranian-1
S_Aleut-1
S_Kinh-2
S_Zapotec-2
S_Pima-1
Ust_Ishim
S_Mende-1
S_Papuan-9
B_Papuan-15
S_BedouinB-1
B_Australian-4
S_Khonda_Dora-1
S_Hawaiian-1
S_Relli-2
S_Greek-1
S_Tajik-2
S_Armenian-2
S_Iranian-2
S_Lahu-2
S_Chane-1
S_Mixtec-1
B_Dinka-3
B_Mandenka-3
S_Mandenka-1
B_Sardinian-3
S_Madiga-1
S_Estonian-2
S_Papuan-4
S_Xibo-1
S_Papuan-7
S_Bengali-1
S_Bulgarian-2
S_Altaian-1
S_Irula-1
S_Papuan-5
S_Papuan-8
S_Kapu-1
S_Uygur-2
S_Balochi-1
S_Oroqen-1
S_Igorot-2
S_Karitiana-1
S_Armenian-1
S_Greek-2
S_Brahui-2
S_Punjabi-2
S_Russian-1
B_French-3
S_Punjabi-1
S_Hazara-2
S_Lezgin-2
S_Bergamo-1
S_Makrani-1
S_North_Ossetian-2
S_Pathan-1
S_Chukchi-1
S_Cambodian-1
S_Hungarian-2
S_Polish-1
S_Spanish-1
S_Estonian-1
S_Jordanian-1
S_Madiga-2
S_Yi-1
S_Thai-1
S_Brahmin-1
S_French-1
S_Burusho-1
S_Jordanian-3
S_Yadava-2
S_Han-2
S_Sardinian-1
S_Orcadian-1
S_Tajik-1
S_Iraqi_Jew-2
S_Sindhi-1
S_Druze-2
S_Abkhasian-2
S_Turkish-1
S_Burmese-1
S_Hezhen-1
S_Miao-1
S_Georgian-2
S_Czech-2
S_Tuscan-2
S_Adygei-1
S_Saami-2
B_Crete-2
S_Burmese-2
S_Kusunda-2
S_Maori-1
S_Dai-2
S_Brahmin-2
S_Brahui-1
S_Balochi-2
S_Samaritan-1
S_Mongola-1
S_Relli-1
S_Papuan-2
S_Papuan-3
S_Naxi-1
S_Mala-2
S_Kapu-2
S_Yadava-1
S_Yemenite_Jew-2
S_Ami-2
S_Even-2
B_Han-3
S_Palestinian-1
S_North_Ossetian-1
S_Chechen-1
S_Bulgarian-1
S_Finnish-3
S_Jordanian-2
S_Basque-1
S_Georgian-1
S_Hazara-1
S_Kyrgyz-1
S_She-2
S_English-1
S_Tu-1
S_Xibo-2
S_Atayal-1
S_Eskimo_Chaplin-1
S_Eskimo_Sireniki-1
S_Tlingit-1
S_Kusunda-1
S_Ami-1
S_Korean-1
S_Kalash-1
B_Karitiana-3
S_Zapotec-1
S_Papuan-12
S_Tujia-1
B_Dai-4
S_Quechua-2
S_Abkhasian-1
S_Yakut-2
S_Japanese-1
S_Japanese-3
Africa
WestEurasia
CentralAsiaSiberia
SouthAsia
Oceania
EastAsia
America
−0.00030
−0.00015
0.00000
37.2
37.3
37.4
37.5
37.6
37.7

G

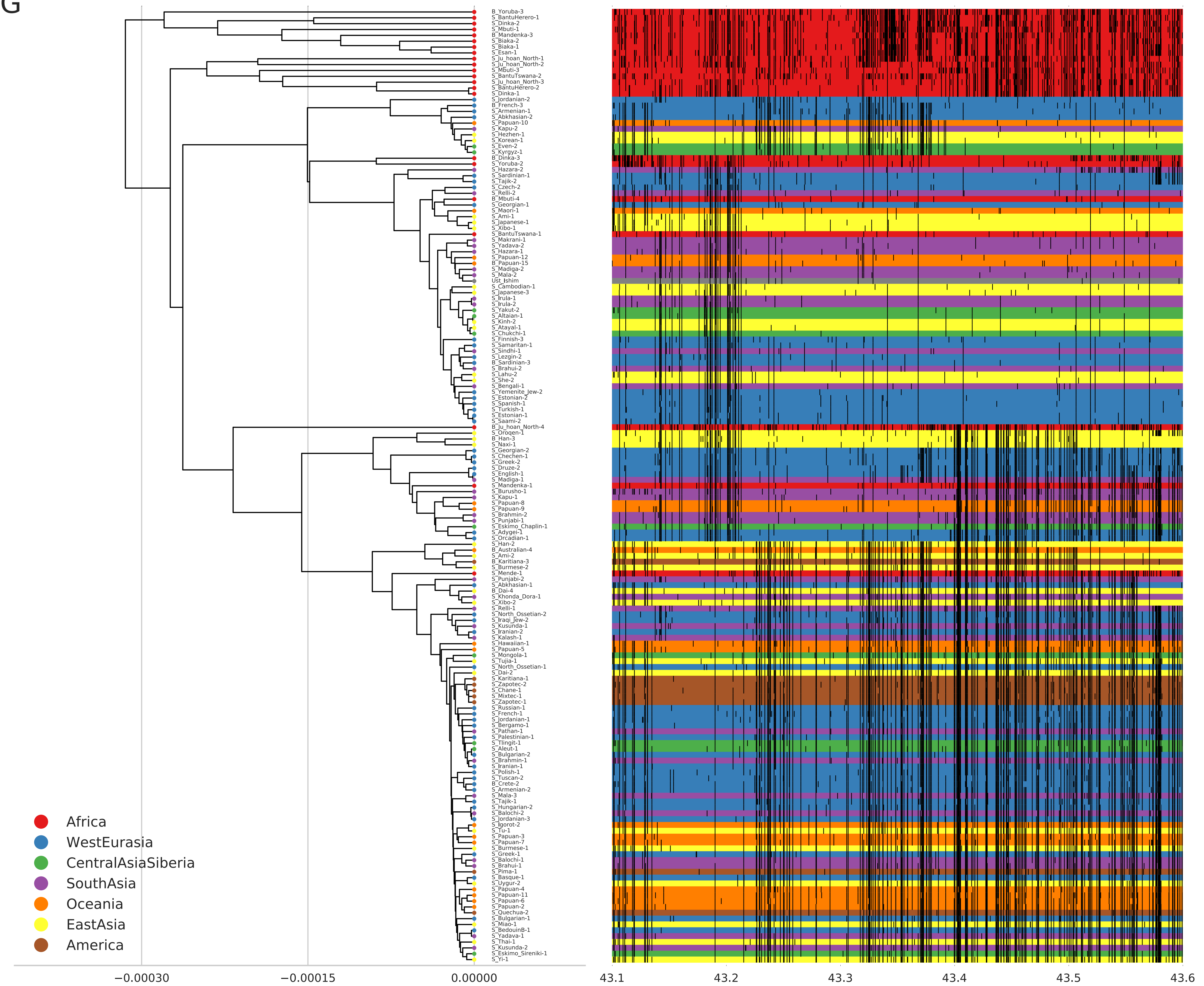

B_Yoruba-3
S_BantuHerero-1
S_Dinka-2
S_Mbuti-1
B_Mandenka-3
S_Biaka-2
S_Biaka-1
S_Esan-1
S_Ju_hoan_North-1
S_Ju_hoan_North-2
S_Mbuti-3
S_BantuTswana-2
S_Ju_hoan_North-3
S_BantuHerero-2
S_Dinka-1
S_Jordanian-2
B_French-3
S_Armenian-1
S_Abkhasian-2
S_Papuan-10
S_Kapu-2
S_Hezhen-1
S_Korean-1
S_Even-2
S_Kyrgyz-1
B_Dinka-3
S_Yoruba-2
S_Hazara-2
S_Sardinian-1
S_Tajik-2
S_Czech-2
S_Relli-2
B_Mbuti-4
S_Georgian-1
S_Maori-1
S_Ami-1
S_Japanese-1
S_Xibo-1
S_BantuTswana-1
S_Makrani-1
S_Yadava-2
S_Hazara-1
S_Papuan-12
B_Papuan-15
S_Madiga-2
S_Mala-2
Ust_Ishim
S_Cambodian-1
S_Japanese-3
S_Irula-1
S_Irula-2
S_Yakut-2
S_Altaian-1
S_Kinh-2
S_Atayal-1
S_Chukchi-1
S_Finnish-3
S_Samaritan-1
S_Sindhi-1
S_Lezgin-2
B_Sardinian-3
S_Brahui-2
S_Lahu-2
S_She-2
S_Bengali-1
S_Yemenite_Jew-2
S_Estonian-2
S_Spanish-1
S_Turkish-1
S_Estonian-1
S_Saami-2
B_Ju_hoan_North-4
S_Oroqen-1
B_Han-3
S_Naxi-1
S_Georgian-2
S_Chechen-1
S_Greek-2
S_Druze-2
S_English-1
S_Madiga-1
S_Mandenka-1
S_Burusho-1
S_Kapu-1
S_Papuan-8
S_Papuan-9
S_Brahmin-2
S_Punjabi-1
S_Eskimo_Chaplin-1
S_Adygei-1
S_Orcadian-1
S_Han-2
B_Australian-4
S_Ami-2
B_Karitiana-3
S_Burmese-2
S_Mende-1
S_Punjabi-2
S_Abkhasian-1
B_Dai-4
S_Khonda_Dora-1
S_Xibo-2
S_Relli-1
S_North_Ossetian-2
S_Iraqi_Jew-2
S_Kusunda-1
S_Iranian-2
S_Kalash-1
S_Hawaiian-1
S_Papuan-5
S_Mongola-1
S_Tujia-1
S_North_Ossetian-1
S_Dai-2
S_Karitiana-1
S_Zapotec-2
S_Chane-1
S_Mixtec-1
S_Zapotec-1
S_Russian-1
S_French-1
S_Jordanian-1
S_Bergamo-1
S_Pathan-1
S_Palestinian-1
S_Tlingit-1
S_Aleut-1
S_Bulgarian-2
S_Brahmin-1
S_Iranian-1
S_Polish-1
S_Tuscan-2
B_Crete-2
S_Armenian-2
S_Mala-3
S_Tajik-1
S_Hungarian-2
S_Balochi-2
S_Jordanian-3
S_Igorot-2
S_Tu-1
S_Papuan-3
S_Papuan-7
S_Burmese-1
S_Greek-1
S_Balochi-1
S_Brahui-1
S_Pima-1
S_Basque-1
S_Uygur-2
S_Papuan-4
S_Papuan-11
S_Papuan-6
S_Papuan-2
S_Quechua-2
S_Bulgarian-1
S_Miao-1
S_BedouinB-1
S_Yadava-1
S_Thai-1
S_Kusunda-2
S_Eskimo_Sireniki-1
S_Yi-1
Africa
WestEurasia
CentralAsiaSiberia
SouthAsia
Oceania
EastAsia
America
−0.00030
−0.00015
0.00000
43.1
43.2
43.3
43.4
43.5
43.6

H

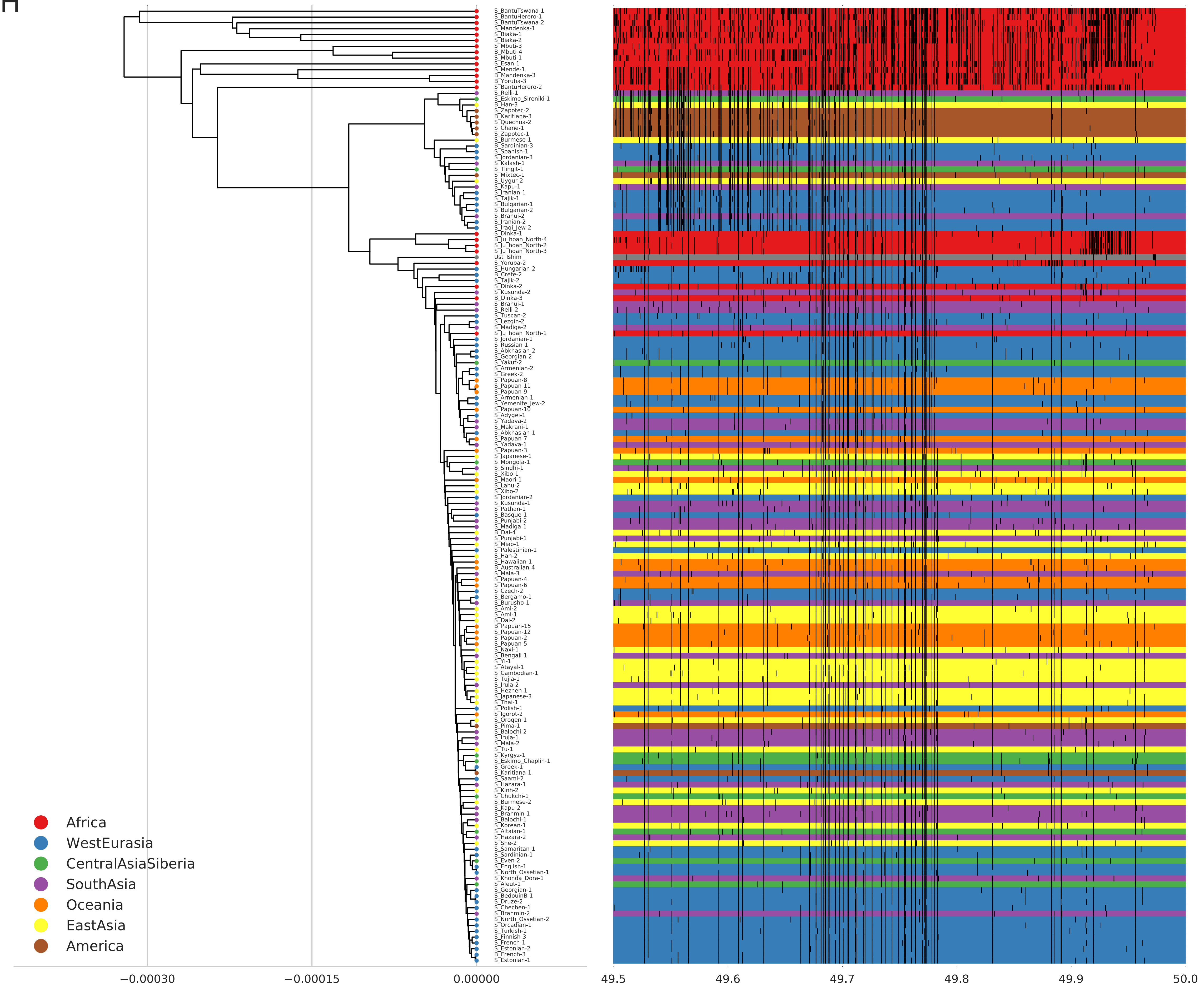

S_BantuTswana-1
S_BantuHerero-1
S_BantuTswana-2
S_Mandenka-1
S_Biaka-1
S_Biaka-2
S_Mbuti-3
B_Mbuti-4
S_Mbuti-1
S_Esan-1
S_Mende-1
B_Mandenka-3
B_Yoruba-3
S_BantuHerero-2
S_Relli-1
S_Eskimo_Sireniki-1
B_Han-3
S_Zapotec-2
B_Karitiana-3
S_Quechua-2
S_Chane-1
S_Zapotec-1
S_Burmese-1
B_Sardinian-3
S_Spanish-1
S_Jordanian-3
S_Kalash-1
S_Tlingit-1
S_Mixtec-1
S_Uygur-2
S_Kapu-1
S_Iranian-1
S_Tajik-1
S_Bulgarian-1
S_Bulgarian-2
S_Brahui-2
S_Iranian-2
S_Iraqi_Jew-2
S_Dinka-1
B_Ju_hoan_North-4
S_Ju_hoan_North-2
S_Ju_hoan_North-3
Ust_Ishim
S_Yoruba-2
S_Hungarian-2
B_Crete-2
S_Tajik-2
S_Dinka-2
S_Kusunda-2
B_Dinka-3
S_Brahui-1
S_Relli-2
S_Tuscan-2
S_Lezgin-2
S_Madiga-2
S_Ju_hoan_North-1
S_Jordanian-1
S_Russian-1
S_Abkhasian-2
S_Georgian-2
S_Yakut-2
S_Armenian-2
S_Greek-2
S_Papuan-8
S_Papuan-11
S_Papuan-9
S_Armenian-1
S_Yemenite_Jew-2
S_Papuan-10
S_Adygei-1
S_Yadava-2
S_Makrani-1
S_Abkhasian-1
S_Papuan-7
S_Yadava-1
S_Papuan-3
S_Japanese-1
S_Mongola-1
S_Sindhi-1
S_Xibo-1
S_Maori-1
S_Lahu-2
S_Xibo-2
S_Jordanian-2
S_Kusunda-1
S_Pathan-1
S_Basque-1
S_Punjabi-2
S_Madiga-1
B_Dai-4
S_Punjabi-1
S_Miao-1
S_Palestinian-1
S_Han-2
S_Hawaiian-1
B_Australian-4
S_Mala-3
S_Papuan-4
S_Papuan-6
S_Czech-2
S_Bergamo-1
S_Burusho-1
S_Ami-2
S_Ami-1
S_Dai-2
B_Papuan-15
S_Papuan-12
S_Papuan-2
S_Papuan-5
S_Naxi-1
S_Bengali-1
S_Yi-1
S_Atayal-1
S_Cambodian-1
S_Tujia-1
S_Irula-2
S_Hezhen-1
S_Japanese-3
S_Thai-1
S_Polish-1
S_Igorot-2
S_Oroqen-1
S_Pima-1
S_Balochi-2
S_Irula-1
S_Mala-2
S_Tu-1
S_Kyrgyz-1
S_Eskimo_Chaplin-1
S_Greek-1
S_Karitiana-1
S_Saami-2
S_Hazara-1
S_Kinh-2
S_Chukchi-1
S_Burmese-2
S_Kapu-2
S_Brahmin-1
S_Balochi-1
S_Korean-1
S_Altaian-1
S_Hazara-2
S_She-2
S_Samaritan-1
S_Sardinian-1
S_Even-2
S_English-1
S_North_Ossetian-1
S_Khonda_Dora-1
S_Aleut-1
S_Georgian-1
S_BedouinB-1
S_Druze-2
S_Chechen-1
S_Brahmin-2
S_North_Ossetian-2
S_Orcadian-1
S_Turkish-1
S_Finnish-3
S_French-1
S_Estonian-2
B_French-3
S_Estonian-1
Africa
WestEurasia
CentralAsiaSiberia
SouthAsia
Oceania
EastAsia
America
−0.00030
−0.00015
0.00000
49.5
49.6
49.7
49.8
49.9
50.0

I

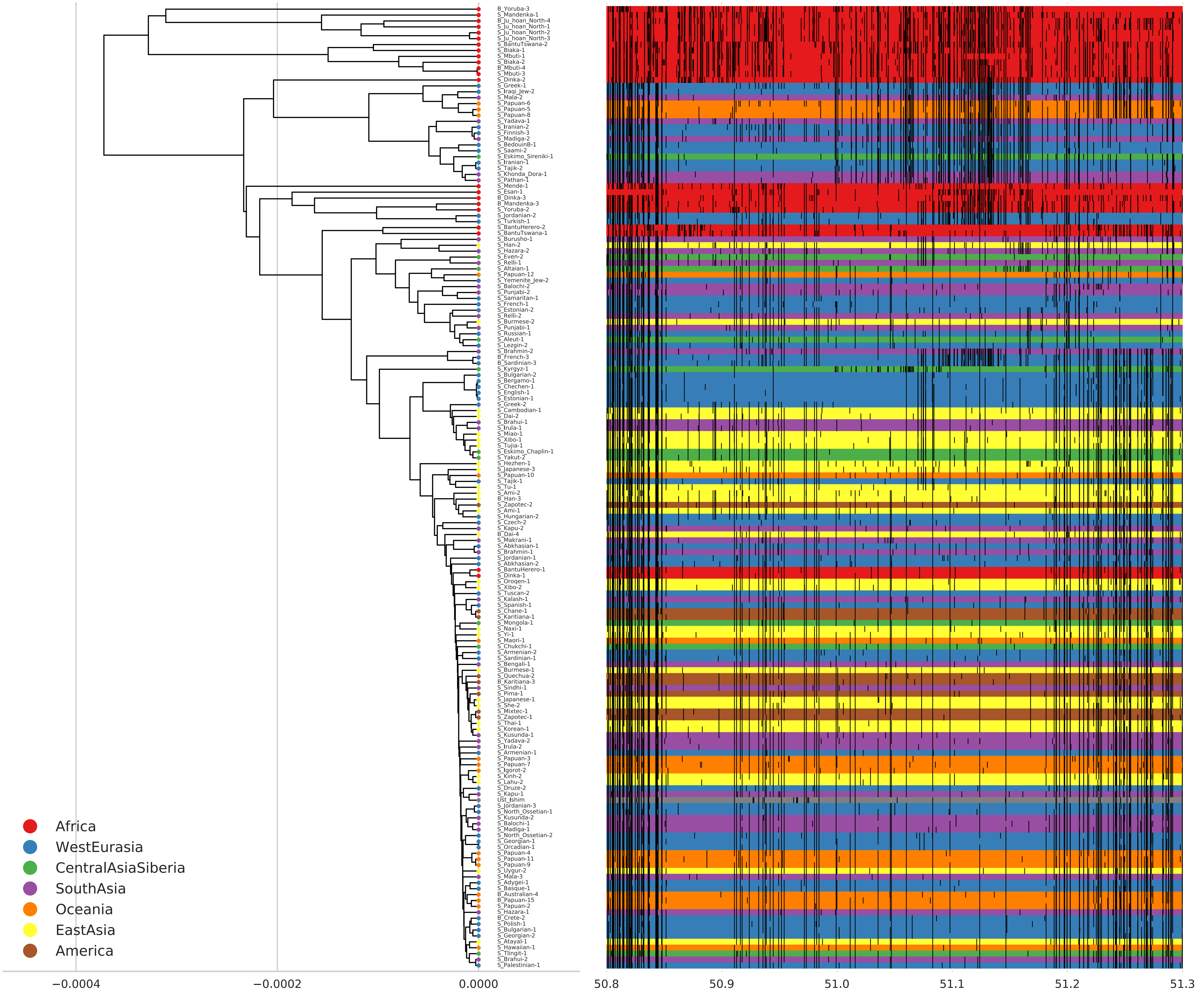

B_Yoruba-3
S_Mandenka-1
B_Ju_hoan_North-4
S_Ju_hoan_North-1
S_Ju_hoan_North-2
S_Ju_hoan_North-3
S_BantuTswana-2
S_Biaka-1
S_Mbuti-1
S_Biaka-2
B_Mbuti-4
S_Mbuti-3
S_Dinka-2
S_Greek-1
S_Iraqi_Jew-2
S_Mala-2
S_Papuan-6
S_Papuan-5
S_Papuan-8
S_Yadava-1
S_Iranian-2
S_Finnish-3
S_Madiga-2
S_BedouinB-1
S_Saami-2
S_Eskimo_Sireniki-1
S_Iranian-1
S_Tajik-2
S_Khonda_Dora-1
S_Pathan-1
S_Mende-1
S_Esan-1
B_Dinka-3
B_Mandenka-3
S_Yoruba-2
S_Jordanian-2
S_Turkish-1
S_BantuHerero-2
S_BantuTswana-1
S_Burusho-1
S_Han-2
S_Hazara-2
S_Even-2
S_Relli-1
S_Altaian-1
S_Papuan-12
S_Yemenite_Jew-2
S_Balochi-2
S_Punjabi-2
S_Samaritan-1
S_French-1
S_Estonian-2
S_Relli-2
S_Burmese-2
S_Punjabi-1
S_Russian-1
S_Aleut-1
S_Lezgin-2
S_Brahmin-2
B_French-3
B_Sardinian-3
S_Kyrgyz-1
S_Bulgarian-2
S_Bergamo-1
S_Chechen-1
S_English-1
S_Estonian-1
S_Greek-2
S_Cambodian-1
S_Dai-2
S_Brahui-1
S_Irula-1
S_Miao-1
S_Xibo-1
S_Tujia-1
S_Eskimo_Chaplin-1
S_Yakut-2
S_Hezhen-1
S_Japanese-3
S_Papuan-10
S_Tajik-1
S_Tu-1
S_Ami-2
B_Han-3
S_Zapotec-2
S_Ami-1
S_Hungarian-2
S_Czech-2
S_Kapu-2
B_Dai-4
S_Makrani-1
S_Abkhasian-1
S_Brahmin-1
S_Jordanian-1
S_Abkhasian-2
S_BantuHerero-1
S_Dinka-1
S_Oroqen-1
S_Xibo-2
S_Tuscan-2
S_Kalash-1
S_Spanish-1
S_Chane-1
S_Karitiana-1
S_Mongola-1
S_Naxi-1
S_Yi-1
S_Maori-1
S_Chukchi-1
S_Armenian-2
S_Sardinian-1
S_Bengali-1
S_Burmese-1
S_Quechua-2
B_Karitiana-3
S_Sindhi-1
S_Pima-1
S_Japanese-1
S_She-2
S_Mixtec-1
S_Zapotec-1
S_Thai-1
S_Korean-1
S_Kusunda-1
S_Yadava-2
S_Irula-2
S_Armenian-1
S_Papuan-3
S_Papuan-7
S_Igorot-2
S_Kinh-2
S_Lahu-2
S_Druze-2
S_Kapu-1
Ust_Ishim
S_Jordanian-3
S_North_Ossetian-1
S_Kusunda-2
S_Balochi-1
S_Madiga-1
S_North_Ossetian-2
S_Georgian-1
S_Orcadian-1
S_Papuan-4
S_Papuan-11
S_Papuan-9
S_Uygur-2
S_Mala-3
S_Adygei-1
S_Basque-1
B_Australian-4
B_Papuan-15
S_Papuan-2
S_Hazara-1
B_Crete-2
S_Polish-1
S_Bulgarian-1
S_Georgian-2
S_Atayal-1
S_Hawaiian-1
S_Tlingit-1
S_Brahui-2
S_Palestinian-1
Africa
WestEurasia
CentralAsiaSiberia
SouthAsia
Oceania
EastAsia
America
−0.0004
−0.0002
0.0000
50.8
50.9
51.0
51.1
51.2
51.3

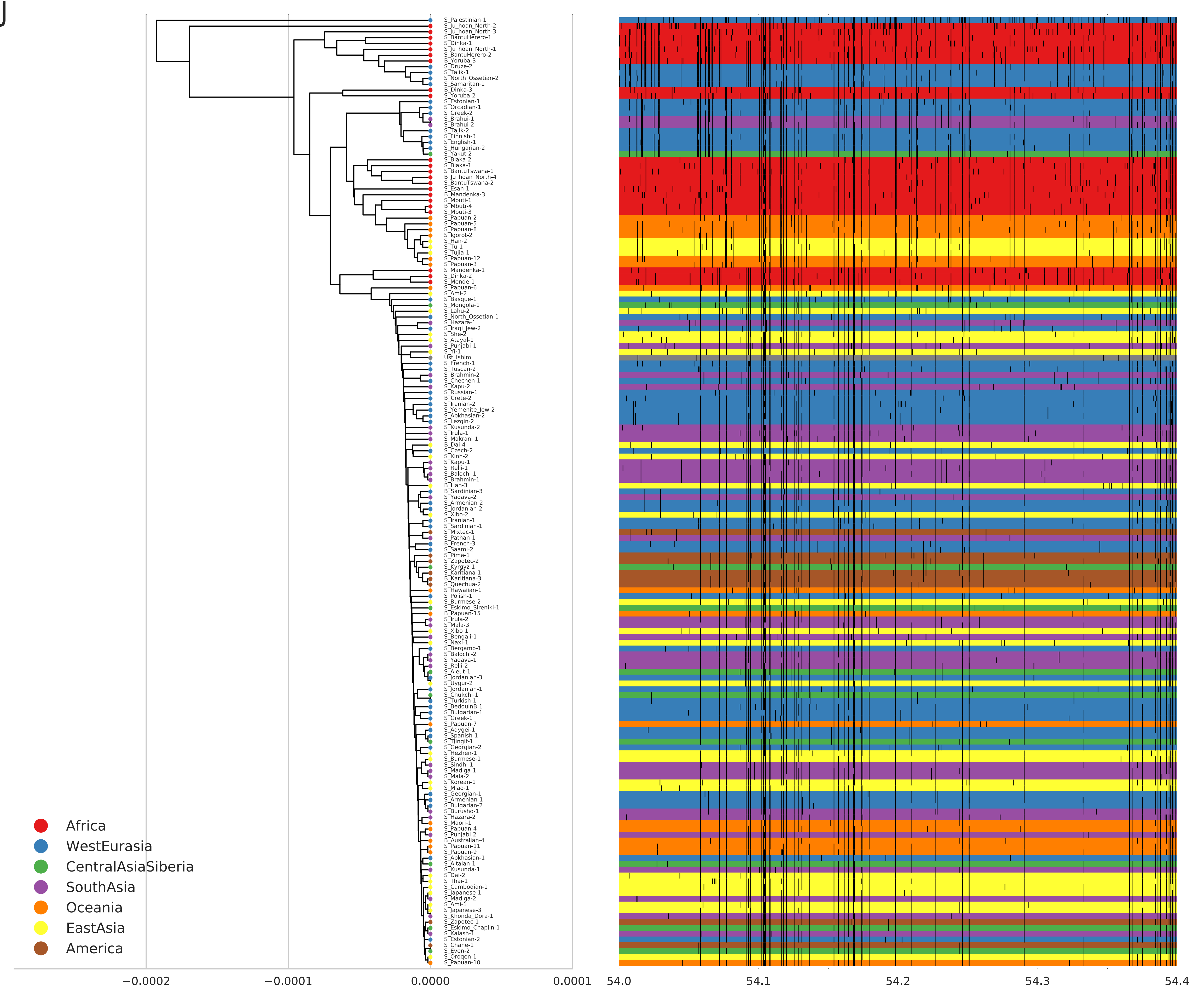
J
Africa
WestEurasia
CentralAsiaSiberia
SouthAsia
Oceania
EastAsia
America
−0.0002
−0.0001
0.0000
0.0001
54.0
54.1
54.2
54.3
54.4

K

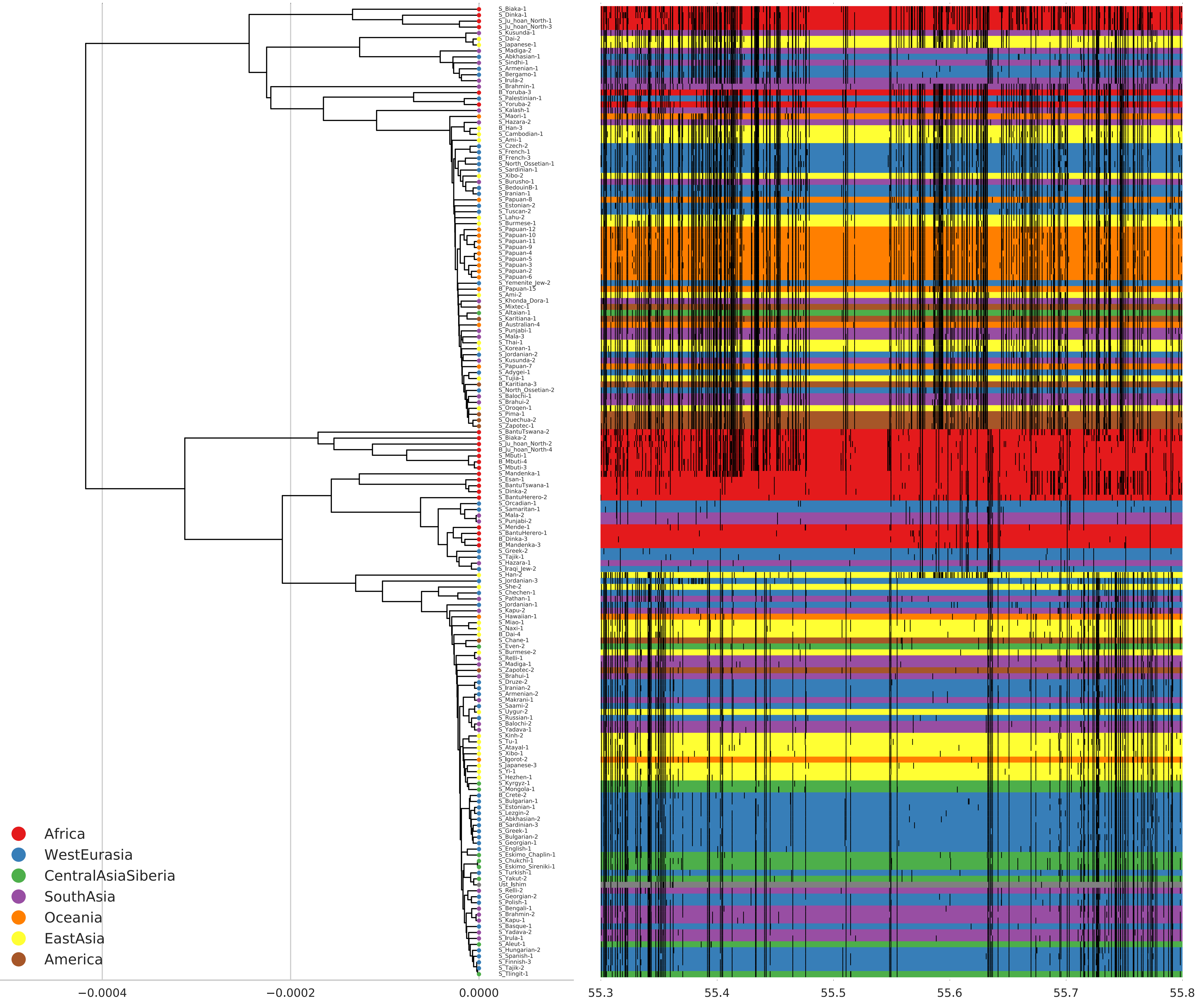

S_Biaka-1
S_Dinka-1
S_Ju_hoan_North-1
S_Ju_hoan_North-3
S_Kusunda-1
S_Dai-2
S_Japanese-1
S_Madiga-2
S_Abkhasian-1
S_Sindhi-1
S_Armenian-1
S_Bergamo-1
S_Irula-2
S_Brahmin-1
B_Yoruba-3
S_Palestinian-1
S_Yoruba-2
S_Kalash-1
S_Maori-1
S_Hazara-2
B_Han-3
S_Cambodian-1
S_Ami-1
S_Czech-2
S_French-1
B_French-3
S_North_Ossetian-1
S_Sardinian-1
S_Xibo-2
S_Burusho-1
S_BedouinB-1
S_Iranian-1
S_Papuan-8
S_Estonian-2
S_Tuscan-2
S_Lahu-2
S_Burmese-1
S_Papuan-12
S_Papuan-10
S_Papuan-11
S_Papuan-9
S_Papuan-4
S_Papuan-5
S_Papuan-3
S_Papuan-2
S_Papuan-6
S_Yemenite_Jew-2
B_Papuan-15
S_Ami-2
S_Khonda_Dora-1
S_Mixtec-1
S_Altaian-1
S_Karitiana-1
B_Australian-4
S_Punjabi-1
S_Mala-3
S_Thai-1
S_Korean-1
S_Jordanian-2
S_Kusunda-2
S_Papuan-7
S_Adygei-1
S_Tujia-1
B_Karitiana-3
S_North_Ossetian-2
S_Balochi-1
S_Brahui-2
S_Oroqen-1
S_Pima-1
S_Quechua-2
S_Zapotec-1
S_BantuTswana-2
S_Biaka-2
S_Ju_hoan_North-2
B_Ju_hoan_North-4
S_Mbuti-1
B_Mbuti-4
S_Mbuti-3
S_Mandenka-1
S_Esan-1
S_BantuTswana-1
S_Dinka-2
S_BantuHerero-2
S_Orcadian-1
S_Samaritan-1
S_Mala-2
S_Punjabi-2
S_Mende-1
S_BantuHerero-1
B_Dinka-3
B_Mandenka-3
S_Greek-2
S_Tajik-1
S_Hazara-1
S_Iraqi_Jew-2
S_Han-2
S_Jordanian-3
S_She-2
S_Chechen-1
S_Pathan-1
S_Jordanian-1
S_Kapu-2
S_Hawaiian-1
S_Miao-1
S_Naxi-1
B_Dai-4
S_Chane-1
S_Even-2
S_Burmese-2
S_Relli-1
S_Madiga-1
S_Zapotec-2
S_Brahui-1
S_Druze-2
S_Iranian-2
S_Armenian-2
S_Makrani-1
S_Saami-2
S_Uygur-2
S_Russian-1
S_Balochi-2
S_Yadava-1
S_Kinh-2
S_Tu-1
S_Atayal-1
S_Xibo-1
S_Igorot-2
S_Japanese-3
S_Yi-1
S_Hezhen-1
S_Kyrgyz-1
S_Mongola-1
B_Crete-2
S_Bulgarian-1
S_Estonian-1
S_Lezgin-2
S_Abkhasian-2
B_Sardinian-3
S_Greek-1
S_Bulgarian-2
S_Georgian-1
S_English-1
S_Eskimo_Chaplin-1
S_Chukchi-1
S_Eskimo_Sireniki-1
S_Turkish-1
S_Yakut-2
Ust_Ishim
S_Relli-2
S_Georgian-2
S_Polish-1
S_Bengali-1
S_Brahmin-2
S_Kapu-1
S_Basque-1
S_Yadava-2
S_Irula-1
S_Aleut-1
S_Hungarian-2
S_Spanish-1
S_Finnish-3
S_Tajik-2
S_Tlingit-1
Africa
WestEurasia
CentralAsiaSiberia
SouthAsia
Oceania
EastAsia
America
−0.0004
−0.0002
0.0000
55.3
55.4
55.5
55.6
55.7
55.8

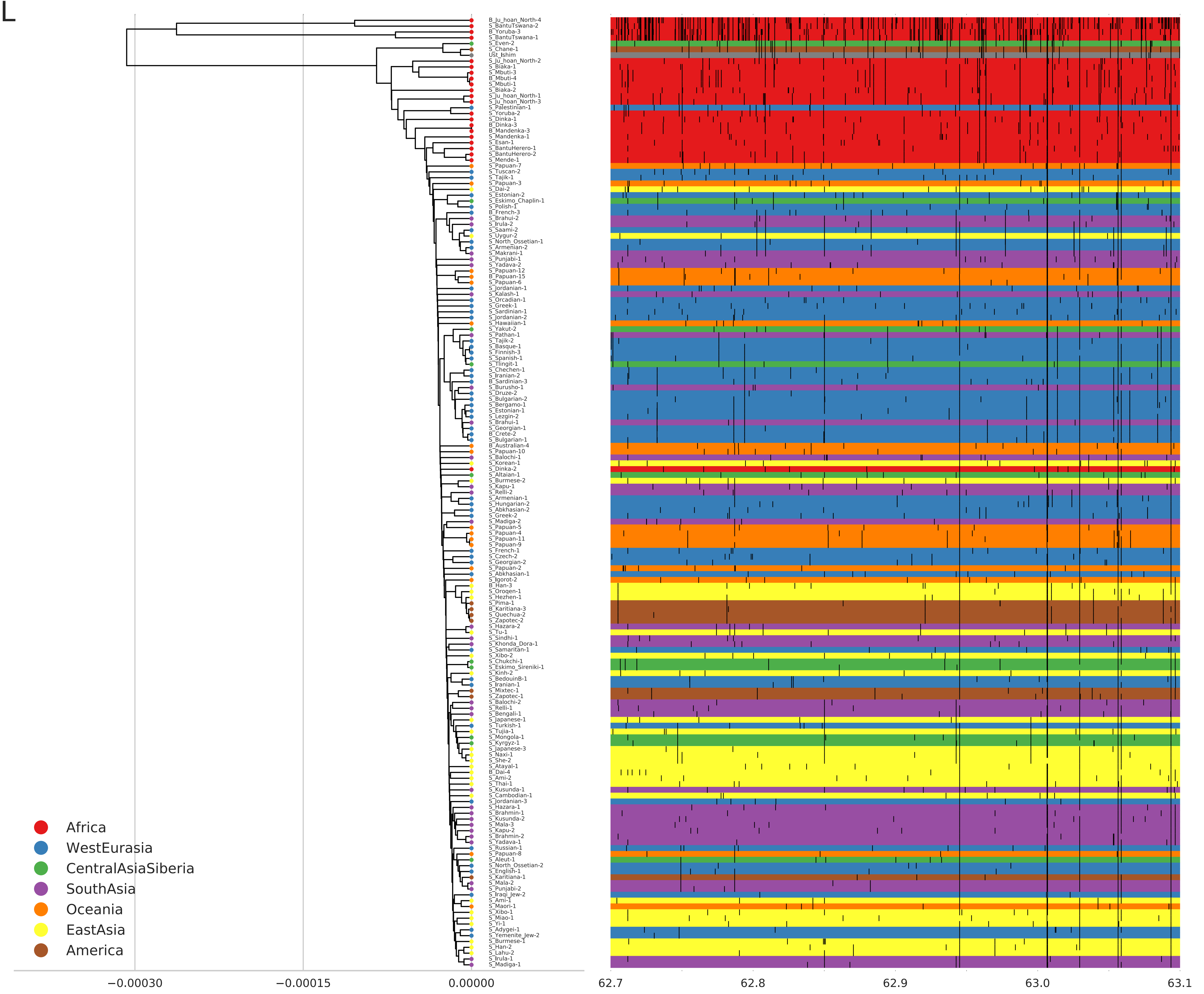
L
B_Ju_hoan_North-4
S_BantuTswana-2
B_Yoruba-3
S_BantuTswana-1
S_Even-2
S_Chane-1
Ust_Ishim
S_Ju_hoan_North-2
S_Biaka-1
S_Mbuti-3
B_Mbuti-4
S_Mbuti-1
S_Biaka-2
S_Ju_hoan_North-1
S_Ju_hoan_North-3
S_Palestinian-1
S_Yoruba-2
S_Dinka-1
B_Dinka-3
B_Mandenka-3
S_Mandenka-1
S_Esan-1
S_BantuHerero-1
S_BantuHerero-2
S_Mende-1
S_Papuan-7
S_Tuscan-2
S_Tajik-1
S_Papuan-3
S_Dai-2
S_Estonian-2
S_Eskimo_Chaplin-1
S_Polish-1
B_French-3
S_Brahui-2
S_Irula-2
S_Saami-2
S_Uygur-2
S_North_Ossetian-1
S_Armenian-2
S_Makrani-1
S_Punjabi-1
S_Yadava-2
S_Papuan-12
B_Papuan-15
S_Papuan-6
S_Jordanian-1
S_Kalash-1
S_Orcadian-1
S_Greek-1
S_Sardinian-1
S_Jordanian-2
S_Hawaiian-1
S_Yakut-2
S_Pathan-1
S_Tajik-2
S_Basque-1
S_Finnish-3
S_Spanish-1
S_Tlingit-1
S_Chechen-1
S_Iranian-2
B_Sardinian-3
S_Burusho-1
S_Druze-2
S_Bulgarian-2
S_Bergamo-1
S_Estonian-1
S_Lezgin-2
S_Brahui-1
S_Georgian-1
B_Crete-2
S_Bulgarian-1
B_Australian-4
S_Papuan-10
S_Balochi-1
S_Korean-1
S_Dinka-2
S_Altaian-1
S_Burmese-2
S_Kapu-1
S_Relli-2
S_Armenian-1
S_Hungarian-2
S_Abkhasian-2
S_Greek-2
S_Madiga-2
S_Papuan-5
S_Papuan-4
S_Papuan-11
S_Papuan-9
S_French-1
S_Czech-2
S_Georgian-2
S_Papuan-2
S_Abkhasian-1
S_Igorot-2
B_Han-3
S_Oroqen-1
S_Hezhen-1
S_Pima-1
B_Karitiana-3
S_Quechua-2
S_Zapotec-2
S_Hazara-2
S_Tu-1
S_Sindhi-1
S_Khonda_Dora-1
S_Samaritan-1
S_Xibo-2
S_Chukchi-1
S_Eskimo_Sireniki-1
S_Kinh-2
S_BedouinB-1
S_Iranian-1
S_Mixtec-1
S_Zapotec-1
S_Balochi-2
S_Relli-1
S_Bengali-1
S_Japanese-1
S_Turkish-1
S_Tujia-1
S_Mongola-1
S_Kyrgyz-1
S_Japanese-3
S_Naxi-1
S_She-2
S_Atayal-1
B_Dai-4
S_Ami-2
S_Thai-1
S_Kusunda-1
S_Cambodian-1
S_Jordanian-3
S_Hazara-1
S_Brahmin-1
S_Kusunda-2
S_Mala-3
S_Kapu-2
S_Brahmin-2
S_Yadava-1
S_Russian-1
S_Papuan-8
S_Aleut-1
S_North_Ossetian-2
S_English-1
S_Karitiana-1
S_Mala-2
S_Punjabi-2
S_Iraqi_Jew-2
S_Ami-1
S_Maori-1
S_Xibo-1
S_Miao-1
S_Yi-1
S_Adygei-1
S_Yemenite_Jew-2
S_Burmese-1
S_Han-2
S_Lahu-2
S_Irula-1
S_Madiga-1
Africa
WestEurasia
CentralAsiaSiberia
SouthAsia
Oceania
EastAsia
America
−0.00030
−0.00015
0.00000
62.7
62.8
62.9
63.0
63.1

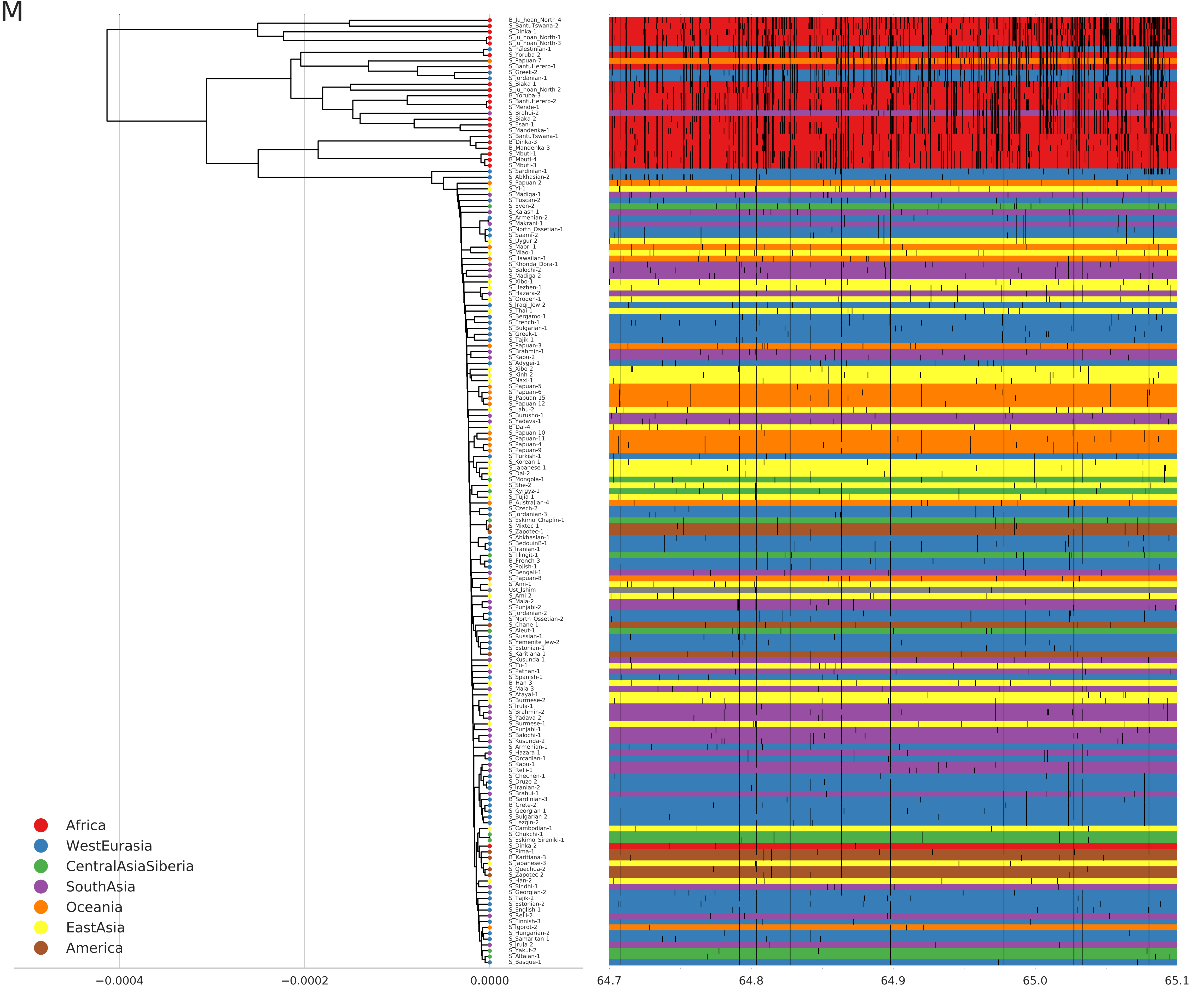
M
B_Ju_hoan_North-4
S_BantuTswana-2
S_Dinka-1
S_Ju_hoan_North-1
S_Ju_hoan_North-3
S_Palestinian-1
S_Yoruba-2
S_Papuan-7
S_BantuHerero-1
S_Greek-2
S_Jordanian-1
S_Biaka-1
S_Ju_hoan_North-2
B_Yoruba-3
S_BantuHerero-2
S_Mende-1
S_Brahui-2
S_Biaka-2
S_Esan-1
S_Mandenka-1
S_BantuTswana-1
B_Dinka-3
B_Mandenka-3
S_Mbuti-1
B_Mbuti-4
S_Mbuti-3
S_Sardinian-1
S_Abkhasian-2
S_Papuan-2
S_Yi-1
S_Madiga-1
S_Tuscan-2
S_Even-2
S_Kalash-1
S_Armenian-2
S_Makrani-1
S_North_Ossetian-1
S_Saami-2
S_Uygur-2
S_Maori-1
S_Miao-1
S_Hawaiian-1
S_Khonda_Dora-1
S_Balochi-2
S_Madiga-2
S_Xibo-1
S_Hezhen-1
S_Hazara-2
S_Oroqen-1
S_Iraqi_Jew-2
S_Thai-1
S_Bergamo-1
S_French-1
S_Bulgarian-1
S_Greek-1
S_Tajik-1
S_Papuan-3
S_Brahmin-1
S_Kapu-2
S_Adygei-1
S_Xibo-2
S_Kinh-2
S_Naxi-1
S_Papuan-5
S_Papuan-6
B_Papuan-15
S_Papuan-12
S_Lahu-2
S_Burusho-1
S_Yadava-1
B_Dai-4
S_Papuan-10
S_Papuan-11
S_Papuan-4
S_Papuan-9
S_Turkish-1
S_Korean-1
S_Japanese-1
S_Dai-2
S_Mongola-1
S_She-2
S_Kyrgyz-1
S_Tujia-1
B_Australian-4
S_Czech-2
S_Jordanian-3
S_Eskimo_Chaplin-1
S_Mixtec-1
S_Zapotec-1
S_Abkhasian-1
S_BedouinB-1
S_Iranian-1
S_Tlingit-1
B_French-3
S_Polish-1
S_Bengali-1
S_Papuan-8
S_Ami-1
Ust_Ishim
S_Ami-2
S_Mala-2
S_Punjabi-2
S_Jordanian-2
S_North_Ossetian-2
S_Chane-1
S_Aleut-1
S_Russian-1
S_Yemenite_Jew-2
S_Estonian-1
S_Karitiana-1
S_Kusunda-1
S_Tu-1
S_Pathan-1
S_Spanish-1
B_Han-3
S_Mala-3
S_Atayal-1
S_Burmese-2
S_Irula-1
S_Brahmin-2
S_Yadava-2
S_Burmese-1
S_Punjabi-1
S_Balochi-1
S_Kusunda-2
S_Armenian-1
S_Hazara-1
S_Orcadian-1
S_Kapu-1
S_Relli-1
S_Chechen-1
S_Druze-2
S_Iranian-2
S_Brahui-1
B_Sardinian-3
B_Crete-2
S_Georgian-1
S_Bulgarian-2
S_Lezgin-2
S_Cambodian-1
S_Chukchi-1
S_Eskimo_Sireniki-1
S_Dinka-2
S_Pima-1
B_Karitiana-3
S_Japanese-3
S_Quechua-2
S_Zapotec-2
S_Han-2
S_Sindhi-1
S_Georgian-2
S_Tajik-2
S_Estonian-2
S_English-1
S_Relli-2
S_Finnish-3
S_Igorot-2
S_Hungarian-2
S_Samaritan-1
S_Irula-2
S_Yakut-2
S_Altaian-1
S_Basque-1
Africa
WestEurasia
CentralAsiaSiberia
SouthAsia
Oceania
EastAsia
America
−0.0004
−0.0002
0.0000
64.7
64.8
64.9
65.0
65.1

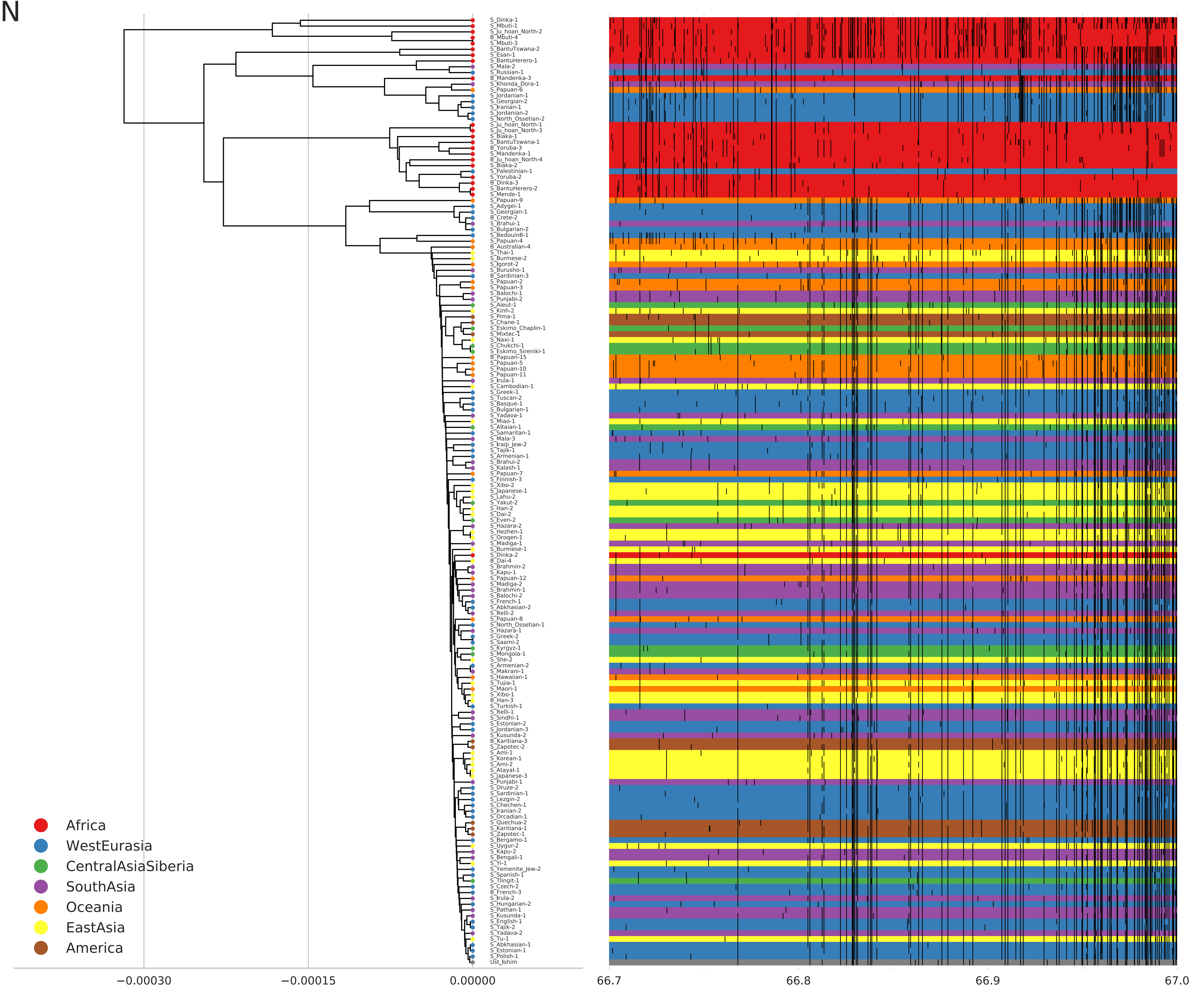
N
S_Dinka-1
S_Mbuti-1
S_Ju_hoan_North-2
B_Mbuti-4
S_Mbuti-3
S_BantuTswana-2
S_Esan-1
S_BantuHerero-1
S_Mala-2
S_Russian-1
B_Mandenka-3
S_Khonda_Dora-1
S_Papuan-6
S_Jordanian-1
S_Georgian-2
S_Iranian-1
S_Jordanian-2
S_North_Ossetian-2
S_Ju_hoan_North-1
S_Ju_hoan_North-3
S_Biaka-1
S_BantuTswana-1
B_Yoruba-3
S_Mandenka-1
B_Ju_hoan_North-4
S_Biaka-2
S_Palestinian-1
S_Yoruba-2
B_Dinka-3
S_BantuHerero-2
S_Mende-1
S_Papuan-9
S_Adygei-1
S_Georgian-1
B_Crete-2
S_Brahui-1
S_Bulgarian-2
S_BedouinB-1
S_Papuan-4
B_Australian-4
S_Thai-1
S_Burmese-2
S_Igorot-2
S_Burusho-1
B_Sardinian-3
S_Papuan-2
S_Papuan-3
S_Balochi-1
S_Punjabi-2
S_Aleut-1
S_Kinh-2
S_Pima-1
S_Chane-1
S_Eskimo_Chaplin-1
S_Mixtec-1
S_Naxi-1
S_Chukchi-1
S_Eskimo_Sireniki-1
B_Papuan-15
S_Papuan-5
S_Papuan-10
S_Papuan-11
S_Irula-1
S_Cambodian-1
S_Greek-1
S_Tuscan-2
S_Basque-1
S_Bulgarian-1
S_Yadava-1
S_Miao-1
S_Altaian-1
S_Samaritan-1
S_Mala-3
S_Iraqi_Jew-2
S_Tajik-1
S_Armenian-1
S_Brahui-2
S_Kalash-1
S_Papuan-7
S_Finnish-3
S_Xibo-2
S_Japanese-1
S_Lahu-2
S_Yakut-2
S_Han-2
S_Dai-2
S_Even-2
S_Hazara-2
S_Hezhen-1
S_Oroqen-1
S_Madiga-1
S_Burmese-1
S_Dinka-2
B_Dai-4
S_Brahmin-2
S_Kapu-1
S_Papuan-12
S_Madiga-2
S_Brahmin-1
S_Balochi-2
S_French-1
S_Abkhasian-2
S_Relli-2
S_Papuan-8
S_North_Ossetian-1
S_Hazara-1
S_Greek-2
S_Saami-2
S_Kyrgyz-1
S_Mongola-1
S_She-2
S_Armenian-2
S_Makrani-1
S_Hawaiian-1
S_Tujia-1
S_Maori-1
S_Xibo-1
B_Han-3
S_Turkish-1
S_Relli-1
S_Sindhi-1
S_Estonian-2
S_Jordanian-3
S_Kusunda-2
B_Karitiana-3
S_Zapotec-2
S_Ami-1
S_Korean-1
S_Ami-2
S_Atayal-1
S_Japanese-3
S_Punjabi-1
S_Druze-2
S_Sardinian-1
S_Lezgin-2
S_Chechen-1
S_Iranian-2
S_Orcadian-1
S_Quechua-2
S_Karitiana-1
S_Zapotec-1
S_Bergamo-1
S_Uygur-2
S_Kapu-2
S_Bengali-1
S_Yi-1
S_Yemenite_Jew-2
S_Spanish-1
S_Tlingit-1
S_Czech-2
B_French-3
S_Irula-2
S_Hungarian-2
S_Pathan-1
S_Kusunda-1
S_English-1
S_Tajik-2
S_Yadava-2
S_Tu-1
S_Abkhasian-1
S_Estonian-1
S_Polish-1
Ust_Ishim
Africa
WestEurasia
CentralAsiaSiberia
SouthAsia
Oceania
EastAsia
America
−0.00030
−0.00015
0.00000
66.7
66.8
66.9
67.0

O

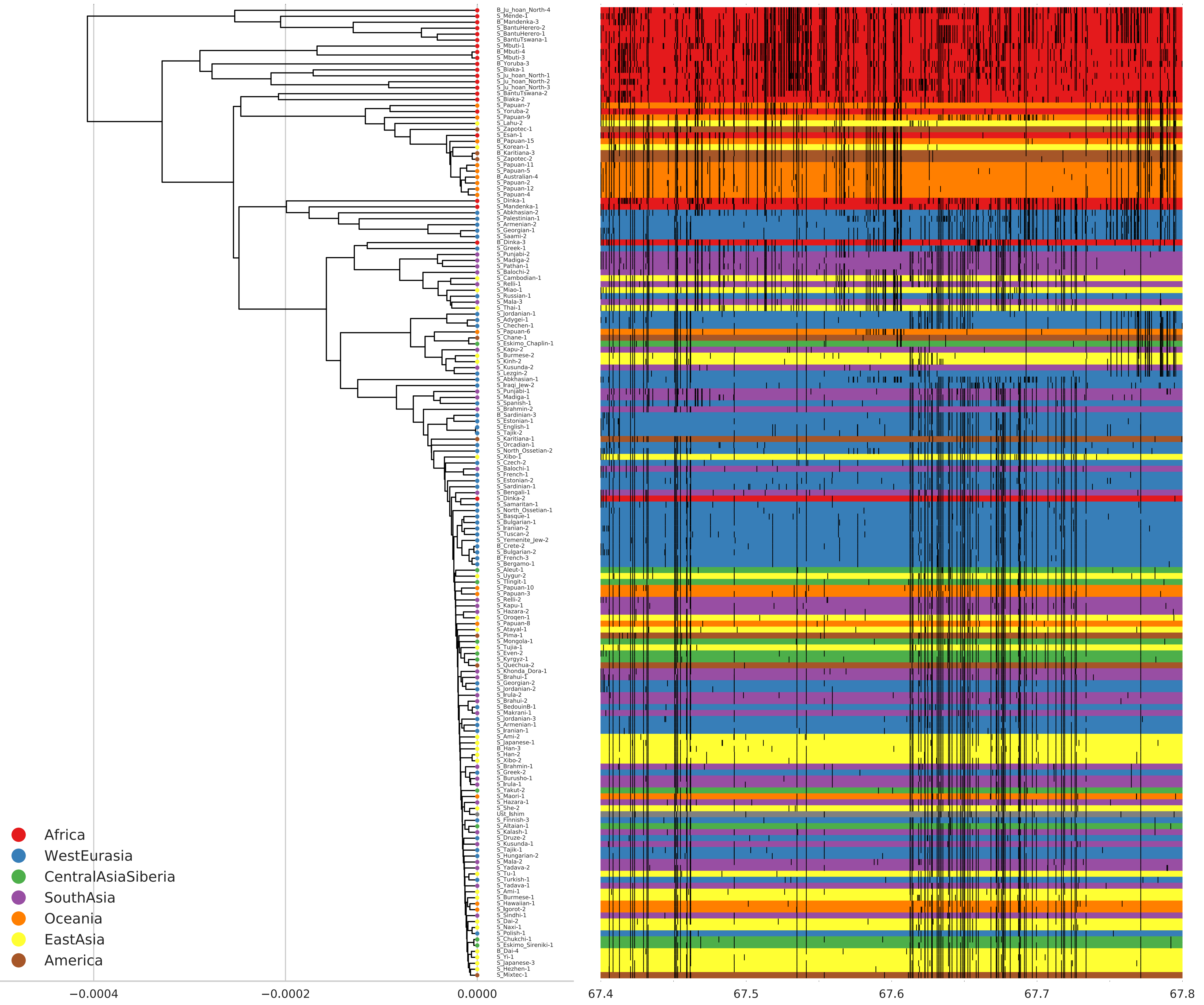
B_Ju_hoan_North-4
S_Mende-1
B_Mandenka-3
S_BantuHerero-2
S_BantuHerero-1
S_BantuTswana-1
S_Mbuti-1
B_Mbuti-4
S_Mbuti-3
B_Yoruba-3
S_Biaka-1
S_Ju_hoan_North-1
S_Ju_hoan_North-2
S_Ju_hoan_North-3
S_BantuTswana-2
S_Biaka-2
S_Papuan-7
S_Yoruba-2
S_Papuan-9
S_Lahu-2
S_Zapotec-1
S_Esan-1
B_Papuan-15
S_Korean-1
B_Karitiana-3
S_Zapotec-2
S_Papuan-11
S_Papuan-5
B_Australian-4
S_Papuan-2
S_Papuan-12
S_Papuan-4
S_Dinka-1
S_Mandenka-1
S_Abkhasian-2
S_Palestinian-1
S_Armenian-2
S_Georgian-1
S_Saami-2
B_Dinka-3
S_Greek-1
S_Punjabi-2
S_Madiga-2
S_Pathan-1
S_Balochi-2
S_Cambodian-1
S_Relli-1
S_Miao-1
S_Russian-1
S_Mala-3
S_Thai-1
S_Jordanian-1
S_Adygei-1
S_Chechen-1
S_Papuan-6
S_Chane-1
S_Eskimo_Chaplin-1
S_Kapu-2
S_Burmese-2
S_Kinh-2
S_Kusunda-2
S_Lezgin-2
S_Abkhasian-1
S_Iraqi_Jew-2
S_Punjabi-1
S_Madiga-1
S_Spanish-1
S_Brahmin-2
B_Sardinian-3
S_Estonian-1
S_English-1
S_Tajik-2
S_Karitiana-1
S_Orcadian-1
S_North_Ossetian-2
S_Xibo-1
S_Czech-2
S_Balochi-1
S_French-1
S_Estonian-2
S_Sardinian-1
S_Bengali-1
S_Dinka-2
S_Samaritan-1
S_North_Ossetian-1
S_Basque-1
S_Bulgarian-1
S_Iranian-2
S_Tuscan-2
S_Yemenite_Jew-2
B_Crete-2
S_Bulgarian-2
B_French-3
S_Bergamo-1
S_Aleut-1
S_Uygur-2
S_Tlingit-1
S_Papuan-10
S_Papuan-3
S_Relli-2
S_Kapu-1
S_Hazara-2
S_Oroqen-1
S_Papuan-8
S_Atayal-1
S_Pima-1
S_Mongola-1
S_Tujia-1
S_Even-2
S_Kyrgyz-1
S_Quechua-2
S_Khonda_Dora-1
S_Brahui-1
S_Georgian-2
S_Jordanian-2
S_Irula-2
S_Brahui-2
S_BedouinB-1
S_Makrani-1
S_Jordanian-3
S_Armenian-1
S_Iranian-1
S_Ami-2
S_Japanese-1
B_Han-3
S_Han-2
S_Xibo-2
S_Brahmin-1
S_Greek-2
S_Burusho-1
S_Irula-1
S_Yakut-2
S_Maori-1
S_Hazara-1
S_She-2
Ust_Ishim
S_Finnish-3
S_Altaian-1
S_Kalash-1
S_Druze-2
S_Kusunda-1
S_Tajik-1
S_Hungarian-2
S_Mala-2
S_Yadava-2
S_Tu-1
S_Turkish-1
S_Yadava-1
S_Ami-1
S_Burmese-1
S_Hawaiian-1
S_Igorot-2
S_Sindhi-1
S_Dai-2
S_Naxi-1
S_Polish-1
S_Chukchi-1
S_Eskimo_Sireniki-1
B_Dai-4
S_Yi-1
S_Japanese-3
S_Hezhen-1
S_Mixtec-1
Africa
WestEurasia
CentralAsiaSiberia
SouthAsia
Oceania
EastAsia
America
−0.0004
−0.0002
0.0000
67.4
67.5
67.6
67.7
67.8

P

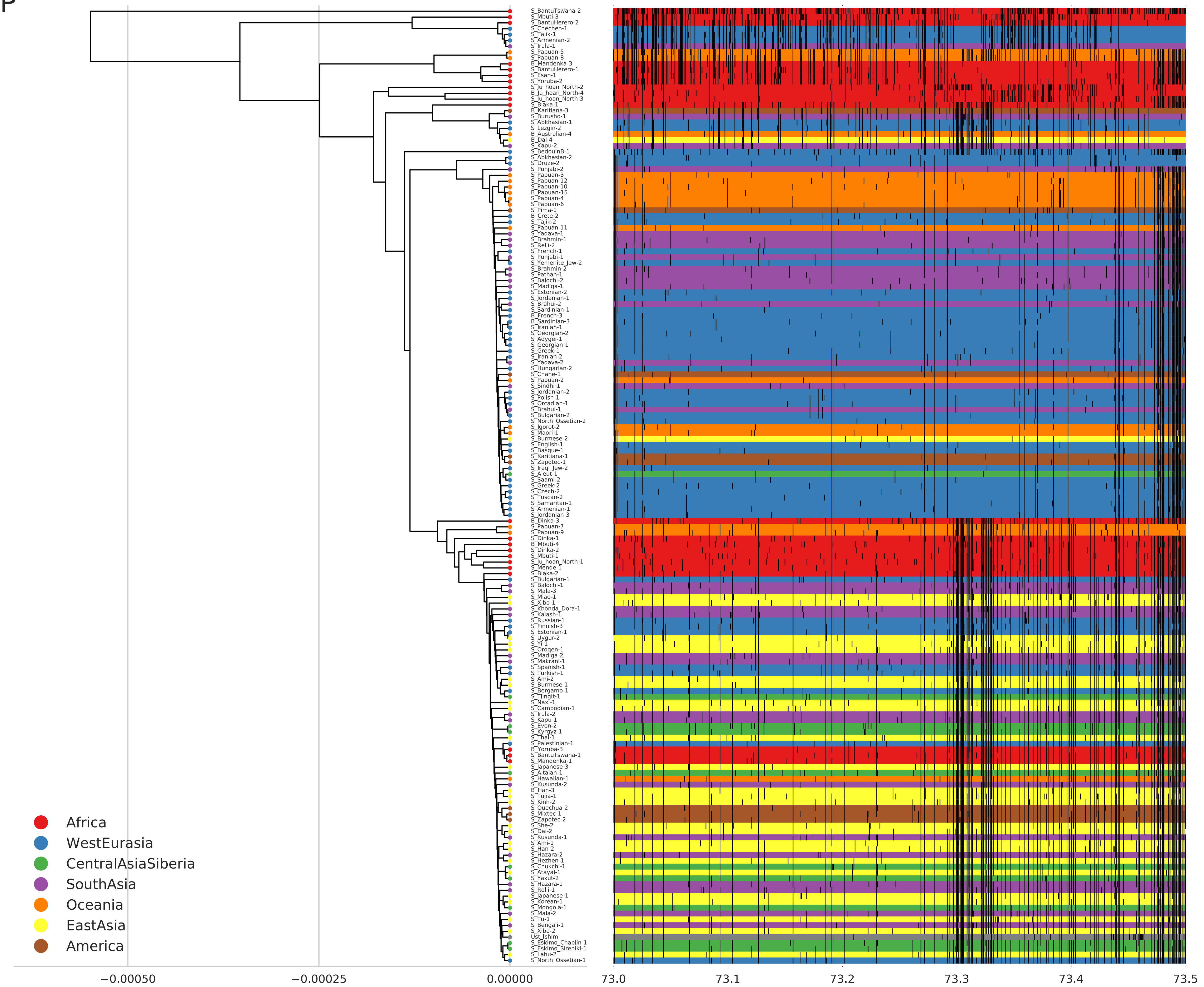

S_BantuTswana-2
S_Mbuti-3
S_BantuHerero-2
S_Chechen-1
S_Tajik-1
S_Armenian-2
S_Irula-1
S_Papuan-5
S_Papuan-8
B_Mandenka-3
S_BantuHerero-1
S_Esan-1
S_Yoruba-2
S_Ju_hoan_North-2
B_Ju_hoan_North-4
S_Ju_hoan_North-3
S_Biaka-1
B_Karitiana-3
S_Burusho-1
S_Abkhasian-1
S_Lezgin-2
B_Australian-4
B_Dai-4
S_Kapu-2
S_BedouinB-1
S_Abkhasian-2
S_Druze-2
S_Punjabi-2
S_Papuan-3
S_Papuan-12
S_Papuan-10
B_Papuan-15
S_Papuan-4
S_Papuan-6
S_Pima-1
B_Crete-2
S_Tajik-2
S_Papuan-11
S_Yadava-1
S_Brahmin-1
S_Relli-2
S_French-1
S_Punjabi-1
S_Yemenite_Jew-2
S_Brahmin-2
S_Pathan-1
S_Balochi-2
S_Madiga-1
S_Estonian-2
S_Jordanian-1
S_Brahui-2
S_Sardinian-1
B_French-3
B_Sardinian-3
S_Iranian-1
S_Georgian-2
S_Adygei-1
S_Georgian-1
S_Greek-1
S_Iranian-2
S_Yadava-2
S_Hungarian-2
S_Chane-1
S_Papuan-2
S_Sindhi-1
S_Jordanian-2
S_Polish-1
S_Orcadian-1
S_Brahui-1
S_Bulgarian-2
S_North_Ossetian-2
S_Igorot-2
S_Maori-1
S_Burmese-2
S_English-1
S_Basque-1
S_Karitiana-1
S_Zapotec-1
S_Iraqi_Jew-2
S_Aleut-1
S_Saami-2
S_Greek-2
S_Czech-2
S_Tuscan-2
S_Samaritan-1
S_Armenian-1
S_Jordanian-3
B_Dinka-3
S_Papuan-7
S_Papuan-9
S_Dinka-1
B_Mbuti-4
S_Dinka-2
S_Mbuti-1
S_Ju_hoan_North-1
S_Mende-1
S_Biaka-2
S_Bulgarian-1
S_Balochi-1
S_Mala-3
S_Miao-1
S_Xibo-1
S_Khonda_Dora-1
S_Kalash-1
S_Russian-1
S_Finnish-3
S_Estonian-1
S_Uygur-2
S_Yi-1
S_Oroqen-1
S_Madiga-2
S_Makrani-1
S_Spanish-1
S_Turkish-1
S_Ami-2
S_Burmese-1
S_Bergamo-1
S_Tlingit-1
S_Naxi-1
S_Cambodian-1
S_Irula-2
S_Kapu-1
S_Even-2
S_Kyrgyz-1
S_Thai-1
S_Palestinian-1
B_Yoruba-3
S_BantuTswana-1
S_Mandenka-1
S_Japanese-3
S_Altaian-1
S_Hawaiian-1
S_Kusunda-2
B_Han-3
S_Tujia-1
S_Kinh-2
S_Quechua-2
S_Mixtec-1
S_Zapotec-2
S_She-2
S_Dai-2
S_Kusunda-1
S_Ami-1
S_Han-2
S_Hazara-2
S_Hezhen-1
S_Chukchi-1
S_Atayal-1
S_Yakut-2
S_Hazara-1
S_Relli-1
S_Japanese-1
S_Korean-1
S_Mongola-1
S_Mala-2
S_Tu-1
S_Bengali-1
S_Xibo-2
Ust_Ishim
S_Eskimo_Chaplin-1
S_Eskimo_Sireniki-1
S_Lahu-2
S_North_Ossetian-1
Africa
WestEurasia
CentralAsiaSiberia
SouthAsia
Oceania
EastAsia
America
−0.00050
−0.00025
0.00000
73.0
73.1
73.2
73.3
73.4
73.5

Q

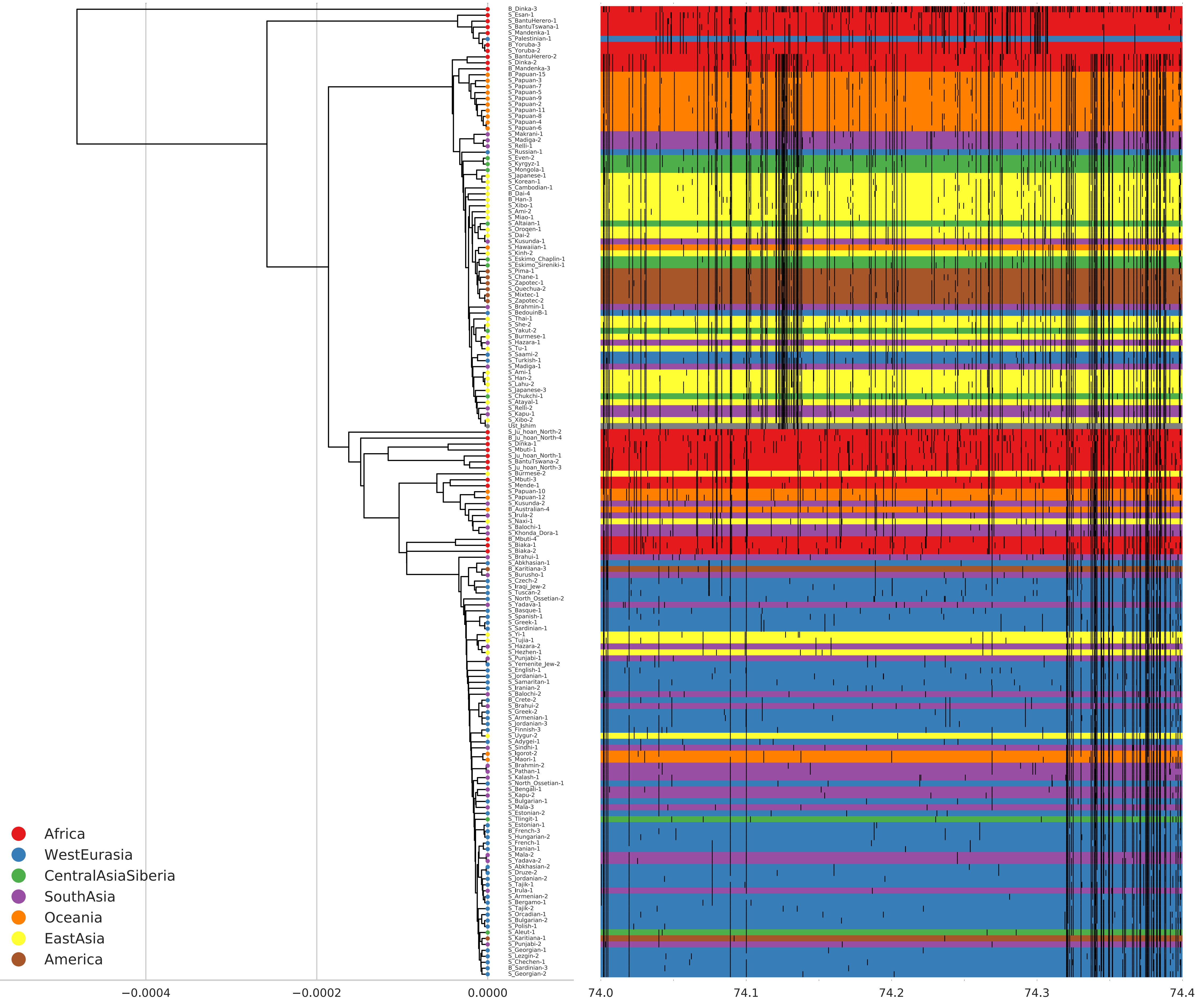
B_Dinka-3
S_Esan-1
S_BantuHerero-1
S_BantuTswana-1
S_Mandenka-1
S_Palestinian-1
B_Yoruba-3
S_Yoruba-2
S_BantuHerero-2
S_Dinka-2
B_Mandenka-3
B_Papuan-15
S_Papuan-3
S_Papuan-7
S_Papuan-5
S_Papuan-9
S_Papuan-2
S_Papuan-11
S_Papuan-8
S_Papuan-4
S_Papuan-6
S_Makrani-1
S_Madiga-2
S_Relli-1
S_Russian-1
S_Even-2
S_Kyrgyz-1
S_Mongola-1
S_Japanese-1
S_Korean-1
S_Cambodian-1
B_Dai-4
B_Han-3
S_Xibo-1
S_Ami-2
S_Miao-1
S_Altaian-1
S_Oroqen-1
S_Dai-2
S_Kusunda-1
S_Hawaiian-1
S_Kinh-2
S_Eskimo_Chaplin-1
S_Eskimo_Sireniki-1
S_Pima-1
S_Chane-1
S_Zapotec-1
S_Quechua-2
S_Mixtec-1
S_Zapotec-2
S_Brahmin-1
S_BedouinB-1
S_Thai-1
S_She-2
S_Yakut-2
S_Burmese-1
S_Hazara-1
S_Tu-1
S_Saami-2
S_Turkish-1
S_Madiga-1
S_Ami-1
S_Han-2
S_Lahu-2
S_Japanese-3
S_Chukchi-1
S_Atayal-1
S_Relli-2
S_Kapu-1
S_Xibo-2
Ust_Ishim
S_Ju_hoan_North-2
B_Ju_hoan_North-4
S_Dinka-1
S_Mbuti-1
S_Ju_hoan_North-1
S_BantuTswana-2
S_Ju_hoan_North-3
S_Burmese-2
S_Mbuti-3
S_Mende-1
S_Papuan-10
S_Papuan-12
S_Kusunda-2
B_Australian-4
S_Irula-2
S_Naxi-1
S_Balochi-1
S_Khonda_Dora-1
B_Mbuti-4
S_Biaka-1
S_Biaka-2
S_Brahui-1
S_Abkhasian-1
B_Karitiana-3
S_Burusho-1
S_Czech-2
S_Iraqi_Jew-2
S_Tuscan-2
S_North_Ossetian-2
S_Yadava-1
S_Basque-1
S_Spanish-1
S_Greek-1
S_Sardinian-1
S_Yi-1
S_Tujia-1
S_Hazara-2
S_Hezhen-1
S_Punjabi-1
S_Yemenite_Jew-2
S_English-1
S_Jordanian-1
S_Samaritan-1
S_Iranian-2
S_Balochi-2
B_Crete-2
S_Brahui-2
S_Greek-2
S_Armenian-1
S_Jordanian-3
S_Finnish-3
S_Uygur-2
S_Adygei-1
S_Sindhi-1
S_Igorot-2
S_Maori-1
S_Brahmin-2
S_Pathan-1
S_Kalash-1
S_North_Ossetian-1
S_Bengali-1
S_Kapu-2
S_Bulgarian-1
S_Mala-3
S_Estonian-2
S_Tlingit-1
S_Estonian-1
B_French-3
S_Hungarian-2
S_French-1
S_Iranian-1
S_Mala-2
S_Yadava-2
S_Abkhasian-2
S_Druze-2
S_Jordanian-2
S_Tajik-1
S_Irula-1
S_Armenian-2
S_Bergamo-1
S_Tajik-2
S_Orcadian-1
S_Bulgarian-2
S_Polish-1
S_Aleut-1
S_Karitiana-1
S_Punjabi-2
S_Georgian-1
S_Lezgin-2
S_Chechen-1
B_Sardinian-3
S_Georgian-2
Africa
WestEurasia
CentralAsiaSiberia
SouthAsia
Oceania
EastAsia
America
−0.0004
−0.0002
0.0000
74.0
74.1
74.2
74.3
74.4

R

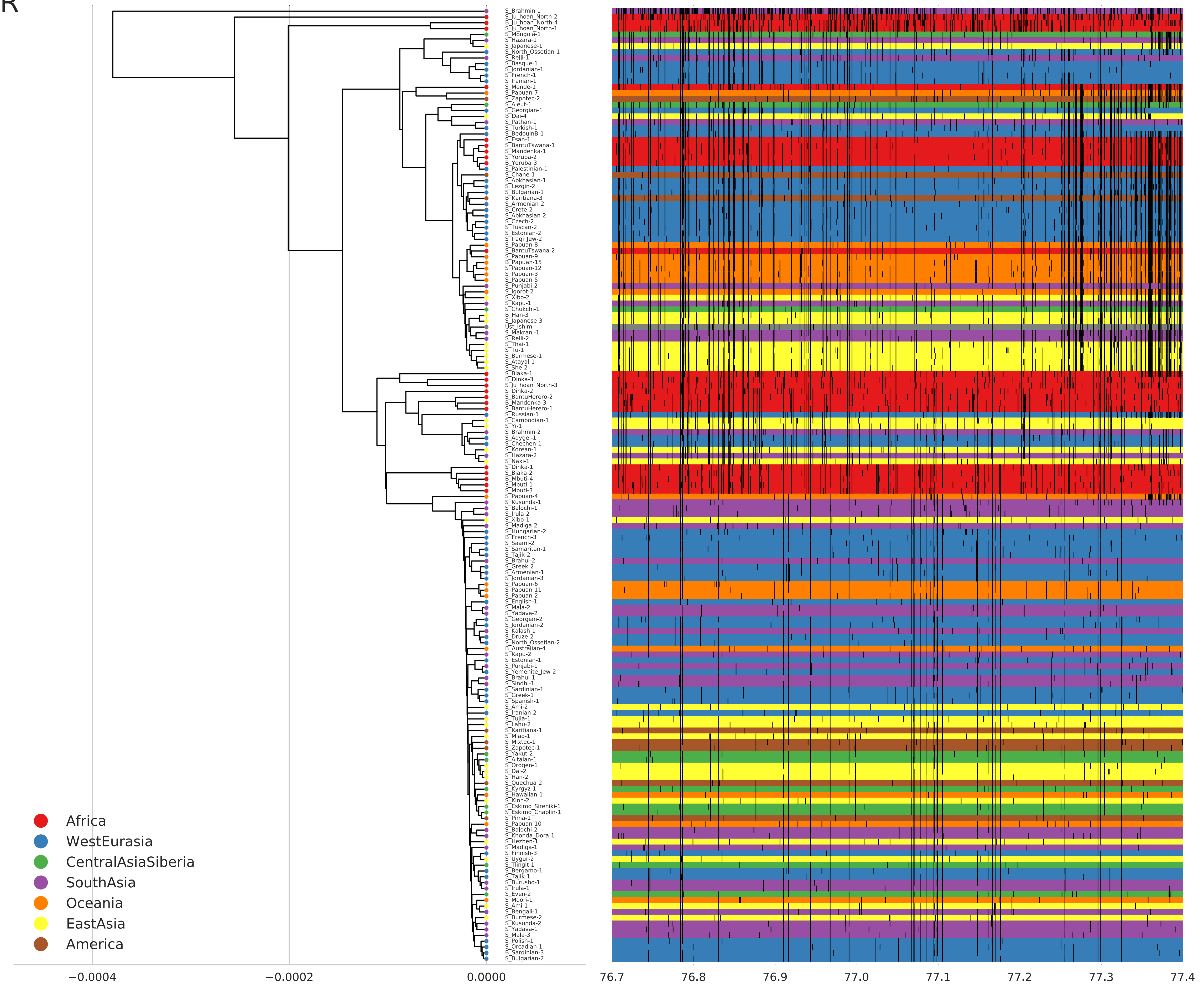
S_Brahmin-1
S_Ju_hoan_North-2
B_Ju_hoan_North-4
S_Ju_hoan_North-1
S_Mongola-1
S_Hazara-1
S_Japanese-1
S_North_Ossetian-1
S_Relli-1
S_Basque-1
S_Jordanian-1
S_French-1
S_Iranian-1
S_Mende-1
S_Papuan-7
S_Zapotec-2
S_Aleut-1
S_Georgian-1
B_Dai-4
S_Pathan-1
S_Turkish-1
S_BedouinB-1
S_Esan-1
S_BantuTswana-1
S_Mandenka-1
S_Yoruba-2
B_Yoruba-3
S_Palestinian-1
S_Chane-1
S_Abkhasian-1
S_Lezgin-2
S_Bulgarian-1
B_Karitiana-3
S_Armenian-2
B_Crete-2
S_Abkhasian-2
S_Czech-2
S_Tuscan-2
S_Estonian-2
S_Iraqi_Jew-2
S_Papuan-8
S_BantuTswana-2
S_Papuan-9
B_Papuan-15
S_Papuan-12
S_Papuan-3
S_Papuan-5
S_Punjabi-2
S_Igorot-2
S_Xibo-2
S_Kapu-1
S_Chukchi-1
B_Han-3
S_Japanese-3
Ust_Ishim
S_Makrani-1
S_Relli-2
S_Thai-1
S_Tu-1
S_Burmese-1
S_Atayal-1
S_She-2
S_Biaka-1
B_Dinka-3
S_Ju_hoan_North-3
S_Dinka-2
S_BantuHerero-2
B_Mandenka-3
S_BantuHerero-1
S_Russian-1
S_Cambodian-1
S_Yi-1
S_Brahmin-2
S_Adygei-1
S_Chechen-1
S_Korean-1
S_Hazara-2
S_Naxi-1
S_Dinka-1
S_Biaka-2
B_Mbuti-4
S_Mbuti-1
S_Mbuti-3
S_Papuan-4
S_Kusunda-1
S_Balochi-1
S_Irula-2
S_Xibo-1
S_Madiga-2
S_Hungarian-2
B_French-3
S_Saami-2
S_Samaritan-1
S_Tajik-2
S_Brahui-2
S_Greek-2
S_Armenian-1
S_Jordanian-3
S_Papuan-6
S_Papuan-11
S_Papuan-2
S_English-1
S_Mala-2
S_Yadava-2
S_Georgian-2
S_Jordanian-2
S_Kalash-1
S_Druze-2
S_North_Ossetian-2
B_Australian-4
S_Kapu-2
S_Estonian-1
S_Punjabi-1
S_Yemenite_Jew-2
S_Brahui-1
S_Sindhi-1
S_Sardinian-1
S_Greek-1
S_Spanish-1
S_Ami-2
S_Iranian-2
S_Tujia-1
S_Lahu-2
S_Karitiana-1
S_Miao-1
S_Mixtec-1
S_Zapotec-1
S_Yakut-2
S_Altaian-1
S_Oroqen-1
S_Dai-2
S_Han-2
S_Quechua-2
S_Kyrgyz-1
S_Hawaiian-1
S_Kinh-2
S_Eskimo_Sireniki-1
S_Eskimo_Chaplin-1
S_Pima-1
S_Papuan-10
S_Balochi-2
S_Khonda_Dora-1
S_Hezhen-1
S_Madiga-1
S_Finnish-3
S_Uygur-2
S_Tlingit-1
S_Bergamo-1
S_Tajik-1
S_Burusho-1
S_Irula-1
S_Even-2
S_Maori-1
S_Ami-1
S_Bengali-1
S_Burmese-2
S_Kusunda-2
S_Yadava-1
S_Mala-3
S_Polish-1
S_Orcadian-1
B_Sardinian-3
S_Bulgarian-2
Africa
WestEurasia
CentralAsiaSiberia
SouthAsia
Oceania
EastAsia
America
−0.0004
−0.0002
0.0000
76.7
76.8
76.9
77.0
77.1
77.2
77.3
77.4

S

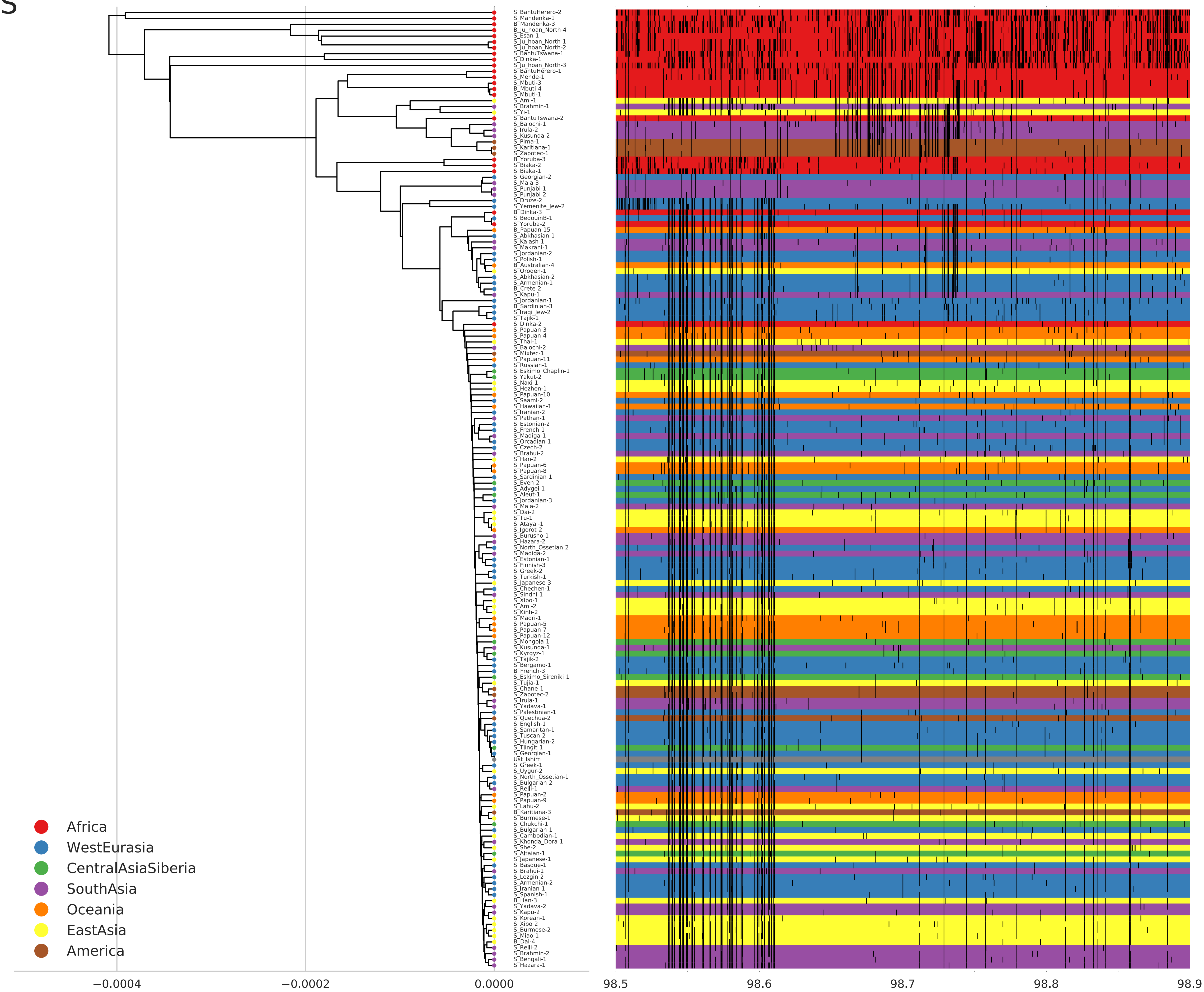
S_BantuHerero-2
S_Mandenka-1
B_Mandenka-3
B_Ju_hoan_North-4
S_Esan-1
S_Ju_hoan_North-1
S_Ju_hoan_North-2
S_BantuTswana-1
S_Dinka-1
S_Ju_hoan_North-3
S_BantuHerero-1
S_Mende-1
S_Mbuti-3
B_Mbuti-4
S_Mbuti-1
S_Ami-1
S_Brahmin-1
S_Yi-1
S_BantuTswana-2
S_Balochi-1
S_Irula-2
S_Kusunda-2
S_Pima-1
S_Karitiana-1
S_Zapotec-1
B_Yoruba-3
S_Biaka-2
S_Biaka-1
S_Georgian-2
S_Mala-3
S_Punjabi-1
S_Punjabi-2
S_Druze-2
S_Yemenite_Jew-2
B_Dinka-3
S_BedouinB-1
S_Yoruba-2
B_Papuan-15
S_Abkhasian-1
S_Kalash-1
S_Makrani-1
S_Jordanian-2
S_Polish-1
B_Australian-4
S_Oroqen-1
S_Abkhasian-2
S_Armenian-1
B_Crete-2
S_Kapu-1
S_Jordanian-1
B_Sardinian-3
S_Iraqi_Jew-2
S_Tajik-1
S_Dinka-2
S_Papuan-3
S_Papuan-4
S_Thai-1
S_Balochi-2
S_Mixtec-1
S_Papuan-11
S_Russian-1
S_Eskimo_Chaplin-1
S_Yakut-2
S_Naxi-1
S_Hezhen-1
S_Papuan-10
S_Saami-2
S_Hawaiian-1
S_Iranian-2
S_Pathan-1
S_Estonian-2
S_French-1
S_Madiga-1
S_Orcadian-1
S_Czech-2
S_Brahui-2
S_Han-2
S_Papuan-6
S_Papuan-8
S_Sardinian-1
S_Even-2
S_Adygei-1
S_Aleut-1
S_Jordanian-3
S_Mala-2
S_Dai-2
S_Tu-1
S_Atayal-1
S_Igorot-2
S_Burusho-1
S_Hazara-2
S_North_Ossetian-2
S_Madiga-2
S_Estonian-1
S_Finnish-3
S_Greek-2
S_Turkish-1
S_Japanese-3
S_Chechen-1
S_Sindhi-1
S_Xibo-1
S_Ami-2
S_Kinh-2
S_Maori-1
S_Papuan-5
S_Papuan-7
S_Papuan-12
S_Mongola-1
S_Kusunda-1
S_Kyrgyz-1
S_Tajik-2
S_Bergamo-1
B_French-3
S_Eskimo_Sireniki-1
S_Tujia-1
S_Chane-1
S_Zapotec-2
S_Irula-1
S_Yadava-1
S_Palestinian-1
S_Quechua-2
S_English-1
S_Samaritan-1
S_Tuscan-2
S_Hungarian-2
S_Tlingit-1
S_Georgian-1
Ust_Ishim
S_Greek-1
S_Uygur-2
S_North_Ossetian-1
S_Bulgarian-2
S_Relli-1
S_Papuan-2
S_Papuan-9
S_Lahu-2
B_Karitiana-3
S_Burmese-1
S_Chukchi-1
S_Bulgarian-1
S_Cambodian-1
S_Khonda_Dora-1
S_She-2
S_Altaian-1
S_Japanese-1
S_Basque-1
S_Brahui-1
S_Lezgin-2
S_Armenian-2
S_Iranian-1
S_Spanish-1
B_Han-3
S_Yadava-2
S_Kapu-2
S_Korean-1
S_Xibo-2
S_Burmese-2
S_Miao-1
B_Dai-4
S_Relli-2
S_Brahmin-2
S_Bengali-1
S_Hazara-1
Africa
WestEurasia
CentralAsiaSiberia
SouthAsia
Oceania
EastAsia
America
−0.0004
−0.0002
0.0000
98.5
98.6
98.7
98.8
98.9

T

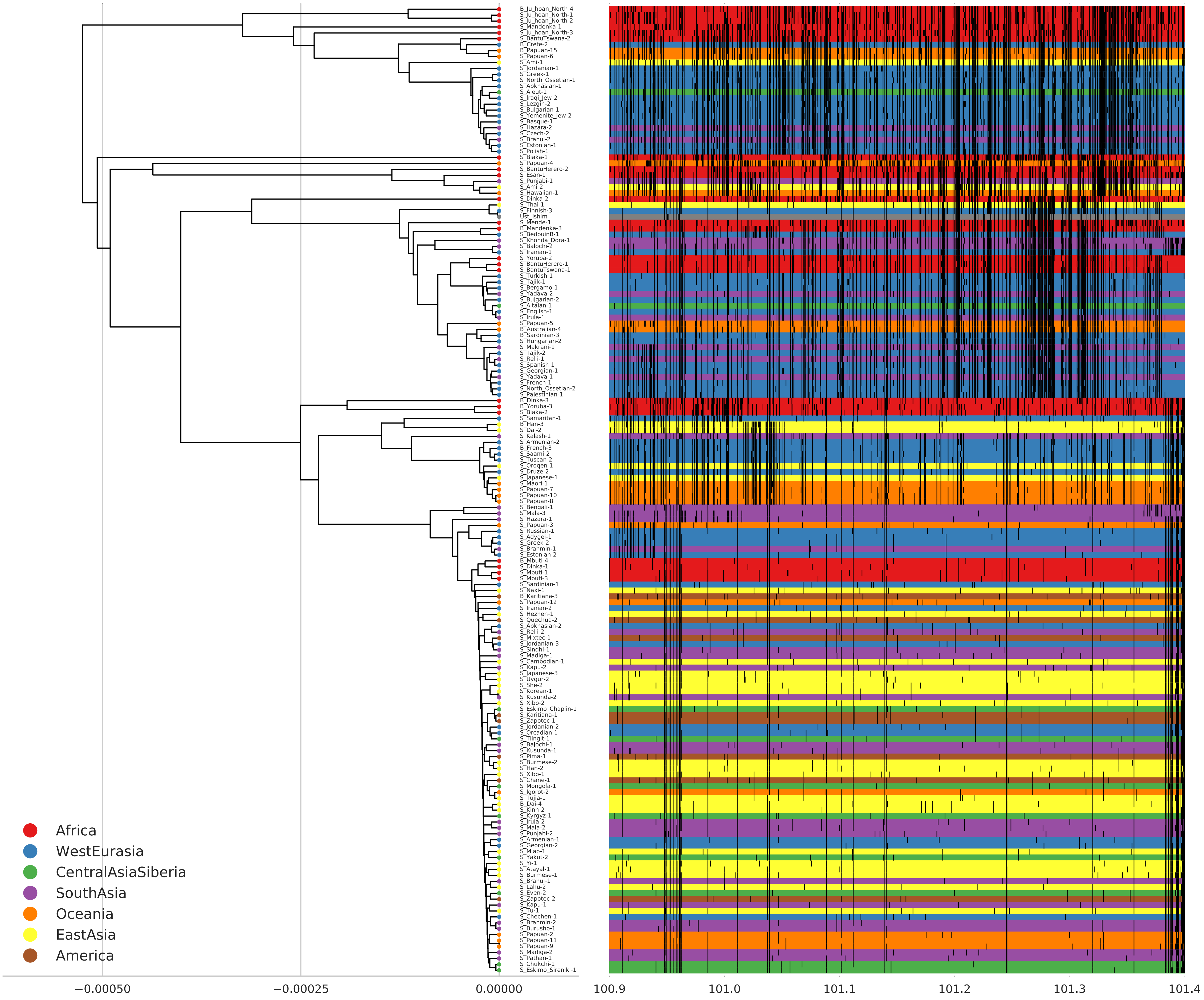

B_Ju_hoan_North-4
S_Ju_hoan_North-1
S_Ju_hoan_North-2
S_Mandenka-1
S_Ju_hoan_North-3
S_BantuTswana-2
B_Crete-2
B_Papuan-15
S_Papuan-6
S_Ami-1
S_Jordanian-1
S_Greek-1
S_North_Ossetian-1
S_Abkhasian-1
S_Aleut-1
S_Iraqi_Jew-2
S_Lezgin-2
S_Bulgarian-1
S_Yemenite_Jew-2
S_Basque-1
S_Hazara-2
S_Czech-2
S_Brahui-2
S_Estonian-1
S_Polish-1
S_Biaka-1
S_Papuan-4
S_BantuHerero-2
S_Esan-1
S_Punjabi-1
S_Ami-2
S_Hawaiian-1
S_Dinka-2
S_Thai-1
S_Finnish-3
Ust_Ishim
S_Mende-1
B_Mandenka-3
S_BedouinB-1
S_Khonda_Dora-1
S_Balochi-2
S_Iranian-1
S_Yoruba-2
S_BantuHerero-1
S_BantuTswana-1
S_Turkish-1
S_Tajik-1
S_Bergamo-1
S_Yadava-2
S_Bulgarian-2
S_Altaian-1
S_English-1
S_Irula-1
S_Papuan-5
B_Australian-4
B_Sardinian-3
S_Hungarian-2
S_Makrani-1
S_Tajik-2
S_Relli-1
S_Spanish-1
S_Georgian-1
S_Yadava-1
S_French-1
S_North_Ossetian-2
S_Palestinian-1
B_Dinka-3
B_Yoruba-3
S_Biaka-2
S_Samaritan-1
B_Han-3
S_Dai-2
S_Kalash-1
S_Armenian-2
B_French-3
S_Saami-2
S_Tuscan-2
S_Oroqen-1
S_Druze-2
S_Japanese-1
S_Maori-1
S_Papuan-7
S_Papuan-10
S_Papuan-8
S_Bengali-1
S_Mala-3
S_Hazara-1
S_Papuan-3
S_Russian-1
S_Adygei-1
S_Greek-2
S_Brahmin-1
S_Estonian-2
B_Mbuti-4
S_Dinka-1
S_Mbuti-1
S_Mbuti-3
S_Sardinian-1
S_Naxi-1
B_Karitiana-3
S_Papuan-12
S_Iranian-2
S_Hezhen-1
S_Quechua-2
S_Abkhasian-2
S_Relli-2
S_Mixtec-1
S_Jordanian-3
S_Sindhi-1
S_Madiga-1
S_Cambodian-1
S_Kapu-2
S_Japanese-3
S_Uygur-2
S_She-2
S_Korean-1
S_Kusunda-2
S_Xibo-2
S_Eskimo_Chaplin-1
S_Karitiana-1
S_Zapotec-1
S_Jordanian-2
S_Orcadian-1
S_Tlingit-1
S_Balochi-1
S_Kusunda-1
S_Pima-1
S_Burmese-2
S_Han-2
S_Xibo-1
S_Chane-1
S_Mongola-1
S_Igorot-2
S_Tujia-1
B_Dai-4
S_Kinh-2
S_Kyrgyz-1
S_Irula-2
S_Mala-2
S_Punjabi-2
S_Armenian-1
S_Georgian-2
S_Miao-1
S_Yakut-2
S_Yi-1
S_Atayal-1
S_Burmese-1
S_Brahui-1
S_Lahu-2
S_Even-2
S_Zapotec-2
S_Kapu-1
S_Tu-1
S_Chechen-1
S_Brahmin-2
S_Burusho-1
S_Papuan-2
S_Papuan-11
S_Papuan-9
S_Madiga-2
S_Pathan-1
S_Chukchi-1
S_Eskimo_Sireniki-1
Africa
WestEurasia
CentralAsiaSiberia
SouthAsia
Oceania
EastAsia
America
−0.00050
−0.00025
0.00000
100.9
101.0
101.1
101.2
101.3
101.4

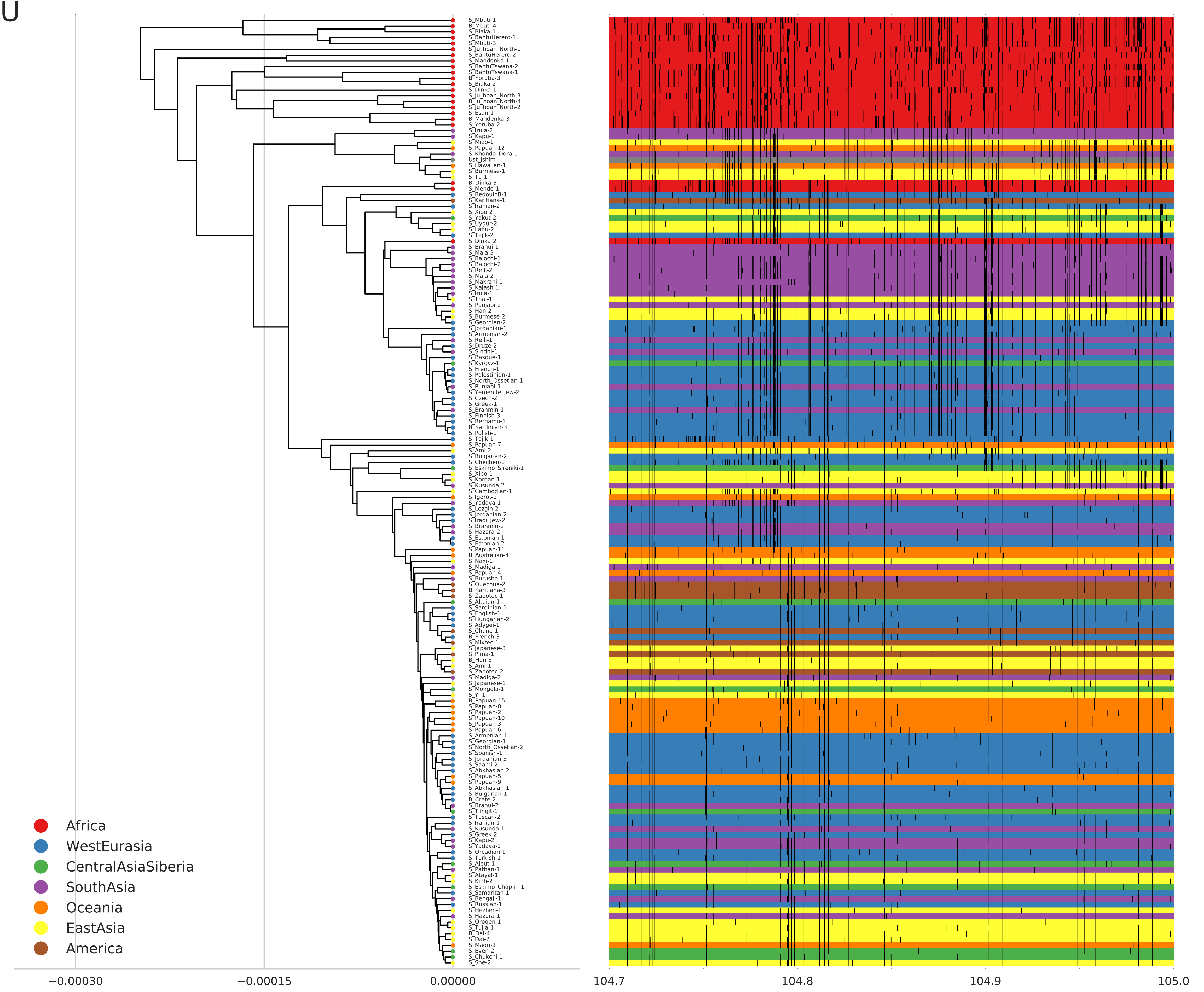
U
S_Mbuti-1
B_Mbuti-4
S_Biaka-1
S_BantuHerero-1
S_Mbuti-3
S_Ju_hoan_North-1
S_BantuHerero-2
S_Mandenka-1
S_BantuTswana-2
S_BantuTswana-1
B_Yoruba-3
S_Biaka-2
S_Dinka-1
S_Ju_hoan_North-3
B_Ju_hoan_North-4
S_Ju_hoan_North-2
S_Esan-1
B_Mandenka-3
S_Yoruba-2
S_Irula-2
S_Kapu-1
S_Miao-1
S_Papuan-12
S_Khonda_Dora-1
Ust_Ishim
S_Hawaiian-1
S_Burmese-1
S_Tu-1
B_Dinka-3
S_Mende-1
S_BedouinB-1
S_Karitiana-1
S_Iranian-2
S_Xibo-2
S_Yakut-2
S_Uygur-2
S_Lahu-2
S_Tajik-2
S_Dinka-2
S_Brahui-1
S_Mala-3
S_Balochi-1
S_Balochi-2
S_Relli-2
S_Mala-2
S_Makrani-1
S_Kalash-1
S_Irula-1
S_Thai-1
S_Punjabi-2
S_Han-2
S_Burmese-2
S_Georgian-2
S_Jordanian-1
S_Armenian-2
S_Relli-1
S_Druze-2
S_Sindhi-1
S_Basque-1
S_Kyrgyz-1
S_French-1
S_Palestinian-1
S_North_Ossetian-1
S_Punjabi-1
S_Yemenite_Jew-2
S_Czech-2
S_Greek-1
S_Brahmin-1
S_Finnish-3
S_Bergamo-1
B_Sardinian-3
S_Polish-1
S_Tajik-1
S_Papuan-7
S_Ami-2
S_Bulgarian-2
S_Chechen-1
S_Eskimo_Sireniki-1
S_Xibo-1
S_Korean-1
S_Kusunda-2
S_Cambodian-1
S_Igorot-2
S_Yadava-1
S_Lezgin-2
S_Jordanian-2
S_Iraqi_Jew-2
S_Brahmin-2
S_Hazara-2
S_Estonian-1
S_Estonian-2
S_Papuan-11
B_Australian-4
S_Naxi-1
S_Madiga-1
S_Papuan-4
S_Burusho-1
S_Quechua-2
B_Karitiana-3
S_Zapotec-1
S_Altaian-1
S_Sardinian-1
S_English-1
S_Hungarian-2
S_Adygei-1
S_Chane-1
B_French-3
S_Mixtec-1
S_Japanese-3
S_Pima-1
B_Han-3
S_Ami-1
S_Zapotec-2
S_Madiga-2
S_Japanese-1
S_Mongola-1
S_Yi-1
B_Papuan-15
S_Papuan-8
S_Papuan-2
S_Papuan-10
S_Papuan-3
S_Papuan-6
S_Armenian-1
S_Georgian-1
S_North_Ossetian-2
S_Spanish-1
S_Jordanian-3
S_Saami-2
S_Abkhasian-2
S_Papuan-5
S_Papuan-9
S_Abkhasian-1
S_Bulgarian-1
B_Crete-2
S_Brahui-2
S_Tlingit-1
S_Tuscan-2
S_Iranian-1
S_Kusunda-1
S_Greek-2
S_Kapu-2
S_Yadava-2
S_Orcadian-1
S_Turkish-1
S_Aleut-1
S_Pathan-1
S_Atayal-1
S_Kinh-2
S_Eskimo_Chaplin-1
S_Samaritan-1
S_Bengali-1
S_Russian-1
S_Hezhen-1
S_Hazara-1
S_Oroqen-1
S_Tujia-1
B_Dai-4
S_Dai-2
S_Maori-1
S_Even-2
S_Chukchi-1
S_She-2
Africa
WestEurasia
CentralAsiaSiberia
SouthAsia
Oceania
EastAsia
America
−0.00030
−0.00015
0.00000
104.7
104.8
104.9
105.0

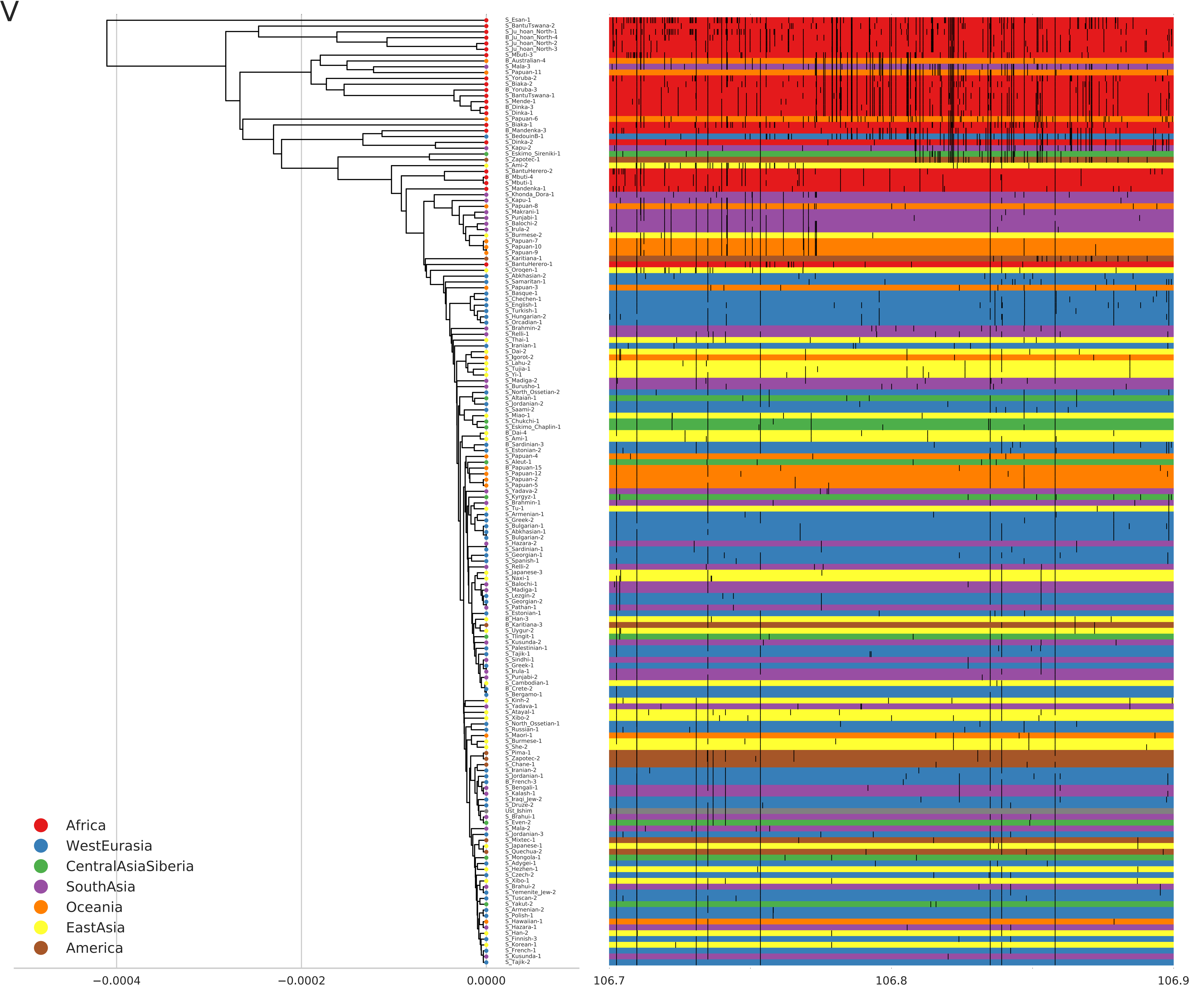
V
Africa
WestEurasia
CentralAsiaSiberia
SouthAsia
Oceania
EastAsia
America
−0.0004
−0.0002
0.0000
106.7
106.8
106.9

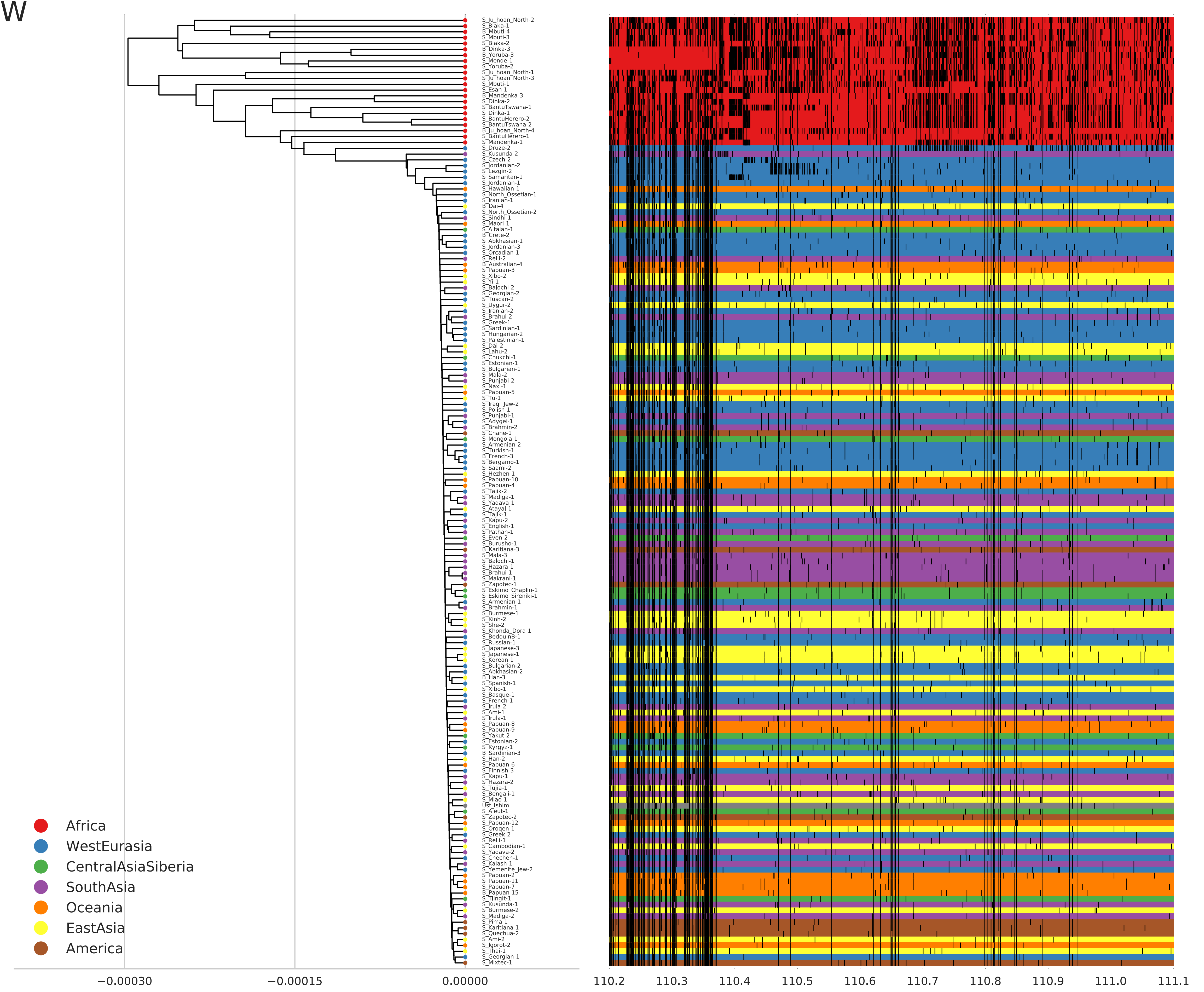
W
Africa
WestEurasia
CentralAsiaSiberia
SouthAsia
Oceania
EastAsia
America
−0.00030
−0.00015
0.00000
110.2
110.3
110.4
110.5
110.6
110.7
110.8
110.9
111.0
111.1

X

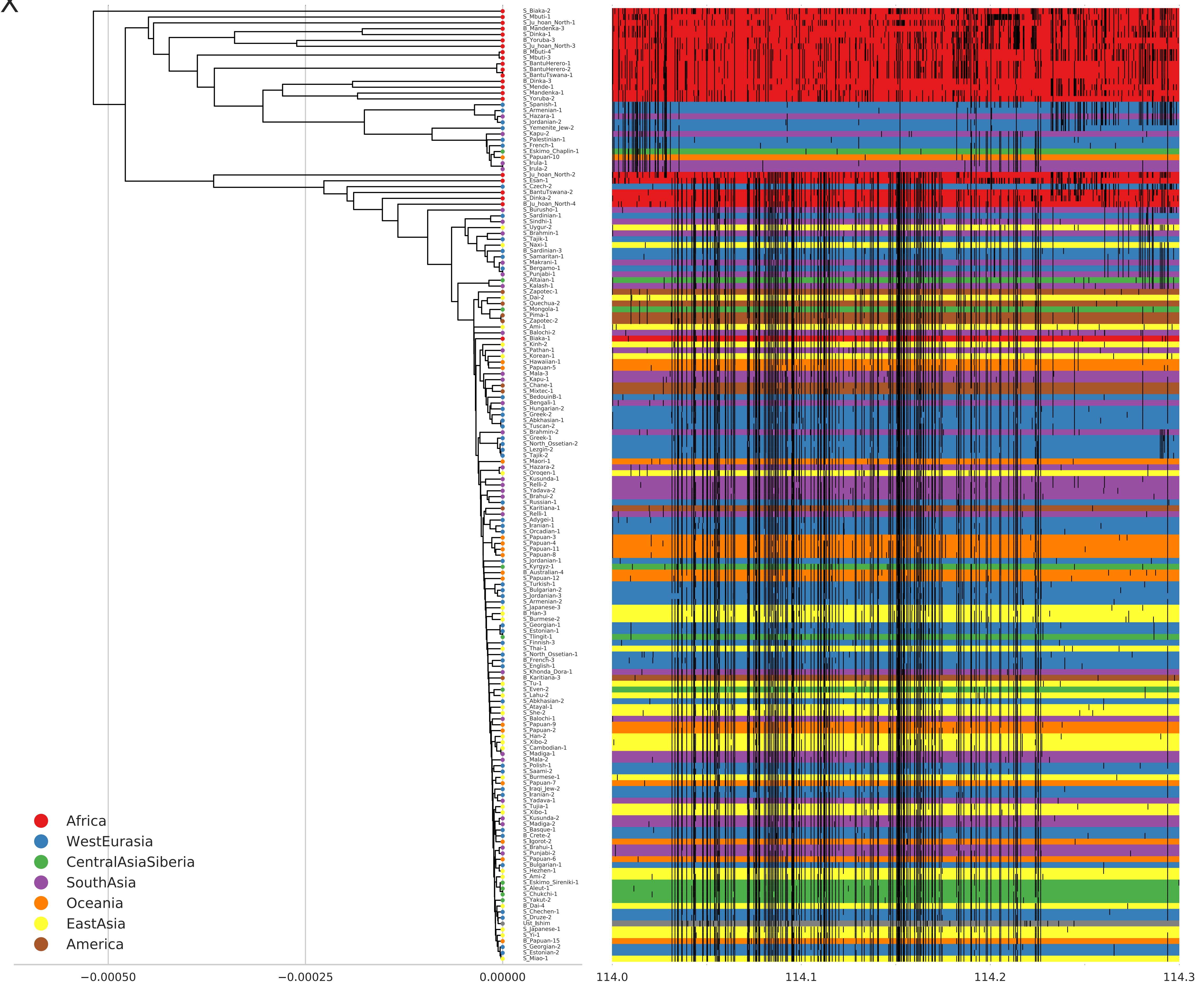
S_Biaka-2
S_Mbuti-1
S_Ju_hoan_North-1
B_Mandenka-3
S_Dinka-1
B_Yoruba-3
S_Ju_hoan_North-3
B_Mbuti-4
S_Mbuti-3
S_BantuHerero-1
S_BantuHerero-2
S_BantuTswana-1
B_Dinka-3
S_Mende-1
S_Mandenka-1
S_Yoruba-2
S_Spanish-1
S_Armenian-1
S_Hazara-1
S_Jordanian-2
S_Yemenite_Jew-2
S_Kapu-2
S_Palestinian-1
S_French-1
S_Eskimo_Chaplin-1
S_Papuan-10
S_Irula-1
S_Irula-2
S_Ju_hoan_North-2
S_Esan-1
S_Czech-2
S_BantuTswana-2
S_Dinka-2
B_Ju_hoan_North-4
S_Burusho-1
S_Sardinian-1
S_Sindhi-1
S_Uygur-2
S_Brahmin-1
S_Tajik-1
S_Naxi-1
B_Sardinian-3
S_Samaritan-1
S_Makrani-1
S_Bergamo-1
S_Punjabi-1
S_Altaian-1
S_Kalash-1
S_Zapotec-1
S_Dai-2
S_Quechua-2
S_Mongola-1
S_Pima-1
S_Zapotec-2
S_Ami-1
S_Balochi-2
S_Biaka-1
S_Kinh-2
S_Pathan-1
S_Korean-1
S_Hawaiian-1
S_Papuan-5
S_Mala-3
S_Kapu-1
S_Chane-1
S_Mixtec-1
S_BedouinB-1
S_Bengali-1
S_Hungarian-2
S_Greek-2
S_Abkhasian-1
S_Tuscan-2
S_Brahmin-2
S_Greek-1
S_North_Ossetian-2
S_Lezgin-2
S_Tajik-2
S_Maori-1
S_Hazara-2
S_Oroqen-1
S_Kusunda-1
S_Relli-2
S_Yadava-2
S_Brahui-2
S_Russian-1
S_Karitiana-1
S_Relli-1
S_Adygei-1
S_Iranian-1
S_Orcadian-1
S_Papuan-3
S_Papuan-4
S_Papuan-11
S_Papuan-8
S_Jordanian-1
S_Kyrgyz-1
B_Australian-4
S_Papuan-12
S_Turkish-1
S_Bulgarian-2
S_Jordanian-3
S_Armenian-2
S_Japanese-3
B_Han-3
S_Burmese-2
S_Georgian-1
S_Estonian-1
S_Tlingit-1
S_Finnish-3
S_Thai-1
S_North_Ossetian-1
B_French-3
S_English-1
S_Khonda_Dora-1
B_Karitiana-3
S_Tu-1
S_Even-2
S_Lahu-2
S_Abkhasian-2
S_Atayal-1
S_She-2
S_Balochi-1
S_Papuan-9
S_Papuan-2
S_Han-2
S_Xibo-2
S_Cambodian-1
S_Madiga-1
S_Mala-2
S_Polish-1
S_Saami-2
S_Burmese-1
S_Papuan-7
S_Iraqi_Jew-2
S_Iranian-2
S_Yadava-1
S_Tujia-1
S_Xibo-1
S_Kusunda-2
S_Madiga-2
S_Basque-1
B_Crete-2
S_Igorot-2
S_Brahui-1
S_Punjabi-2
S_Papuan-6
S_Bulgarian-1
S_Hezhen-1
S_Ami-2
S_Eskimo_Sireniki-1
S_Aleut-1
S_Chukchi-1
S_Yakut-2
B_Dai-4
S_Chechen-1
S_Druze-2
Ust_Ishim
S_Japanese-1
S_Yi-1
B_Papuan-15
S_Georgian-2
S_Estonian-2
S_Miao-1
Africa
WestEurasia
CentralAsiaSiberia
SouthAsia
Oceania
EastAsia
America
−0.00050
−0.00025
0.00000
114.0
114.1
114.2
114.3

Y

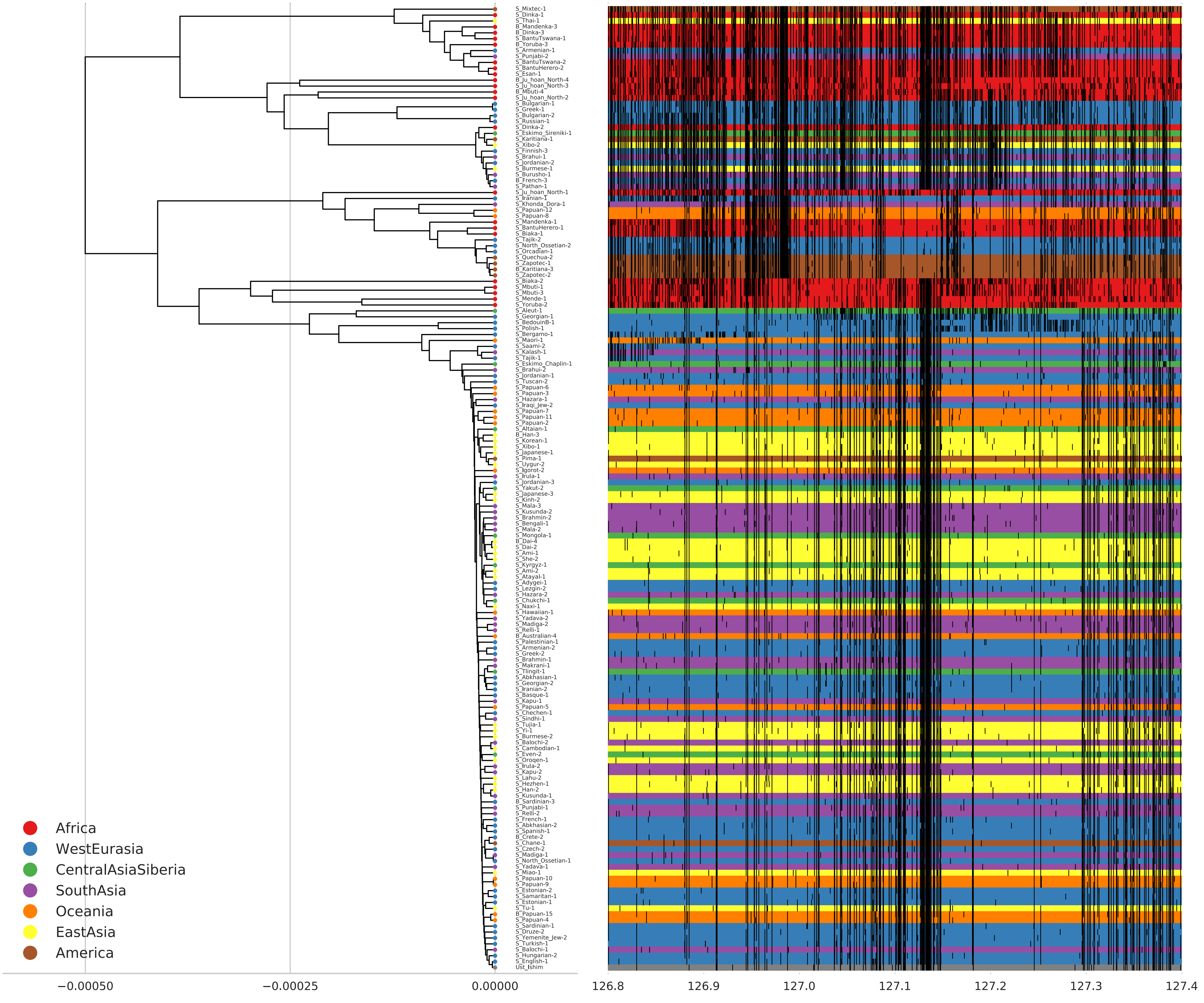

Africa
WestEurasia
CentralAsiaSiberia
SouthAsia
Oceania
EastAsia
America
−0.00050
−0.00025
0.00000
126.8
126.9
127.0
127.1
127.2
127.3
127.4

Z

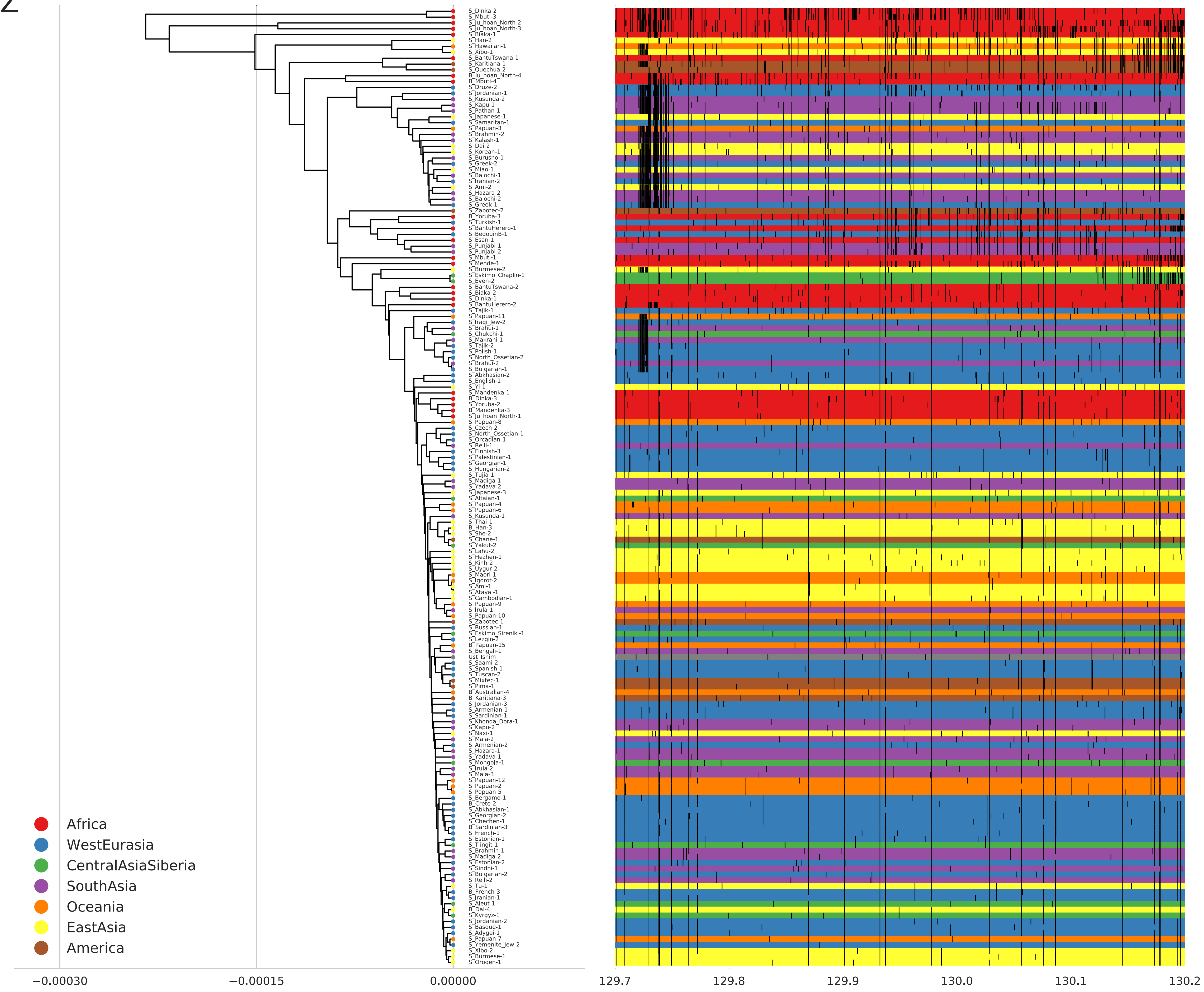
S_Dinka-2
S_Mbuti-3
S_Ju_hoan_North-2
S_Ju_hoan_North-3
S_Biaka-1
S_Han-2
S_Hawaiian-1
S_Xibo-1
S_BantuTswana-1
S_Karitiana-1
S_Quechua-2
B_Ju_hoan_North-4
B_Mbuti-4
S_Druze-2
S_Jordanian-1
S_Kusunda-2
S_Kapu-1
S_Pathan-1
S_Japanese-1
S_Samaritan-1
S_Papuan-3
S_Brahmin-2
S_Kalash-1
S_Dai-2
S_Korean-1
S_Burusho-1
S_Greek-2
S_Miao-1
S_Balochi-1
S_Iranian-2
S_Ami-2
S_Hazara-2
S_Balochi-2
S_Greek-1
S_Zapotec-2
B_Yoruba-3
S_Turkish-1
S_BantuHerero-1
S_BedouinB-1
S_Esan-1
S_Punjabi-1
S_Punjabi-2
S_Mbuti-1
S_Mende-1
S_Burmese-2
S_Eskimo_Chaplin-1
S_Even-2
S_BantuTswana-2
S_Biaka-2
S_Dinka-1
S_BantuHerero-2
S_Tajik-1
S_Papuan-11
S_Iraqi_Jew-2
S_Brahui-1
S_Chukchi-1
S_Makrani-1
S_Tajik-2
S_Polish-1
S_North_Ossetian-2
S_Brahui-2
S_Bulgarian-1
S_Abkhasian-2
S_English-1
S_Yi-1
S_Mandenka-1
B_Dinka-3
S_Yoruba-2
B_Mandenka-3
S_Ju_hoan_North-1
S_Papuan-8
S_Czech-2
S_North_Ossetian-1
S_Orcadian-1
S_Relli-1
S_Finnish-3
S_Palestinian-1
S_Georgian-1
S_Hungarian-2
S_Tujia-1
S_Madiga-1
S_Yadava-2
S_Japanese-3
S_Altaian-1
S_Papuan-4
S_Papuan-6
S_Kusunda-1
S_Thai-1
B_Han-3
S_She-2
S_Chane-1
S_Yakut-2
S_Lahu-2
S_Hezhen-1
S_Kinh-2
S_Uygur-2
S_Maori-1
S_Igorot-2
S_Ami-1
S_Atayal-1
S_Cambodian-1
S_Papuan-9
S_Irula-1
S_Papuan-10
S_Zapotec-1
S_Russian-1
S_Eskimo_Sireniki-1
S_Lezgin-2
B_Papuan-15
S_Bengali-1
Ust_Ishim
S_Saami-2
S_Spanish-1
S_Tuscan-2
S_Mixtec-1
S_Pima-1
B_Australian-4
B_Karitiana-3
S_Jordanian-3
S_Armenian-1
S_Sardinian-1
S_Khonda_Dora-1
S_Kapu-2
S_Naxi-1
S_Mala-2
S_Armenian-2
S_Hazara-1
S_Yadava-1
S_Mongola-1
S_Irula-2
S_Mala-3
S_Papuan-12
S_Papuan-2
S_Papuan-5
S_Bergamo-1
B_Crete-2
S_Abkhasian-1
S_Georgian-2
S_Chechen-1
B_Sardinian-3
S_French-1
S_Estonian-1
S_Tlingit-1
S_Brahmin-1
S_Madiga-2
S_Estonian-2
S_Sindhi-1
S_Bulgarian-2
S_Relli-2
S_Tu-1
B_French-3
S_Iranian-1
S_Aleut-1
B_Dai-4
S_Kyrgyz-1
S_Jordanian-2
S_Basque-1
S_Adygei-1
S_Papuan-7
S_Yemenite_Jew-2
S_Xibo-2
S_Burmese-1
S_Oroqen-1
Africa
WestEurasia
CentralAsiaSiberia
SouthAsia
Oceania
EastAsia
America
−0.00030
−0.00015
0.00000
129.7
129.8
129.9
130.0
130.1
130.2

$

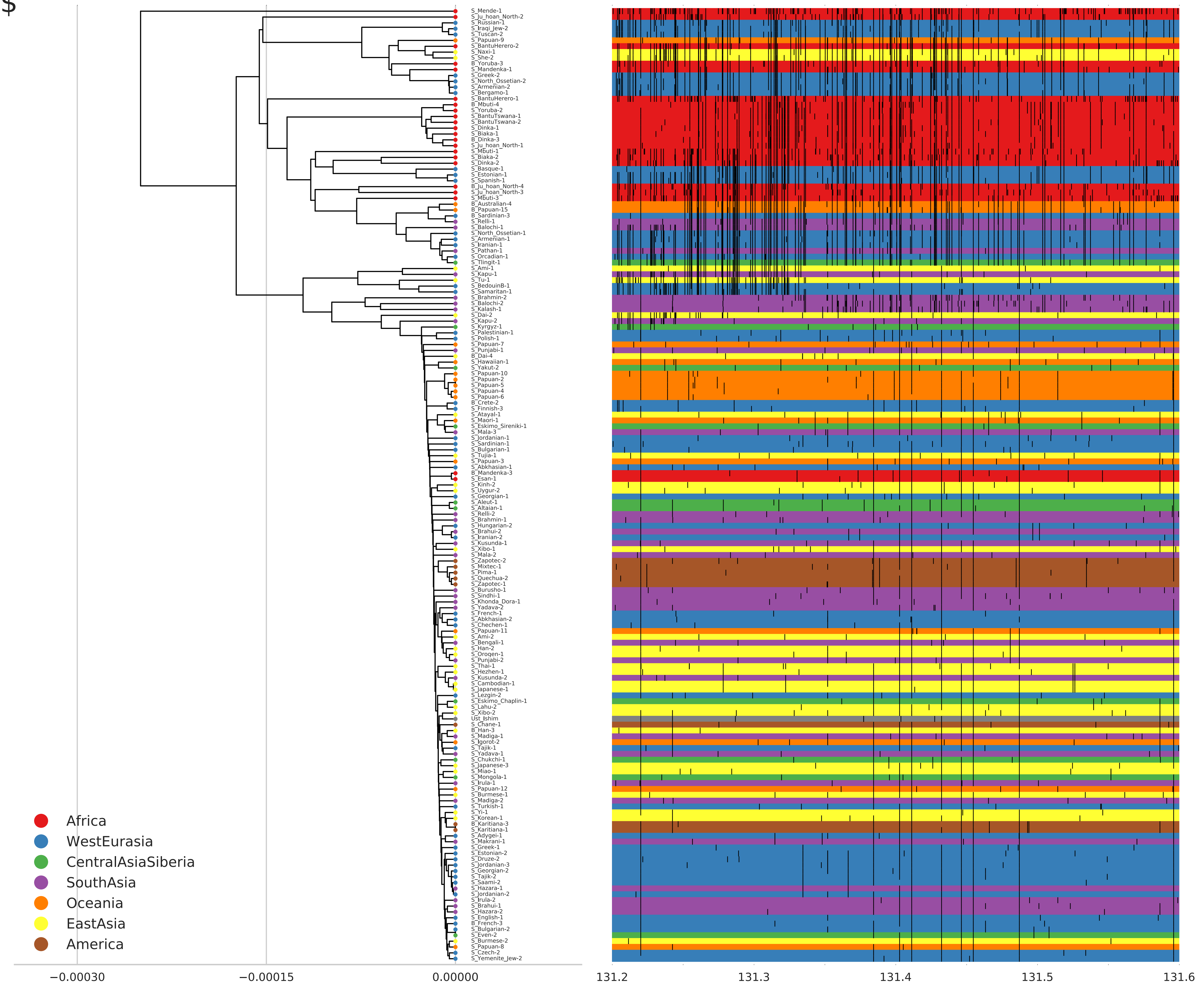
S_Mende-1
S_Ju_hoan_North-2
S_Russian-1
S_Iraqi_Jew-2
S_Tuscan-2
S_Papuan-9
S_BantuHerero-2
S_Naxi-1
S_She-2
B_Yoruba-3
S_Mandenka-1
S_Greek-2
S_North_Ossetian-2
S_Armenian-2
S_Bergamo-1
S_BantuHerero-1
B_Mbuti-4
S_Yoruba-2
S_BantuTswana-1
S_BantuTswana-2
S_Dinka-1
S_Biaka-1
B_Dinka-3
S_Ju_hoan_North-1
S_Mbuti-1
S_Biaka-2
S_Dinka-2
S_Basque-1
S_Estonian-1
S_Spanish-1
B_Ju_hoan_North-4
S_Ju_hoan_North-3
S_Mbuti-3
B_Australian-4
B_Papuan-15
B_Sardinian-3
S_Relli-1
S_Balochi-1
S_North_Ossetian-1
S_Armenian-1
S_Iranian-1
S_Pathan-1
S_Orcadian-1
S_Tlingit-1
S_Ami-1
S_Kapu-1
S_Tu-1
S_BedouinB-1
S_Samaritan-1
S_Brahmin-2
S_Balochi-2
S_Kalash-1
S_Dai-2
S_Kapu-2
S_Kyrgyz-1
S_Palestinian-1
S_Polish-1
S_Papuan-7
S_Punjabi-1
B_Dai-4
S_Hawaiian-1
S_Yakut-2
S_Papuan-10
S_Papuan-2
S_Papuan-5
S_Papuan-4
S_Papuan-6
B_Crete-2
S_Finnish-3
S_Atayal-1
S_Maori-1
S_Eskimo_Sireniki-1
S_Mala-3
S_Jordanian-1
S_Sardinian-1
S_Bulgarian-1
S_Tujia-1
S_Papuan-3
S_Abkhasian-1
B_Mandenka-3
S_Esan-1
S_Kinh-2
S_Uygur-2
S_Georgian-1
S_Aleut-1
S_Altaian-1
S_Relli-2
S_Brahmin-1
S_Hungarian-2
S_Brahui-2
S_Iranian-2
S_Kusunda-1
S_Xibo-1
S_Mala-2
S_Zapotec-2
S_Mixtec-1
S_Pima-1
S_Quechua-2
S_Zapotec-1
S_Burusho-1
S_Sindhi-1
S_Khonda_Dora-1
S_Yadava-2
S_French-1
S_Abkhasian-2
S_Chechen-1
S_Papuan-11
S_Ami-2
S_Bengali-1
S_Han-2
S_Oroqen-1
S_Punjabi-2
S_Thai-1
S_Hezhen-1
S_Kusunda-2
S_Cambodian-1
S_Japanese-1
S_Lezgin-2
S_Eskimo_Chaplin-1
S_Lahu-2
S_Xibo-2
Ust_Ishim
S_Chane-1
B_Han-3
S_Madiga-1
S_Igorot-2
S_Tajik-1
S_Yadava-1
S_Chukchi-1
S_Japanese-3
S_Miao-1
S_Mongola-1
S_Irula-1
S_Papuan-12
S_Burmese-1
S_Madiga-2
S_Turkish-1
S_Yi-1
S_Korean-1
B_Karitiana-3
S_Karitiana-1
S_Adygei-1
S_Makrani-1
S_Greek-1
S_Estonian-2
S_Druze-2
S_Jordanian-3
S_Georgian-2
S_Tajik-2
S_Saami-2
S_Hazara-1
S_Jordanian-2
S_Irula-2
S_Brahui-1
S_Hazara-2
S_English-1
B_French-3
S_Bulgarian-2
S_Even-2
S_Burmese-2
S_Papuan-8
S_Czech-2
S_Yemenite_Jew-2
Africa
WestEurasia
CentralAsiaSiberia
SouthAsia
Oceania
EastAsia
America
−0.00030
−0.00015
0.00000
131.2
131.3
131.4
131.5
131.6

%

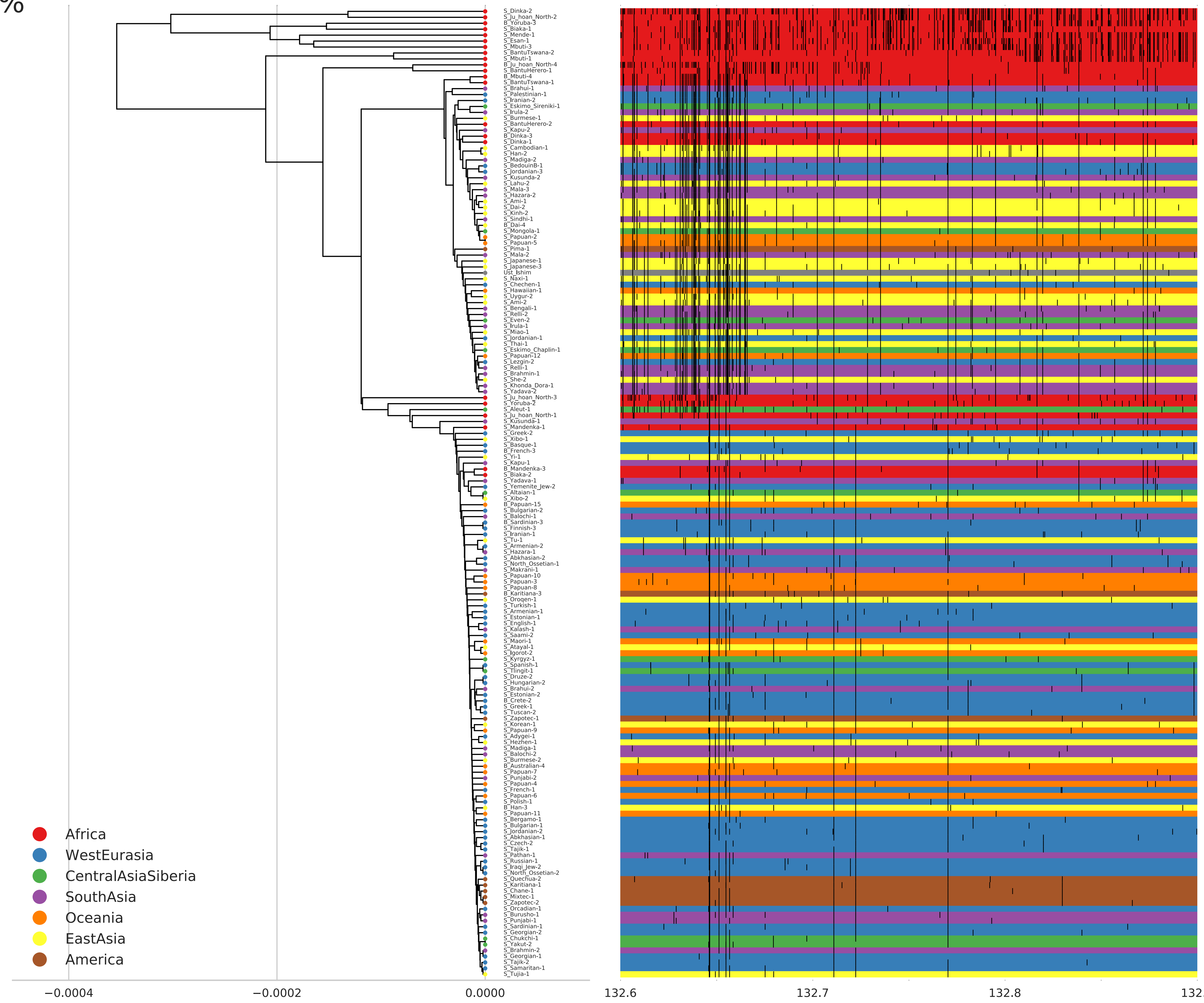

Africa
WestEurasia
CentralAsiaSiberia
SouthAsia
Oceania
EastAsia
America
−0.0004
−0.0002
0.0000
132.6
132.7
132.8
132.9

&

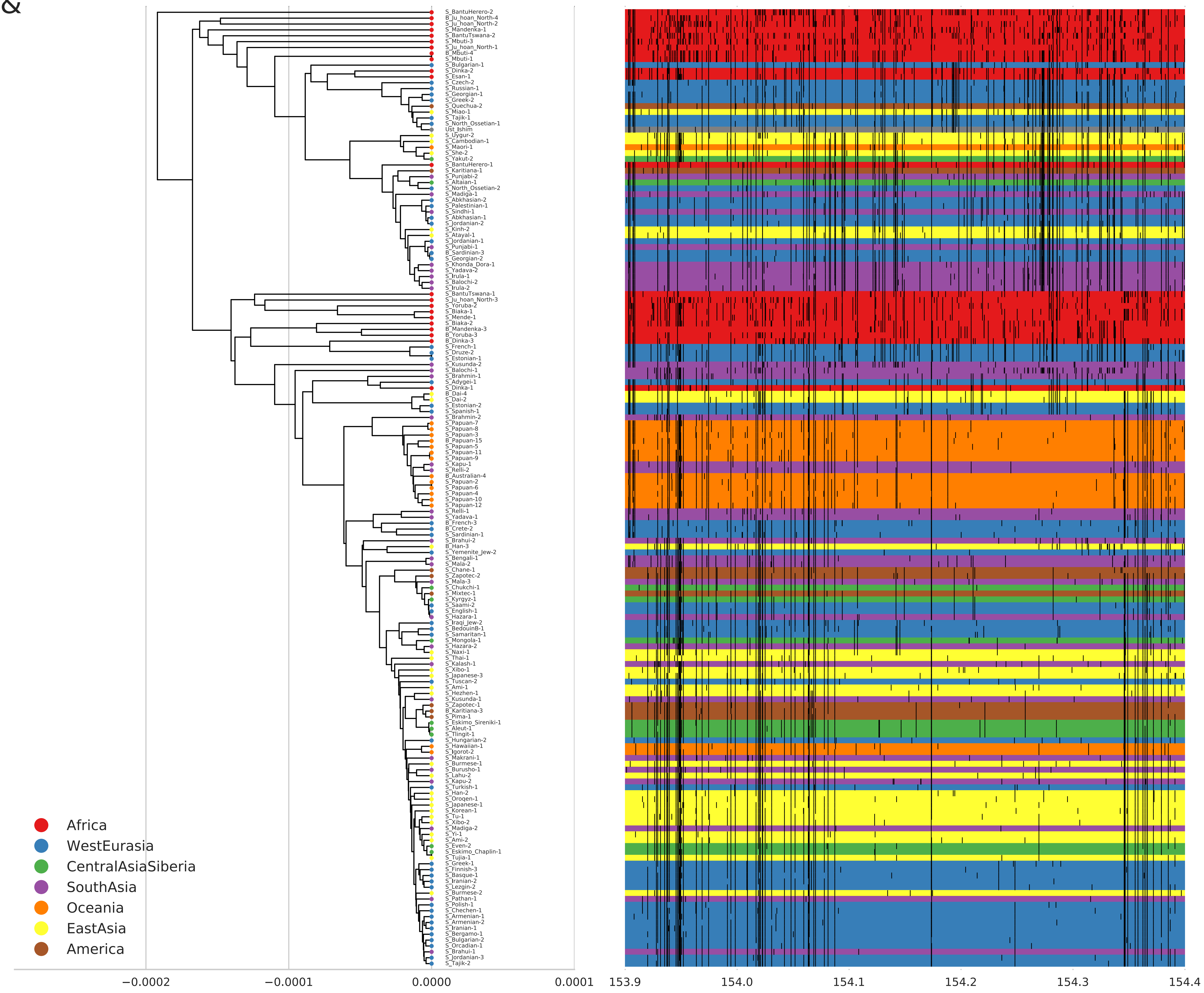
S_BantuHerero-2
B_Ju_hoan_North-4
S_Ju_hoan_North-2
S_Mandenka-1
S_BantuTswana-2
S_Mbuti-3
S_Ju_hoan_North-1
B_Mbuti-4
S_Mbuti-1
S_Bulgarian-1
S_Dinka-2
S_Esan-1
S_Czech-2
S_Russian-1
S_Georgian-1
S_Greek-2
S_Quechua-2
S_Miao-1
S_Tajik-1
S_North_Ossetian-1
Ust_Ishim
S_Uygur-2
S_Cambodian-1
S_Maori-1
S_She-2
S_Yakut-2
S_BantuHerero-1
S_Karitiana-1
S_Punjabi-2
S_Altaian-1
S_North_Ossetian-2
S_Madiga-1
S_Abkhasian-2
S_Palestinian-1
S_Sindhi-1
S_Abkhasian-1
S_Jordanian-2
S_Kinh-2
S_Atayal-1
S_Jordanian-1
S_Punjabi-1
B_Sardinian-3
S_Georgian-2
S_Khonda_Dora-1
S_Yadava-2
S_Irula-1
S_Balochi-2
S_Irula-2
S_BantuTswana-1
S_Ju_hoan_North-3
S_Yoruba-2
S_Biaka-1
S_Mende-1
S_Biaka-2
B_Mandenka-3
B_Yoruba-3
B_Dinka-3
S_French-1
S_Druze-2
S_Estonian-1
S_Kusunda-2
S_Balochi-1
S_Brahmin-1
S_Adygei-1
S_Dinka-1
B_Dai-4
S_Dai-2
S_Estonian-2
S_Spanish-1
S_Brahmin-2
S_Papuan-7
S_Papuan-8
S_Papuan-3
B_Papuan-15
S_Papuan-5
S_Papuan-11
S_Papuan-9
S_Kapu-1
S_Relli-2
B_Australian-4
S_Papuan-2
S_Papuan-6
S_Papuan-4
S_Papuan-10
S_Papuan-12
S_Relli-1
S_Yadava-1
B_French-3
B_Crete-2
S_Sardinian-1
S_Brahui-2
B_Han-3
S_Yemenite_Jew-2
S_Bengali-1
S_Mala-2
S_Chane-1
S_Zapotec-2
S_Mala-3
S_Chukchi-1
S_Mixtec-1
S_Kyrgyz-1
S_Saami-2
S_English-1
S_Hazara-1
S_Iraqi_Jew-2
S_BedouinB-1
S_Samaritan-1
S_Mongola-1
S_Hazara-2
S_Naxi-1
S_Thai-1
S_Kalash-1
S_Xibo-1
S_Japanese-3
S_Tuscan-2
S_Ami-1
S_Hezhen-1
S_Kusunda-1
S_Zapotec-1
B_Karitiana-3
S_Pima-1
S_Eskimo_Sireniki-1
S_Aleut-1
S_Tlingit-1
S_Hungarian-2
S_Hawaiian-1
S_Igorot-2
S_Makrani-1
S_Burmese-1
S_Burusho-1
S_Lahu-2
S_Kapu-2
S_Turkish-1
S_Han-2
S_Oroqen-1
S_Japanese-1
S_Korean-1
S_Tu-1
S_Xibo-2
S_Madiga-2
S_Yi-1
S_Ami-2
S_Even-2
S_Eskimo_Chaplin-1
S_Tujia-1
S_Greek-1
S_Finnish-3
S_Basque-1
S_Iranian-2
S_Lezgin-2
S_Burmese-2
S_Pathan-1
S_Polish-1
S_Chechen-1
S_Armenian-1
S_Armenian-2
S_Iranian-1
S_Bergamo-1
S_Bulgarian-2
S_Orcadian-1
S_Brahui-1
S_Jordanian-3
S_Tajik-2
Africa
WestEurasia
CentralAsiaSiberia
SouthAsia
Oceania
EastAsia
America
−0.0002
−0.0001
0.0000
0.0001
153.9
154.0
154.1
154.2
154.3
154.4
